# A journey towards developing a new cleavable crosslinker reagent for in-cell crosslinking

**DOI:** 10.1101/2024.11.05.621843

**Authors:** Fränze Müller, Bogdan Razvan Brutiu, Iakovos Saridakis, Thomas Leischner, Micha Birklbauer, Manuel Matzinger, Mathias Madalinski, Thomas Lendl, Saad Shaaban, Viktoria Dorfer, Nuno Maulide, Karl Mechtler

## Abstract

Crosslinking mass spectrometry (XL-MS) is a powerful technology that recently emerged as an essential complementary tool for elucidating protein structures and mapping interactions within a protein network. Crosslinkers which are amenable to post-linking backbone cleavage simplify peptide identification, aid in 3D structure determination and enable system-wide studies of protein-protein interactions (PPIs) in cellular environments. However, state-of-the-art cleavable linkers are fraught with practical limitations, including extensive evaluation of fragmentation energies and fragmentation behaviour of the crosslinker backbone. We herein introduce DiSPASO as a lysine-selective, MS-cleavable cross-linker with an alkyne handle for affinity enrichment. DiSPASO was designed and developed for efficient cell membrane permeability and crosslinking while securing low cellular perturbation. We tested DiSPASO employing three different copper-based enrichment strategies using model systems with increasing complexity (Cas9-Halo, purified ribosomes, live cells). Fluorescence microscopy in-cell crosslinking experiments revealed a rapid uptake of DiSPASO into HEK 293 cells within 5 minutes. While DiSPASO represents progress in cellular PPI analysis, its limitations and low crosslinking yield in cellular environments require careful optimisation of the crosslinker design, highlighting the complexity of developing effective XL-MS tools and the importance of continuous innovation in accurately mapping PPI networks within dynamic cellular environments.

**Graphical Abstract:** 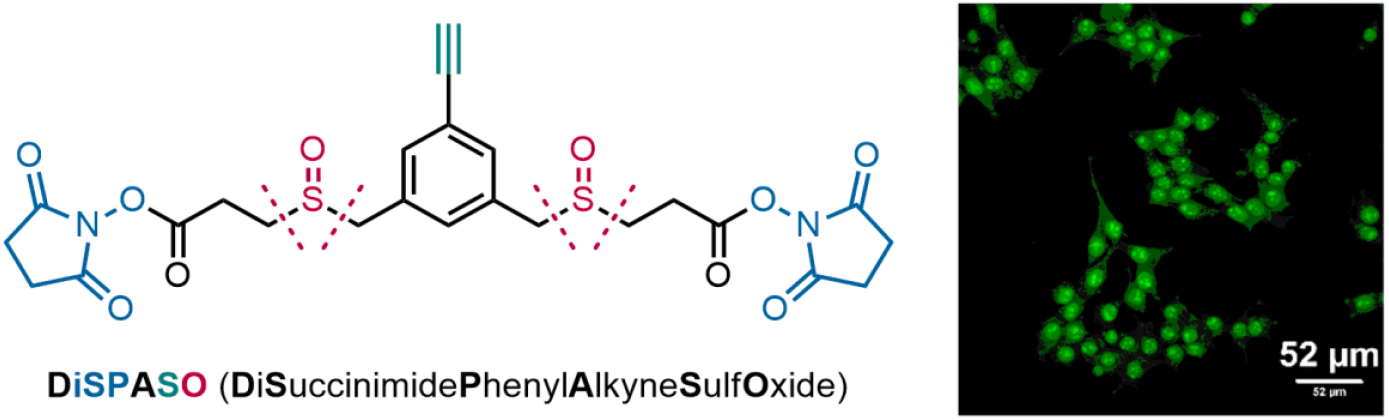

## Introduction

Crosslinking mass spectrometry (XL-MS) has seen significant advancements over the past few decades, evolving into a powerful complementary technique for studying the spatial arrangement of proteins and their interactions within complexes^1–5^. XL-MS is particularly useful for studying protein structures and interactions in their native environments, including cells, organelles, and tissues^1,2,6^ and has therefore significantly enhanced structural analysis by offering precise distance constraints, intrinsic to their chemical properties^7^. This capability is particularly valuable for resolving the three-dimensional architecture of proteins and protein complexes, especially when combined with complementary techniques such as HDX-MS or cryo-EM^3,7–11^.

MS-cleavable crosslinkers, a class of crosslinkers amenable to backbone fragmentation upon MS acquisition, have gained attention in XL-MS by greatly simplifying the MS analysis and facilitating the identification of crosslinked peptides^12–17^. The data analysis process is supported by the generation of characteristic doublets of fragment ions during MS2 fragmentation, eventually enabling straightforward and accurate identification of crosslinks even in complex mixtures. Moreover, cleavable crosslinkers have enabled system-wide XL-MS studies, capturing protein-protein interactions (PPIs) in cellular environments, including weak or transient interactions^18–21^. This advancement allows for the exploration of the structural dynamics of proteins in their native state. Despite advancements in data analysis, there remain challenges in accurately determining false discovery rates (FDR) for identified PPIs. Furthermore, the complexity of the data requires careful handling of error estimation to ensure the reliability of reported PPIs^22,23^. Additionally, there are practical limitations such as the requirement for specific conditions for cleavage, the potential for incomplete cleavage, and the need for specialized mass spectrometry setups to detect and analyze cleavage products effectively^24^. Hence, crosslinkers are often combined with enrichable groups, such as biotin^25– 29^, known for its high affinity towards streptavidin or avidin, and sometimes with MS-cleavable groups, for proteome-wide analyses^30–32^. However, the strong streptavidin-biotin interaction can raise difficulties upon release from the solid support, prompting alternative strategies like solid-supported monomeric avidin or incorporating release groups in the crosslinker backbone^28,33,34^. Various release groups, including PEG^25^, pinacol esters^25,30^, azobenzenes^27^, disulfides^27,35^, and photocleavable groups^27^, have been explored. Click chemistry groups offer versatility, serving as capture or enrichment groups, with reversible options like acid-cleavable acetal or disulfide bonds^35–41^. While this innovation marks significant progress, it also presents opportunities for further refinement, as demonstrated by the development of a negatively charged phosphonic acid as a crosslinker reagent^42^. While this offers simplicity, its negative charge inhibits cell membrane crossing, hindering in-cell crosslinking. This hurdle could be overcome by protecting the negative charge with a chemical protection group^43^. Chemical modifications in crosslinking reagents offer promising functionalities but also pose challenges like increased hydrophobicity. Cleavable crosslinkers have advanced XL-MS, enhancing protein structure analysis, though further improvements are needed to overcome current limitations.

To overcome the aforementioned limitations, we hereby introduce DiSPASO (**1**) as a novel lysine reactive, MS-cleavable and membrane-permeable crosslinker featuring an alkyne-based click chemistry handle for affinity enrichment. This study provides a detailed exploration of the design, synthesis, and careful characterization of DiSPASO, providing a comprehensive evaluation of its chemical properties, effectiveness in mapping protein-protein interactions, and performance relative to existing crosslinkers. This analysis will also explore DiSPASO’s fragmentation behaviour in mass spectrometry, highlighting both its advantages and the challenges encountered, thereby offering insights into its potential and areas for further optimization in crosslinking mass spectrometry applications.

## Results

### Synthesis of a novel cleavable and enrichable crosslinker reagent DiSPASO and its modular perspective

We designed the crosslinker DiSPASO (**1**) bearing 4 key elements: a stable underexplored core decorated with an enrichment site (terminal alkyne), two side-chains containing the cleavable sulfoxide moieties and terminal NHS-esters for targeting lysins. A retrosynthetic analysis of compound (**1**) suggested a common intermediate **S5** in the synthesis pathway. **S5** was efficiently synthesized from commercially available diester amine **S1** through a series of reactions: a Sandmeyer reaction to introduce the aryl bromide, reduction of the ester groups, an Appel-type bromination and finally substitution with the corresponding methyl ester-bearing thiol. **S5** was obtained on a gram scale. The aryl bromide moiety presents an ideal candidate for cross-coupling with various enrichment sites. In this study, we chose a Sonogashira coupling. A global deprotection to the diacid, followed up by NHS coupling formed precursor **S8a**. Optimized oxidation delivered the novel enrichable cross-linker DiSPASO (**1**) in over 150 mg (**Figure 1A**). With a streamlined synthesis in hand, the NHP derivative (**2**) was also obtained after two simple steps (**Figure 1B**). The high-yielding steps can easily be scaled up.

**Figure 1:**
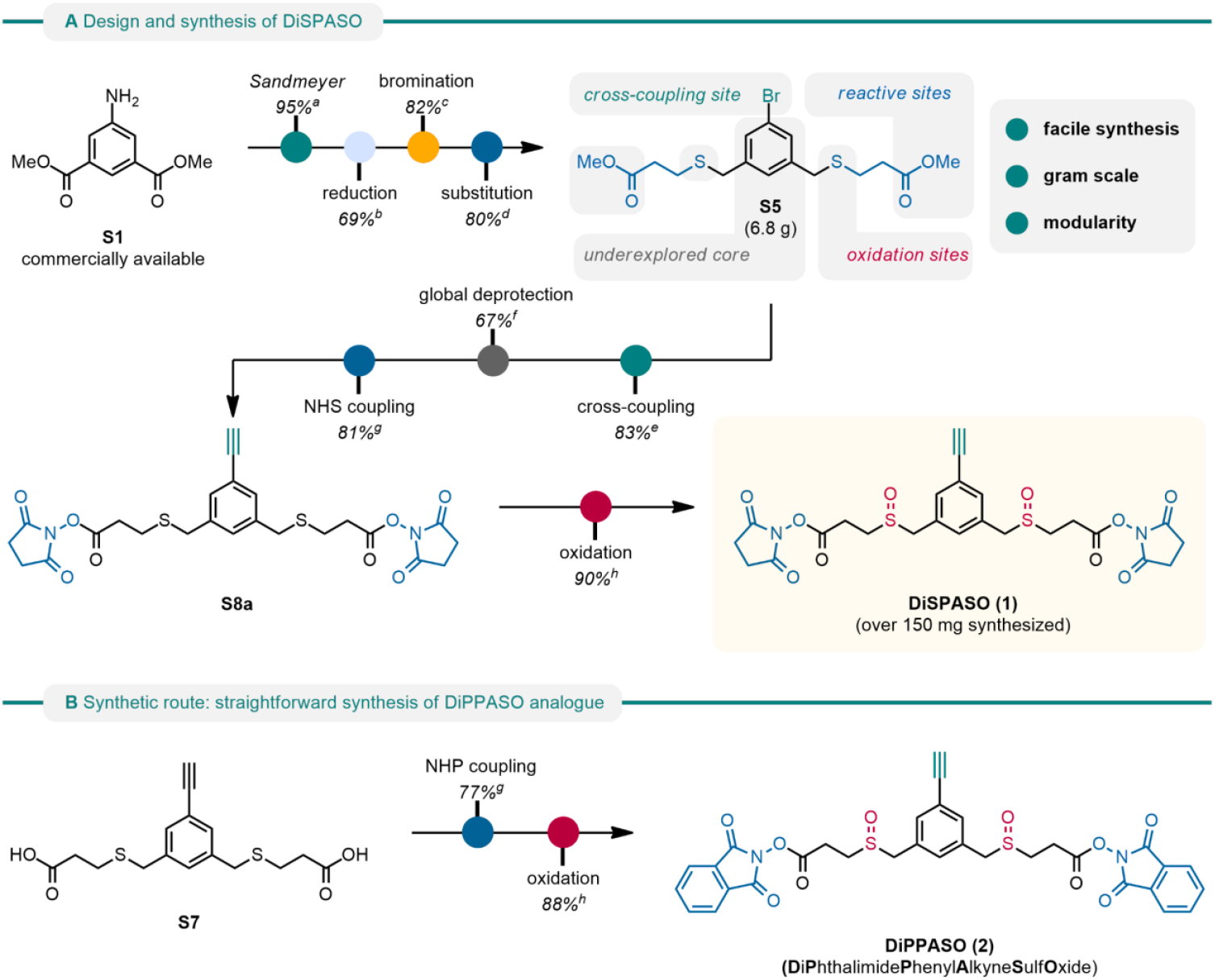
(A) Design and synthesis of DiSPASO. (B) Synthetic route: straightforward synthesis of DiPPASO. ^a^Sandmeyer Reaction: 1.2 eq. NaNO^2^/HBr, 0 °C, 1h, 1.4 eq. CuBr, 0 °C, 18 h. ^b^Reduction: 2.0 eq. LiAlH4, THF, 0 °C, 2 h. ^c^Bromination: 8.5 eq. PBr3, DCM, 40 °C, 15 h, then H^2^O, 23 °C, 20 h. ^d^Substitution: 2.05 eq. methyl 3-mercaptopropanoate, 2.05 eq. K^2^CO3, DMF, 40 °C, 30 h. ^e^Cross-coupling: 1.0 eq. TMS-acetylene, 0.05 eq. Pd(dppf)Cl^2^·CH^2^Cl^2^, 0.05 eq. CuI, THF, 60 °C, 20 h. ^f^Global deprotection: 6.0 eq. LiOH, THF/H^2^O (3:1), 23 °C, 20 h. ^g^NHS/NHP coupling: 2.2 eq. NHS or NHP, 2.2 eq. EDCI•HCl, 0.2 eq. NEt3, DMF, 23 °C, 24 h. ^h^Oxidation: 2.0 eq. mCPBA, DCM, 0 °C, 2 h.

### DiSPASO in context to other in-cell and lysate crosslinkers

DiSPASO was engineered as an in-cell crosslinker reagent with good cell membrane permeability properties while maintaining the solubility of the reagent during the in-cell crosslinking procedure. For a direct comparison of DiSPASO in the context of other state-of-the-art crosslinkers used in in-cell or cell lysate crosslinking, we used a topological polar surface area (tPSA) against partition coefficient (cLogP) plot (**Figure 2**). This visualization offers critical insights into the chemical properties of compounds, essential for design and optimization purposes. It effectively showcases the delicate balance between hydrophilicity and lipophilicity, crucial for understanding a compound’s permeability across biological membranes. Compounds exhibiting lower tPSA values and higher cLogP values tend to display enhanced membrane permeability, owing to their greater affinity for lipid bilayers. DiSPASO shows medium membrane permeability and hydrophobicity properties compared to other crosslinker reagents and is in line with the BSP crosslinker published in Gao *et al*.^*36,37*^.

**Figure 2:**
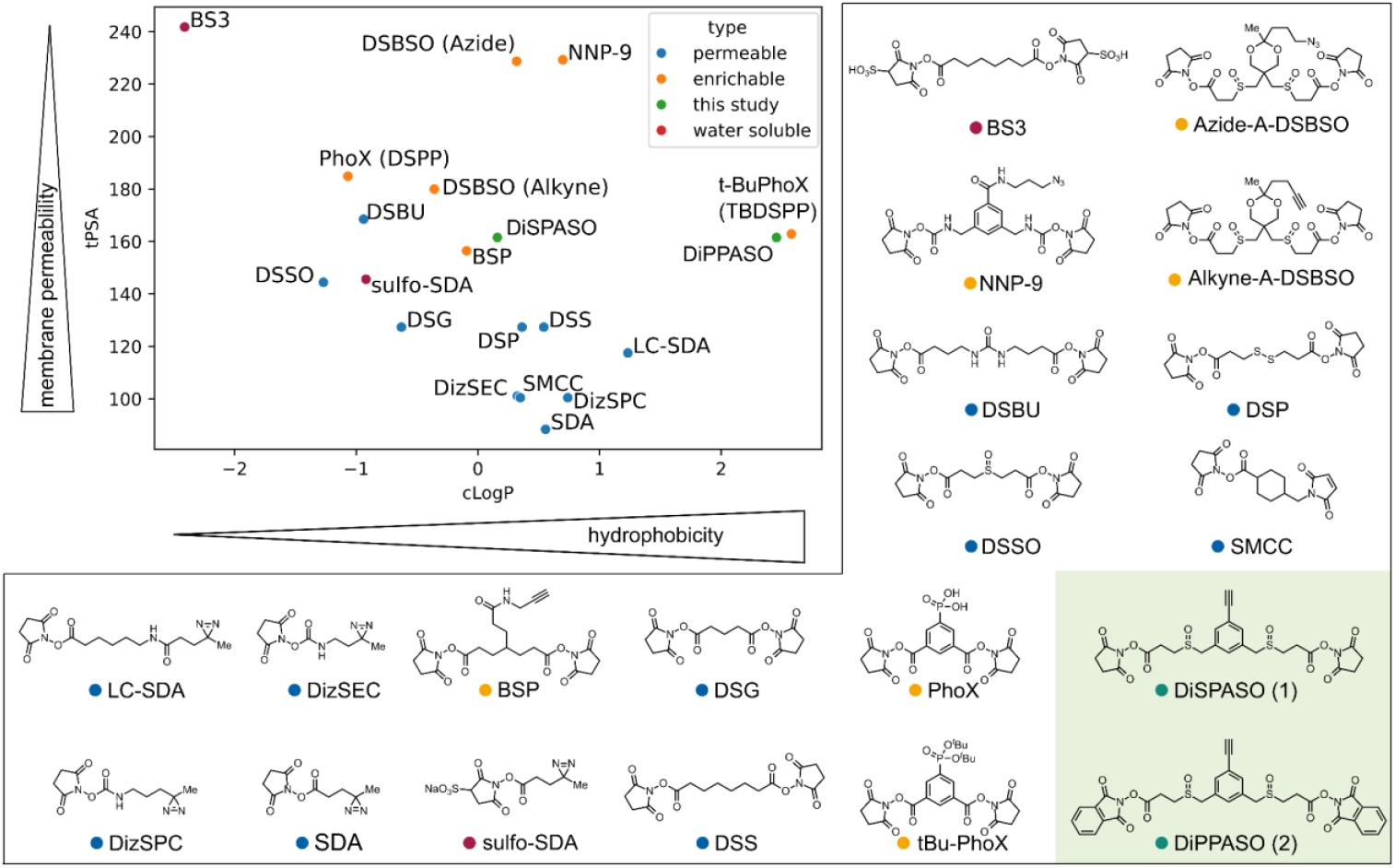
Topological Polar Surface area plot of commonly used crosslinker reagents. DiSPASO is shown in the centre of the plot (green dot) with moderate membrane permeability and hydrophobicity compared to other common crosslinker regents. DiPPASO (not presented in this study) shows similar properties to t-BuPhoX (TBDSPP) with a high hydrophobicity compared to other crosslinkers. The clouds of enrichable and permeable crosslinkers are clustered in two distinct parts of the plot, with DiSPASO at the interface between both groups. Classification of the crosslinkers was extracted from the Thermo Scientific crosslinker selection tool. If a crosslinker shows two properties of the classifications the more dominant property was selected for plotting.

Trifunctional crosslinkers, such as DiSPASO, enhance crosslinking mass spectrometry investigations by boosting sensitivity and specificity, primarily through an affinity tag that aids in enriching low-abundant crosslinked peptides that are typically masked by abundant linear peptides. This advancement is crucial for uncovering elusive protein interactions and enriching XL-MS data depth. DiSPASO’s effectiveness in cellular entry and membrane permeation was evaluated using three different enrichment strategies in increasing sample complexities starting from single peptide crosslinking towards live HEK cell crosslinking.

#### Comparison of DSBSO and DiSPASO crosslinking using Cas9-Halo

We optimized the affinity enrichment by assessing the following strategies (***Figure 3)***. The first enrichment employing picolyl azide ***(Figure 2A, (3)***) as a click handle and immobilized metal affinity chromatography (IMAC) for crosslinked peptide enrichment was tested using a single synthetic peptide (Ac-WGGGGR**K**SSAAR-COOH) with a defined crosslink-site (***Figure S1A***) and additionally using Cas9-Halo as the model protein (***Figure S2A***) to ensure a fully controlled environment. Both experiments demonstrated maximal crosslinked peptides/intensity when using 5 mM picolyl azide, with a click reaction efficiency of 99.3%. Click reaction efficiency was monitored by converting crosslinked products from DiSPASO crosslinking (***Figure S2A, blue bars***) to products formed after click reaction with picolyl azide (***Figure S2A, orange bars***), completing the click reaction when all DiSPASO links converted to click product BPNP (4) in ***Figure 3A***.

**Figure 3:**
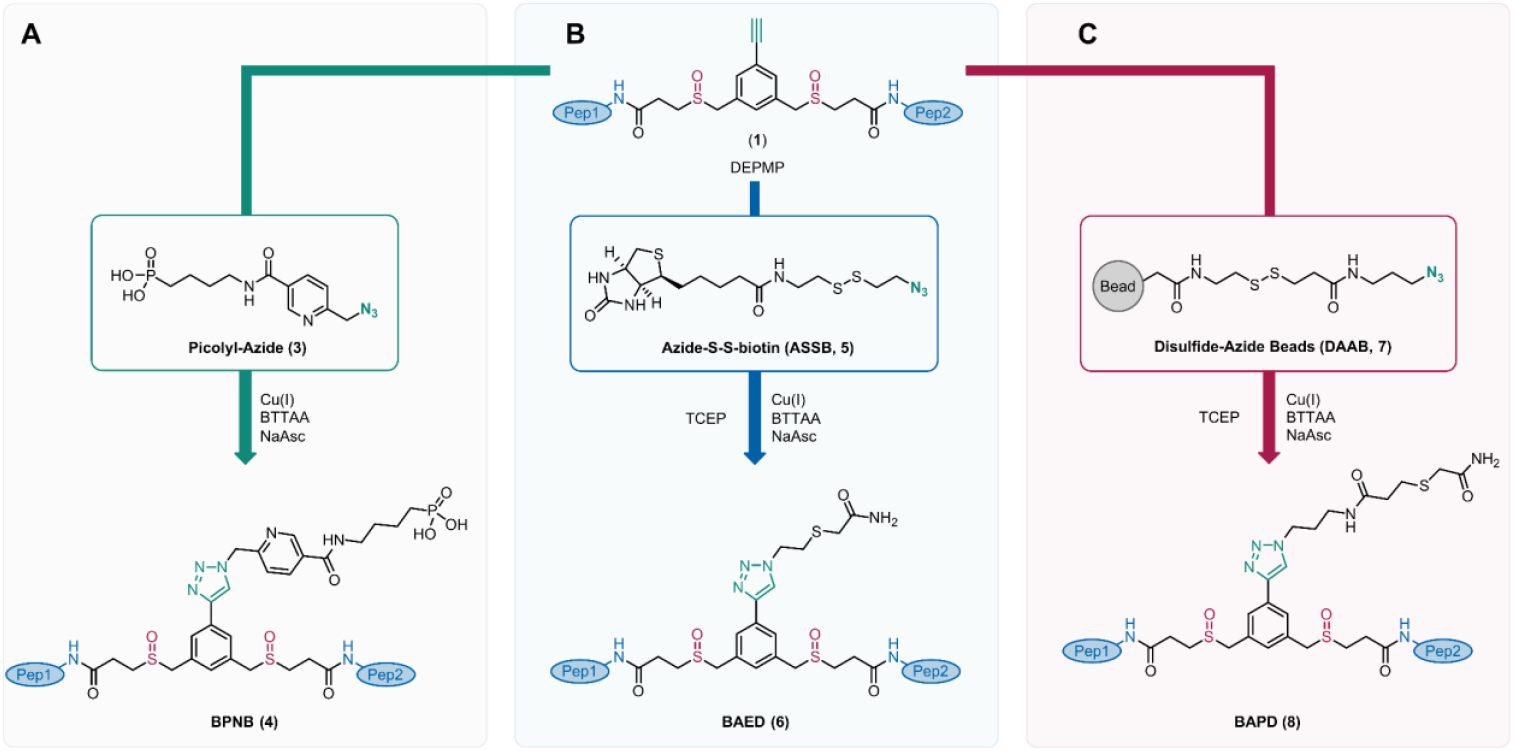
Overview of DiSPASO enrichment strategies. A: DiSPASO enrichment using a picolyl azide as a click chemistry reagent and an IMAC-based enrichment strategy to retrieve crosslinked peptides from a complex mixture. B: Azide-S-S-biotin (ASSB) based enrichment using ASSB as a click chemistry reagent and biotin-streptavidin bead strategy to enrich for crosslinked peptides. The elution of bead-bound crosslinked peptides is performed using the reduction of the disulfide bond of the ASSB compound. C: Simplified version of B. The click reagent Disulfide Azide is already coupled to the beads and the click reaction is taking place directly on the beads. Elution of crosslinked peptides is performed after the click reaction and washing procedure of the beads using a reducing reagent. IUPAC names of all compounds used in this manuscript are described in Table S4.

This principle was applied to all three enrichment strategies, tracking efficiency by monitoring mass shifts after each reaction’s conversion. Following click reaction optimization, the optimal higher-energy collision dissociation (HCD) was determined via single peptide crosslinking. The best-combined score for a crosslinked peptide was achieved with an HCD of 34 (***Figure S1B***), despite the main doublet intensity decreasing with higher HCDs (***Figure S1C***). A balance between peptide backbone fragmentation and doublet intensity required for crosslink identification was essential. Consequently, optimal fragmentation energies of 25, 27, and 32 were selected for further experiments.

The fragmentation pattern of DiSPASO was evaluated on Cas9 crosslinking data (***Figure 4***). Six potential fragments for the DiSPASO crosslinker were calculated based on the substitutions of the structures shown in ***Figure 4A***. The full crosslinker mass was also considered as fragment 7 to account for the theoretical cleavage of NH-crosslinker/ peptide bounds. Fragment ETFP (9), ETHMP (10) and alkene (11) represent the common and expected fragmentation pattern that is known from several fragmentation studies regarding DSBSO^30,44^ and DSSO^20,45,46^. Additionally, a second theoretical fragmentation pattern was tested due to low identification rates using DiSPASO for Cas9 crosslinking. The second fragmentation pattern includes a sulfenic acid fragment (13) or a thiol fragment (14) on the short side and an unexpected fragment EMP (12) on the long side of the crosslinker. Each fragment was defined separately according to the monoisotopic masses in ***Figure 4D&E***. Possible doublet distances were calculated according to equation 1 in the method section “Software adjustments”. This resulted in theoretical 15 doublet distances that can occur during fragmentation of DiSPASO. The same procedure was applied to DSBSO to compare the fragmentation behaviour of both crosslinkers.

**Figure 4:**
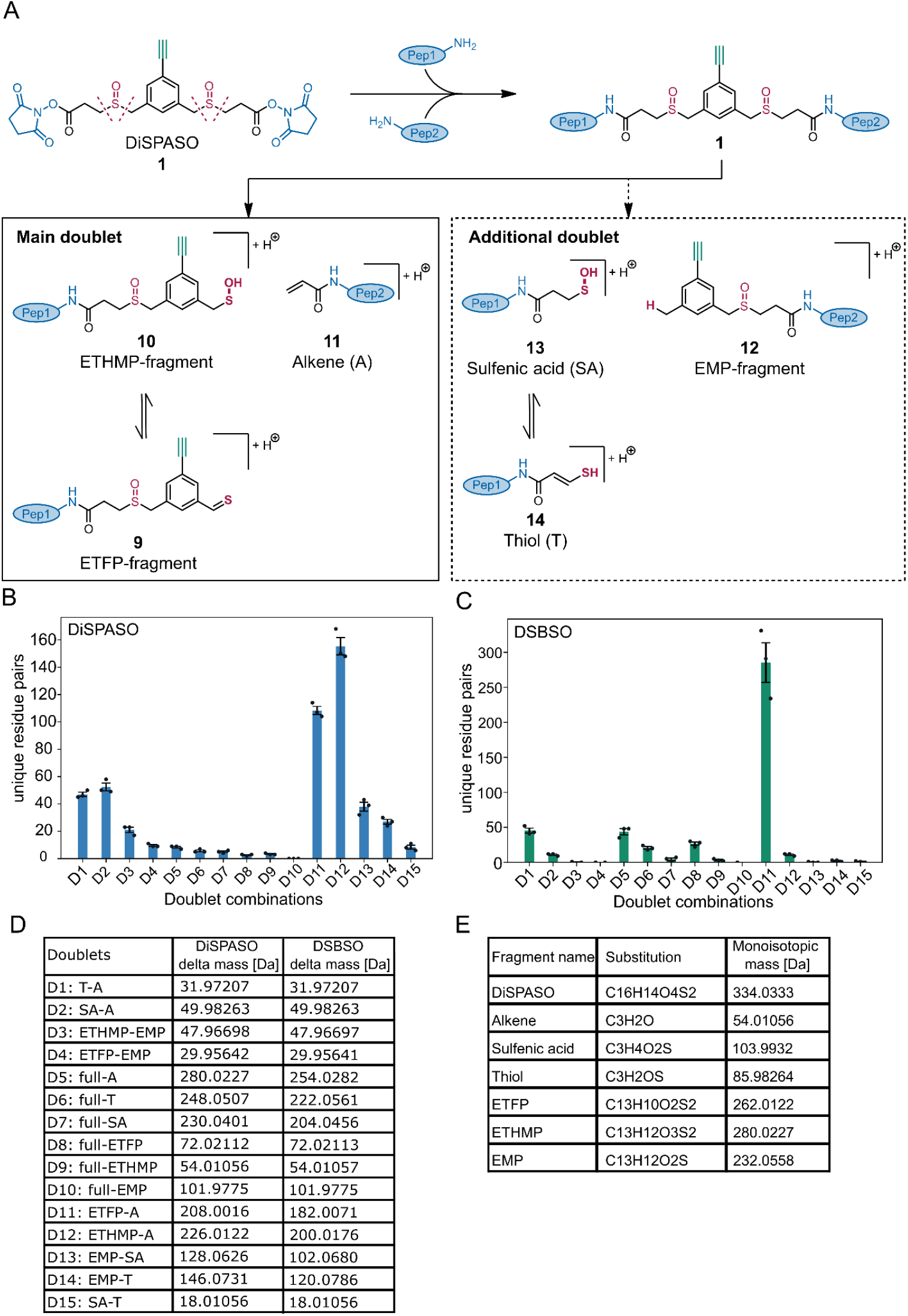
Possible fragments of DiSPASO during MS2 fragmentation and evaluation of MS Annika search setting regarding doublet distances. A: Fragmentation products of DiSPASO after high-energy collision dissociation (HCD) showing structures of DSSO-like cleavage products (alkene 11, ETFP 9 and ETHMP-fragments 10, main doublet) and possible additional doublets of the long side of the crosslinker (EMP 12 with SA 13 or T 14 as stubs, dotted grey box). B: Numbers of residue pairs identified from Cas9 crosslinked with DiSPASO while searching with each doublet distance separately. D12 (ETHMP-alkene doublet) shows the maximum number of identified links with D11 (ETFP-alkene) as the second and D1, D2 (alkene-thiol, sulfenic acid-alkene) as the third abundant doublets. C: Numbers of residue pairs identified from Cas9 crosslinked with DSBSO while searching with each doublet distance separately. D11 (ETFP-alkene doublet) shows the maximum number of identified links. D: Table of doublet definitions and delta masses of all possible doublets. E: Table of substitutions and monoisotopic masses of all fragments including DiSPASO as full construct bound to peptides. IUPAC names of all compounds used in this manuscript are described in Table S4.

Azide-DSBSO shows similar chemistry on the crosslinker backbone since it has the same length and cleavage site as DiSPASO. The center of Azide-DSBSO also comprises a ring system but in contrast to DiSPASO it is not a conjugated pi-ring system and instead of an alkyne functionality DSBSO employs an azide group for the enrichment and click chemistry. Hence, DSBSO was considered an ideal positive control to compare to the fragmentation pattern that was already published before^30,45^.

Surprisingly, DiSPASO showed various fragmentation sites with doublet D11 and D12 as the main occurring doublet distances. D11 represents the doublet distance between ETFP (9) and alkene (11) with a distance of 208 Da. The second main doublet is D12, ETHMP and alkene with a distance of 226.01 Da. Both doublets belong to the known fragmentation pattern published for DSSO before^34^. Additionally, DiSPASO shows doublets for the less likely fragmentation pattern between fragment EMP (12) and sulfenic acid (13) or thiol (14), D13 and D14 respectively, and alkene and thiol or sulfenic acid alone, D1 and D2 respectively. Whereas D13 and D14 represent 25% and 17% of the identified links, respectively, D1 and D2 account for 30% and 34%, highlighting the increased backbone fragmentation of the DiSPASO crosslinker. In contrast, DSBSO shows only one main doublet distance D11, which is defined by a distance of 182 Da between the alkene fragment and the long thiol fragment of DSBSO. D11 resulted in 285 identifiable links on average. D1 and D5 represent the second most abundant doublet distances with 16% and 15%, respectively. Although DSBSO also shows additional doublet distances, the main doublet distance stands out compared to all others. In conclusion, DiSPASO shows a different fragmentation pattern, that yields an almost complete scattering of the crosslinker backbone, compared to DSBSO, although the main cleavage sites are the same in perspective of the backbone. Hence, for further DiSPASO analysis the main doublet distances D11 and D12 were employed for crosslink identifications in MS Annika.

With all optimised parameters in place, the picolyl azide enrichment yielded 312 unique residue pairs compared to 259 identified residue pairs in the non-enriched control sample ***(Figure 5A***). The picolyl enrichment increased the number of links by 17% but the overall performance compared to DSBSO is lowered by 33% for 500ng (695 links DSBSO, 465 links DiSPASO) and 29% for 200ng injection amount (363 links DSBSO, 259 links DiSPASO). Nevertheless, the technical reproducibility of DiSPASO crosslinks before and after enrichment is high with 37% and 47% overlap between triplicates, respectively (***Figure 5B***). The recovery rate of crosslink before and after enrichment is 49% with 21% uniquely identified in the enriched sample.

**Figure 5:**
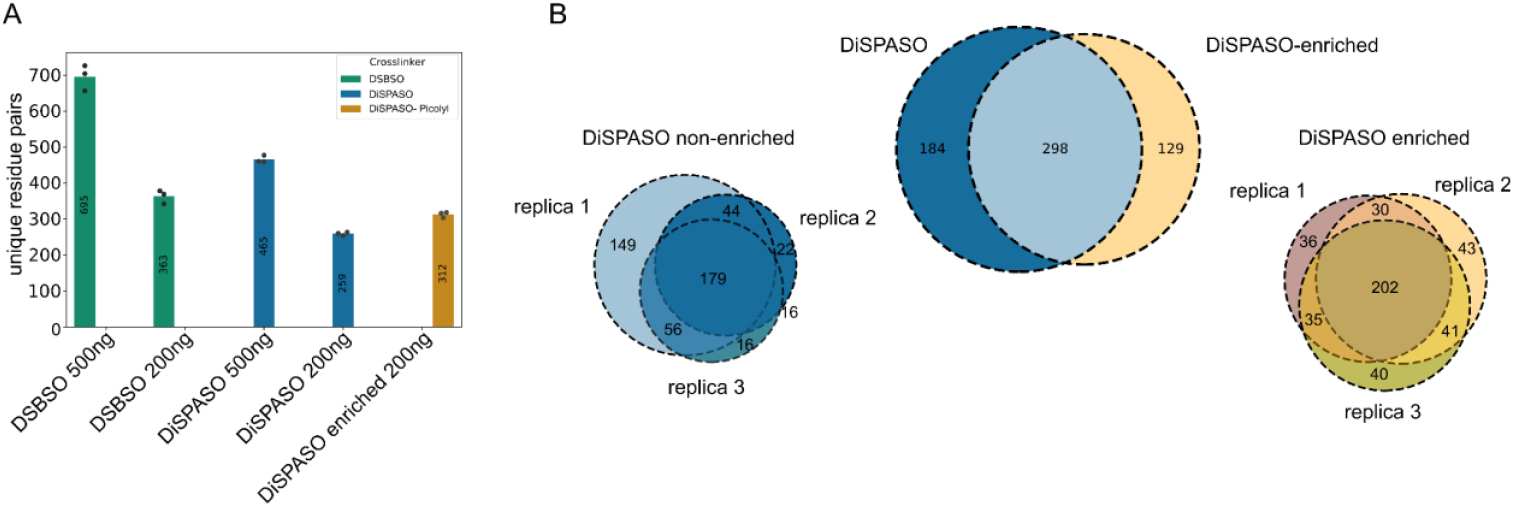
Comparison of DiSPASO and DSBSO using Cas9 as a model protein, as well as enrichment performance of DiSPASO using picolyl azide as enrichment tag. A: Number of identified unique crosslinks (unique residue pairs) in a triplicate experiment of Cas9 crosslinked with either DiSPASO or DSBSO in different injection amounts (500 and 200 ng) Blue indicates non-enriched crosslinked peptides, yellow enriched crosslinked peptides from Cas9 DiSPASO experiments. The green colour represents Cas9 crosslinking results using DSBSO. Dots on top of the bar show individual residue pair numbers of each replicate. The mean value of residue pairs is plotted in the middle of the bar. B: Overlap of identified crosslinks after click reaction and enrichment using picolyl azide as click chemistry reagent. After click reaction and enrichment using picolyl azide the numbers of crosslinks show an overlap of 49% with 21% unique to the enriched crosslinked sample. Left bottom of panel B: Overlap of technical replicates of non-enriched Cas9 links. Right bottom of panel B: Overlap of technical replicates of enriched Cas9 links.

The picolyl azide enrichment strategy was tested in a more complex system with crosslinked Cas9 as spike-in and HeLa cell lysate as background to increase the overall complexity of the sample (***Figure 6A, Figure S2B***). The spike-in amount ranged from 0.5 ug to 10 ug crosslinked Cas9 in 100 ug HeLa lysate. The lower the spike-in amount the fewer crosslinks could be identified after enrichment. For 0.5 ug only 14 crosslinks and in the 1:100 ratio (near in-cell condition) 20 crosslinks of a triplicate injection could be identified. In-cell crosslinking experiments in HeLa cells were performed to prove the hypothesis that the picolyl azide enrichment strategy is underperforming in real-case scenarios. Indeed, for HeLa in-cell crosslinking only 1 crosslink could be identified (***Figure 6A, grey background***). To improve the enrichment of crosslinked peptides in in-cell crosslinking experiments we moved to a different enrichment strategy using an Azide-S-S-biotin click chemistry handle.

**Figure 6:**
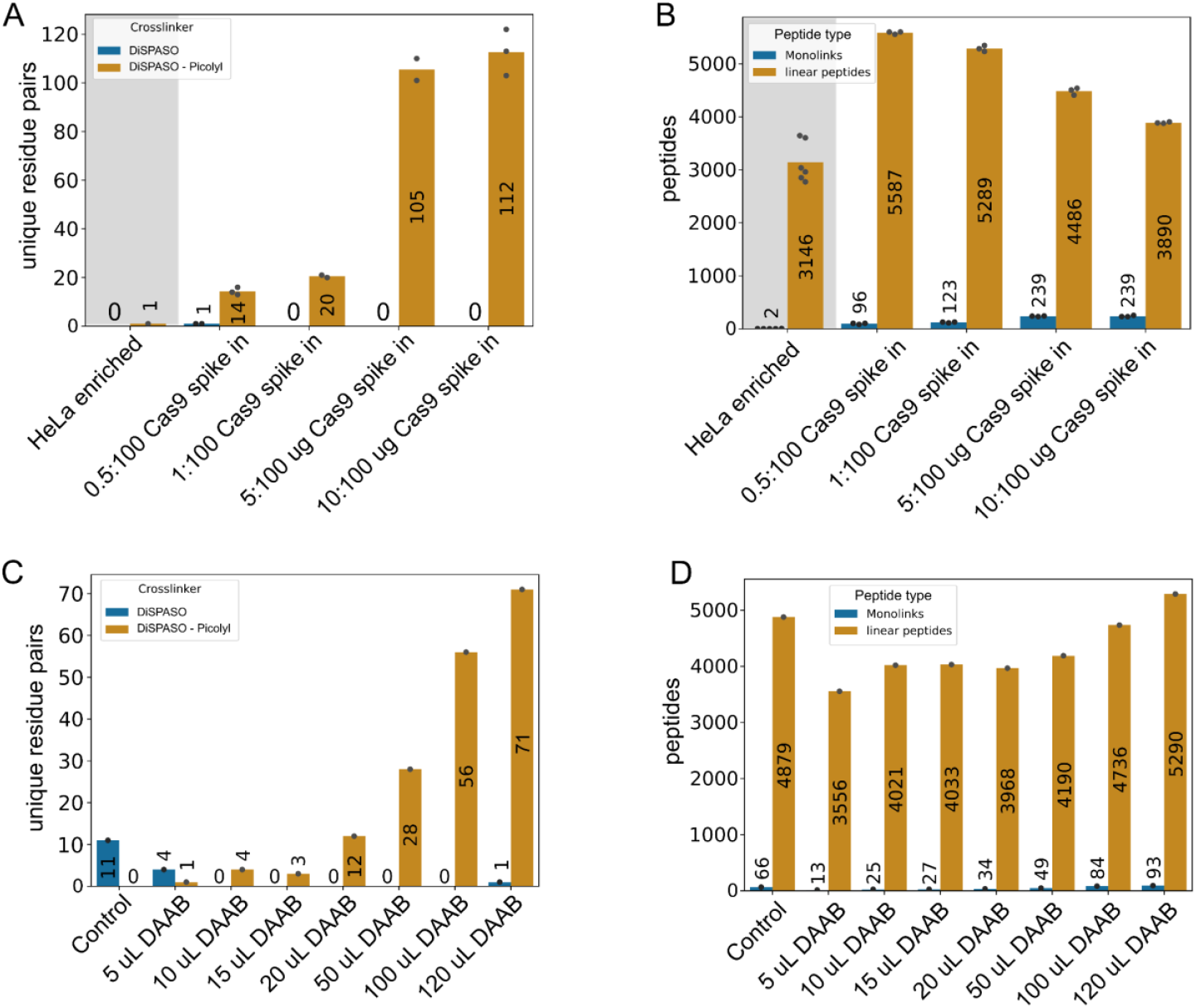
Application of DiSPASO enrichment strategies in increasing sample complexity. A: Spike -in experiment of crosslinked Cas9 in HeLa background. The Cas9 spike-in amount increased from 0.5 ug to 10 ug in a constant background of 100 ug HeLa. A HeLa in-cell crosslinking sample is used as a “control” sample. The amount of enriched Cas9 links increases with the amount of Cas9 spike-in. B: Analysis of Monolinks and linear peptides of the Cas9 spike-in experiment. The Monolinks and linear peptides of the “control” sample show in general fewer peptides due to the different experimental setup of in-cell crosslinking experiments in comparison to a spike-in experiment. The HeLa background peptides of the Cas9 spike-in decrease with increasing enrichable Cas9 crosslinks but cannot be depleted completely. C: HEK 293 in-cell crosslinking experiment using DiSPASO for crosslinking and Disulfide Azide Agarose beads (DAAB) for click chemistry-based enrichment of crosslinks. The bead amount is increased from 5 uL beads slurry to 120 uL, the control sample is crosslinked with DiSPASO without enrichment. With an increasing number of beads, the number of identifiable crosslinks after enrichment increases. D: Analysis of Monolinks and linear peptides of the in-cell crosslinking experiment. Linear peptides could not be depleted after crosslink enrichment.

### Azide-S-S-biotin and Disulfide azide agarose bead enrichment

Azide-S-S-biotin (ASSB), known for its effectiveness in protein and peptide labelling and enrichment, was utilized to enhance crosslinked peptide enrichment in in-cell crosslinking, leveraging its cleavable disulfide bond for peptide release post-enrichment (***Figure 3B***). The strong binding (free binding energy of −41.17 kcal/mol^47^, Kd of ∼10^−30^ M) of the biotin group to streptavidin beads can be fully exhausted, while binding efficiency can be granted due to the reduction of the disulfide bond. The resulting enriched and cleaved crosslinked peptide is small in size, which ensures minimal impact on ionization efficiency during the electrospray ionization (ESI) process.

ASSB’s optimization was conducted using the Cas9-Halo model, crosslinked with DiSPASO. Concentration titrations for ASSB were set from 1 mM to 20 mM, finding an optimal performance plateau at 10 mM with 175 unique residue pairs identified (***Figure S3A***). Despite varying ASSB concentrations, a consistent loss of crosslinked peptides was observed in the enrichment flowthrough, with a maximum loss of 333 links (***Figure S3B***). The concentration of sodium ascorbate (NaAsc), crucial for efficient click reactions without compromising the disulfide bond in ASSB, was also optimized, reaching a peak efficiency at 20 mM NaAsc with 242 unique links (***Figure S3C***), but with a significant sample loss of 39% after enrichment. Adjusting NaAsc to 30 mM aimed to balance click reaction efficiency with minimal side reactions. Bead volume optimization for enrichment used ranges from 10 µL to 100 µL of MBS bead slurry, with the highest number of detected links (217) at 70 µL (***Figure S3E)***, though not entirely preventing sample loss. The sample loss could be reduced by 56% (158 links 20 mM ASSB vs. 95 links 70 uL bead slurry) but not avoided, even after using 100 uL of bead slurry (***Figure S3F***).

Various bead types were tested (***Figure S4A & B***) without markedly reducing sample loss, leading to the adoption of a third strategy using disulfide azide agarose beads (***Figure 3C***), streamlining the workflow by clicking crosslinked peptides directly to beads. No additional purification steps are needed to avoid unspecific binding by free ASSB molecules and therefore blocking binding sites on the beads. All approaches, despite the methodical optimization and strategic shifts, underscored the inherent challenges in achieving efficient and loss-minimized enrichment of crosslinked peptides.

### In-cell crosslinking using DiSPASO

The picolyl azide enrichment strategy was challenged by crosslinking live HeLa cells directly in a 6-well plate using the DiSPASO crosslinker. Unfortunately, this experiment resulted in only 1 crosslink although several attempts were made to push this workflow towards success (***Figure 6A, grey background***).

Therefore, the ASSB enrichment workflow was applied to isolated commercial *E*.*coli* ribosomes in a complex HEK 293 cell background to proceed with the crosslink enrichment in live cells (***Figure S7A & B***). The ASSB enrichment strategy resulted in 14 ribosomal crosslinks that could be enriched from a 1:100 mixture (1 ug crosslinked ribosome spike-in in 100 ug HEK 293 cell lysate). This result was not efficient enough for in-cell crosslinking experiments and hence, we tried to improve the enrichment procedure by employing Disulfide azide agarose beads to directly click the crosslinked peptides to the enrichment handle and the beads in one step. We here titrated the bead amount using crosslinked HEK 293 cells. The cells were crosslinked directly in a 10 cm dish using 5 mM DiSPASO resulting in approximately 9e6 crosslinked cells. The sample was split to each condition equally with a bead amount of 5-120 uL tested (***Figure 6C***). 71 unique crosslinks could be detected from the 120 uL bead sample, the maximum of crosslinks in this experiment. The linear peptide background could not be reduced across the conditions, possibly due to the unspecific binding of peptides to the beads which is common in bead-based enrichment strategies. The problem of high linear peptide background stayed constant across all tested enrichment strategies and experiments, even with extensive washing procedures, which pinpoints a systematic problem for this kind of enrichment workflow (***Figure 6B & D)***.

Despite the promising membrane permeability and solubility properties of DiSPASO, mass spectrometry experiments yielded low identification rates in in-cell crosslinking studies. To investigate this discrepancy, we proceeded with confocal microscopy to directly assess the crosslinker’s performance in live cells on a visual basis.

### Membrane permeability assessment for DiSPASO in HEK cells

DiSPASO’s performance for in-cell crosslinking was further evaluated through confocal microscopy, showing rapid uptake by HEK 293 cells, visible in cell nuclei within minutes (***Figure 7A & Figure S5***). The HEK 293 cells, treated with 5 mM DiSPASO over 30 minutes, revealed a strong increase in signal within the first 5 minutes, reaching a plateau thereafter (***Figure 8A***).

**Figure 7:**
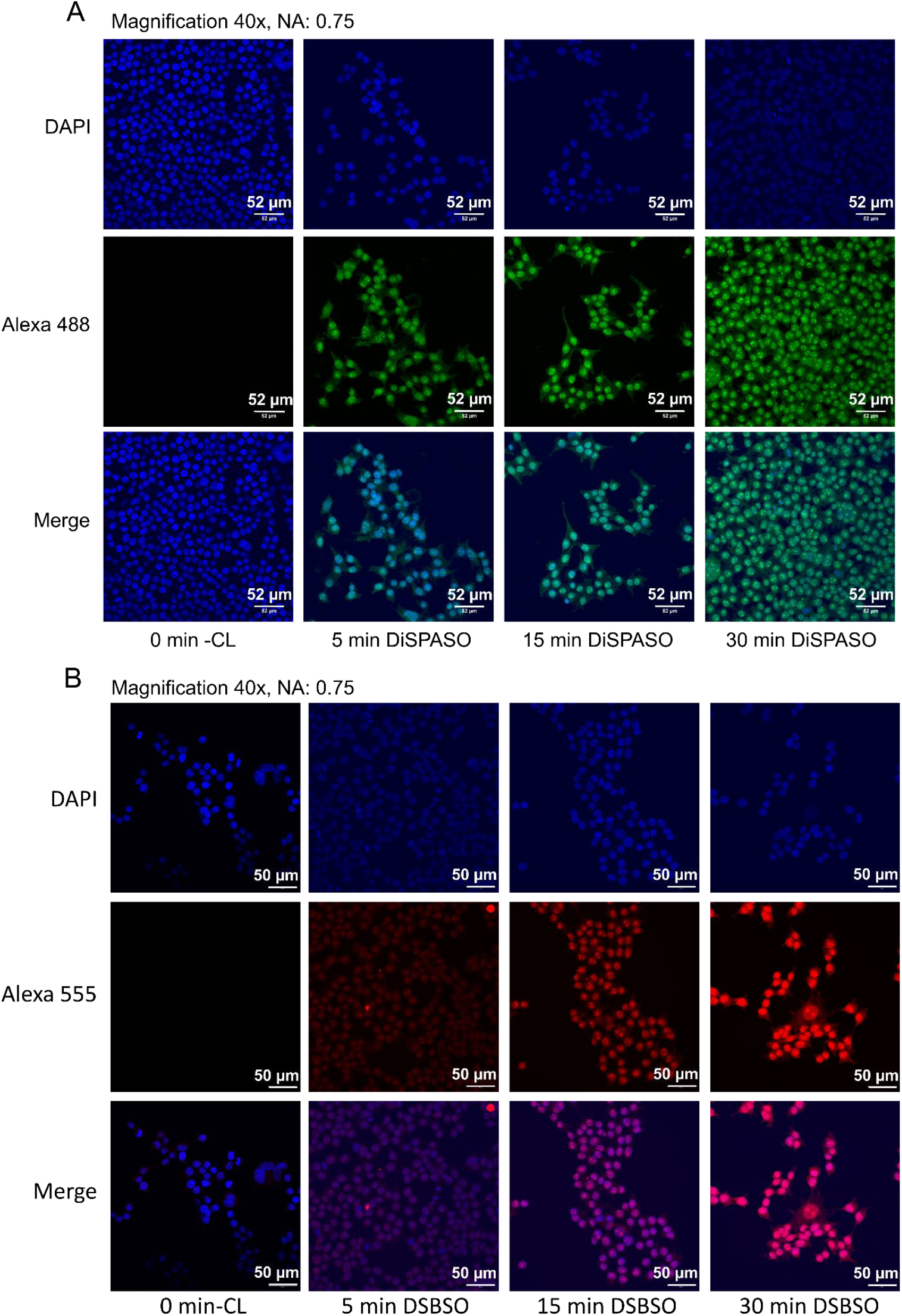
Comparison of DiSPASO and DSBSO uptake during in-cell crosslinking experiments in HEK 293 cells. A: Confocal microscopy images of DiSPASO during in-cell crosslinking experiments with a crosslink duration of 0 min (Control sample without crosslinking, first left panel), 5min (left second panel), 15 min (third left panel), 30min (last panel). The nuclei fluorescence signal of DAPI is shown in the upper panel in blue, fluorescence of crosslinked proteins after click reaction to Alexa 488 (green) in the middle panel and a merge of both channels on the bottom. The images were taken on an Olympus Spinning Disk Confocal microscope (2-024) using a magnification of 40 and a numerical aperture of 0.75. B: In comparison the DSBSO in-cell experiments were performed in the same way except for the fluorophore. DSBSO has an azide as click reaction handle and therefore Alexa 555 (red) was used to visualise crosslinked proteins. The crosslink duration was set to 0 min (Control sample without crosslinking, first left panel), 5 min (left second panel), 15 min (third left panel), 30 min (last panel). The nuclei fluorescence signal of DAPI is shown in the upper panel in blue, fluorescence of crosslinked proteins after click reaction to Alexa 555 (red) in the middle panel and a merge of both channels on the bottom. The images were also taken on an Olympus Spinning Disk Confocal microscope (2-024) using a magnification of 40 and a numerical aperture of 0.75.

**Figure 8:**
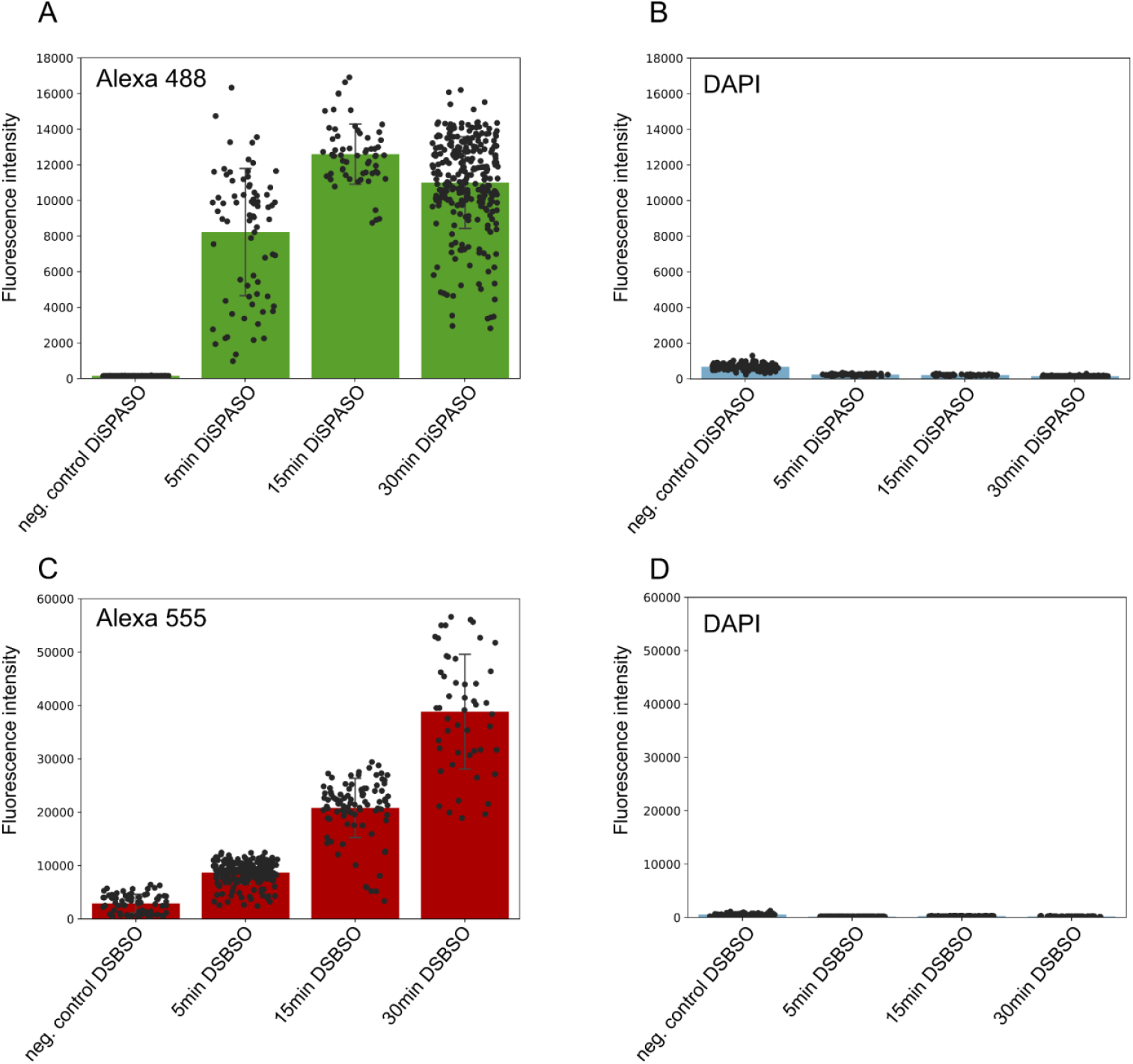
Quantitation of crosslink fluorescence signals within nuclei of DiSPASO vs. DSBSO microscopy images. A: Quantitation of the green Alexa 488 signal of DiSPASO crosslinked proteins. The fluorescence intensity increases fast within the first 5 min after reaching a plateau at 30 minutes. Each dot represents a signal intensity of crosslinked proteins in a nucleus. B: Nuclei control signal intensity of DAPI. The background intensity of DAPI is low in comparison to the intensity of crosslinked proteins. C: Quantitation of the red Alexa 555 signal of DSBSO crosslinked proteins. The fluorescence intensity increases slowly and reaches its maximum at 30min. D: Nuclei control signal intensity of DAPI. The background intensity of DAPI is also low in comparison to the intensity of crosslinked proteins. Quantitation of signal intensities of all images was performed in Fiji ImageJ (version 1.54f).

This indicates not only DiSPASO’s adeptness at penetrating cellular membranes but also its quick reactivity once inside the cell. Such attributes are critical for effective in-cell crosslinking, enabling the capture of a broad spectrum of protein-protein interactions in their native cellular environment. The microscopy images serve as a direct visual affirmation of DiSPASO’s capabilities, highlighting the controversy between the visual and the mass spectrometry results. To also evaluate the crosslinker performance compared to other commonly used in-cell crosslinking chemistry, we employed Azide-DSBSO^30,40^ as a benchmark reactant. For a fair comparison, HEK 293 cells were treated with 5 mM DSBSO in the same manner and parallel to the DiSPASO treatment. The difference in the click handle resulted in the use of two separate fluorescence dyes for each crosslinker. To label DiSPASO, the green dye AlexaFluor 488 and for DSBSO the red dye AlexaFluor 555 was used to visualise the performance of in-cell crosslinking (***Figure 7B***). In contrast to DiSPASO, which enters the cell very quickly, DSBSO needs longer incubation times and reaches its maximum performance after 30 min (***Figure 8C***). The extended reaction time may be attributed to the presence of the charged azide functionality incorporated into the crosslinker that decreases the membrane permeability of the crosslinker as shown in ***Figure 2***.

Our microscopy data, which validate the predicted chemical properties of DiSPASO, illustrate the crucial balance between hydrophilicity and lipophilicity that governs a compound’s permeability across biological membranes. While these visual confirmations align with theoretical expectations, they also highlight the challenges encountered during sample preparation and mass spectrometry analysis, revealing the complexity of translating theoretical advancements into practical applications. We hypothesize that the ability of the crosslinker to diffuse throughout the cell and crosslink proteins in all cellular compartments may depend on the hydrolysis of one reactive NHS-ester group, which increases its solubility. This could explain the observed discrepancy between the fluorescence signal, where the remaining NHS-ester reacts with proteins, and the low overall crosslinking yield in the mass spectrometry data, as indicated by the emergence of monolinks.

## Discussion

The introduction of DiSPASO in crosslinking mass spectrometry (XL-MS) has opened new avenues for elucidating protein-protein interactions (PPIs) within cells. DiSPASO has shown promising results via confocal microscopy, particularly with its successful cellular uptake and quick reactivity once inside the cell, highlighting its potential for in-depth cellular studies as well as revealing hurdles during mass spectrometry analysis.

In general, cleavable crosslinkers have labile bonds that break at lower energies than the peptide backbone, producing a distinct fragmentation pattern useful for analysing complex mixtures. However, an excess of cleavage sites can reduce the identification rate, as observed with DiSPASO in our study. Consequently, one of the two signal doublets resulting from crosslinker cleavage may be absent, possibly due to further fragmentation or decreased signal intensity caused by an excess of fragment species in the spectra^20,46,48^. In contrast to the anticipated cleavage pattern, our findings reveal that DiSPASO exhibits unexpected additional cleavage sites beyond those expected for DSBSO, resulting in the formation of extra fragments. While it has been documented that carbon−sulfur bonds in benzyl mercaptans can rapidly dissociate under specific conditions, such as UVPD irradiation at 213 nm and 266 nm^49,50^, this phenomenon was previously limited to certain wavelengths. Interestingly, recent studies have shown that C−S bond-selective photodissociation with 213 nm is augmented when sulfur is absent from an aromatic system by one methylene group (sp^3^ carbon) but hampered when sulfur is directly attached to a sp^2^ carbon. Surprisingly, our experiments indicate that this cleavage mechanism occurs even during standard HCD fragmentation.

Despite the challenges of complex fragmentation patterns and the need for refining data analysis, these findings offer valuable insights that drive further advancements in the field. The development of DiSPASO marks a significant step forward in enhancing cellular PPI mapping with high specificity and efficiency. However, the challenges encountered in translating its theoretical advantages into practical utility reveal a critical discrepancy that underscores the need for ongoing refinement in crosslinker technology. DiSPASO highlights the intricate balance between chemical innovation and biological functionality, emphasizing that advancements in crosslinker design must address current limitations to fully realize the potential of XL-MS in studying cellular mechanisms. While DiSPASO represents progress in PPI analysis, its limitations necessitate a cautious approach and highlight the complexity of developing effective crosslinking tools. Continuous innovation in structural design, particularly in simplifying crosslinkers and reducing potential cleavage sites, is essential. Additionally, advancements in computational tools and crosslinking search algorithms are crucial to overcoming challenges related to the search space in crosslinking data, enhancing the performance of non-cleavable crosslinkers for in-cell studies. These improvements will simplify data analysis and pave the way for more effective, streamlined XL-MS applications, ultimately bringing us closer to accurate and efficient mapping of PPI networks. Further studies involving different enrichment handles and reactive sites are ongoing to expand the capabilities of DiSPASO and similar crosslinkers.

## Data availability

The mass spectrometry proteomics data have been deposited to the ProteomeXchange Consortium (http://proteomecentral.proteomexchange.org) via the PRIDE partner repository^51^ with the dataset identifier PXD056091.

## Acknowledgements

This work was supported by the infrastructure funding 4*th* call 2022/01 (AT-SCP) of the Austrian Research Promotion Agency (FFG) and the project LS20-079 of the Vienna Science and Technology Fund (WWTF). This work was further funded by the ESPRIT program project number ESP 566 (Grant-DOI 10.55776/ESP566), P35045-B project (Grant-DOI 10.55776/P35045) and the F 8801-B Meiosis project (Grant-DOI 10.55776/F88) of the Austrian Science Fund (FWF). All LC-MS/MS analyses in Vienna were performed on the Vienna BioCenter Core Facilities instrument pool. We thank the Vienna Biocentre BioOptics facility for help and advice with microscopy imaging. Synthesis was performed at the Institute of Organic Chemistry of the University of Vienna. Funding from the Austrian Academy of Sciences (DOC Fellowship to B.R.B.) is acknowledged. We thank the University of Vienna for its generous support.

*This research was funded in whole, or in part, by the Austrian Science Fund (FWF). For open access, the author has applied a CC BY public copyright license to any Author Accepted Manuscript version arising from this submission*.

## Ethics approval and consent to participate

Not applicable.

## Consent for publication

Not applicable.

## References

1. Tang, X., Wippel, H. H., Chavez, J. D. & Bruce, J. E. Crosslinking mass spectrometry: A link between structural biology and systems biology. Protein Sci 30, 773–784 (2021).

2. Yu, C. & Huang, L. New advances in cross-linking mass spectrometry toward structural systems biology. Curr Opin Chem Biol 76, 102357 (2023).

3. Kalathiya, U. et al. Interfaces with Structure Dynamics of the Workhorses from Cells Revealed through Cross-Linking Mass Spectrometry (CLMS). Biomolecules 11, (2021).

4. Piersimoni, L., Kastritis, P. L., Arlt, C. & Sinz, A. Cross-Linking Mass Spectrometry for Investigating Protein Conformations and Protein-Protein Interactions-A Method for All Seasons. Chem Rev 122, 7500–7531 (2022).

5. O’Reilly, F. J. & Rappsilber, J. Cross-linking mass spectrometry: methods and applications in structural, molecular and systems biology. Nat. Struct. Mol. Biol. 25, 1000–1008 (2018).

6. de Jong, L., Roseboom, W. & Kramer, G. Towards low false discovery rate estimation for protein-protein interactions detected by chemical cross-linking. Biochim Biophys Acta Proteins Proteom 1869, 140655 (2021).

7. Low, T. Y. et al. Recent progress in mass spectrometry-based strategies for elucidating protein-protein interactions. Cell Mol Life Sci 78, 5325–5339 (2021).

8. Pal, S., Ganesan, K. & Eswaran, S. Chemical Crosslinking-Mass Spectrometry (CXL-MS) for Proteomics, Antibody-Drug Conjugates (ADCs) and Cryo-Electron Microscopy (cryo-EM). IUBMB Life 70, 947–960 (2018).

9. Zhang, Z. et al. Progress, Challenges and Opportunities of NMR and XL-MS for Cellular Structural Biology. JACS Au 4, 369–383 (2024).

10. Graziadei, A. & Rappsilber, J. Leveraging crosslinking mass spectrometry in structural and cell biology. Structure 30, 37–54 (2022).

11. Schneider, M., Belsom, A. & Rappsilber, J. Protein tertiary structure by crosslinking/mass spectrometry. Trends Biochem. Sci. 43, 157–169 (2018).

12. Matzinger, M. & Mechtler, K. Cleavable Cross-Linkers and Mass Spectrometry for the Ultimate Task of Profiling Protein-Protein Interaction Networks. J. Proteome Res. 20, 78–93 (2021).

13. Sinz, A. Divide and conquer: cleavable cross-linkers to study protein conformation and protein-protein interactions. Anal. Bioanal. Chem. 409, 33–44 (2017).

14. Petrotchenko, E. V. & Borchers, C. H. Crosslinking combined with mass spectrometry for structural proteomics. Mass Spectrom. Rev. 29, 862–876 (2010).

15. Sinz, A. Cross-Linking/Mass Spectrometry for Studying Protein Structures and Protein-Protein Interactions: Where Are We Now and Where Should We Go from Here? Angew. Chem. Int. Ed Engl. 57, 6390–6396 (2018).

16. Yu, C. & Huang, L. Cross-linking mass spectrometry: An emerging technology for interactomics and structural biology. Anal. Chem. 90, 144–165 (2018).

17. Wheat, A. et al. Protein interaction landscapes revealed by advanced in vivo cross-linking-mass spectrometry. Proc. Natl. Acad. Sci. U. S. A. 118, e2023360118 (2021).

18. Yugandhar, K. et al. MaXLinker: Proteome-wide Cross-link Identifications with High Specificity and Sensitivity. Mol. Cell. Proteomics 19, 554–568 (2020).

19. Liu, F., Rijkers, D. T. S., Post, H. & Heck, A. J. R. Proteome-wide profiling of protein assemblies by cross-linking mass spectrometry. Nat. Methods 12, 1179–1184 (2015).

20. Liu, F., Lössl, P., Scheltema, R., Viner, R. & Heck, A. J. R. Optimized fragmentation schemes and data analysis strategies for proteome-wide cross-link identification. Nat. Commun. 8, 15473 (2017).

21. Götze, M., Iacobucci, C., Ihling, C. H. & Sinz, A. A Simple Cross-Linking/Mass Spectrometry Workflow for Studying System-wide Protein Interactions. Anal. Chem. 91, 10236–10244 (2019).

22. Lenz, S. et al. Reliable identification of protein-protein interactions by crosslinking mass spectrometry. Nat. Commun. 12, 3564 (2021).

23. Bogdanow, B. et al. Enhancing inter-link coverage in cross-linking mass spectrometry through context-sensitive subgrouping and decoy fusion. bioRxiv (2023) doi:10.1101/2023.07.19.549678.

24. Giese, S. H., Fischer, L. & Rappsilber, J. A Study into the Collision-induced Dissociation (CID) Behavior of Cross-Linked Peptides. Mol. Cell. Proteomics 15, 1094–1104 (2016).

25. Trester-Zedlitz, M. et al. A modular cross-linking approach for exploring protein interactions. J. Am. Chem. Soc. 125, 2416–2425 (2003).

26. Zhang, H. et al. Identification of protein-protein interactions and topologies in living cells with chemical cross-linking and mass spectrometry. Mol. Cell. Proteomics 8, 409–420 (2009).

27. Tan, D. et al. Trifunctional cross-linker for mapping protein-protein interaction networks and comparing protein conformational states. eLife 5, e12509 (2016).

28. Sinz, A., Kalkhof, S. & Ihling, C. Mapping protein interfaces by a trifunctional cross-linker combined with MALDI-TOF and ESI-FTICR mass spectrometry. J. Am. Soc. Mass Spectrom. 16, 1921–1931 (2005).

29. Chu, F., Mahrus, S., Craik, C. S. & Burlingame, A. L. Isotope-coded and affinity-tagged cross-linking (ICATXL): an efficient strategy to probe protein interaction surfaces. J. Am. Chem. Soc. 128, 10362–10363 (2006).

30. Kaake, R. M. et al. A new in vivo cross-linking mass spectrometry platform to define protein-protein interactions in living cells. Mol. Cell. Proteomics 13, 3533–3543 (2014).

31. Schweppe, D. K. et al. Mitochondrial protein interactome elucidated by chemical cross-linking mass spectrometry. Proc. Natl. Acad. Sci. U. S. A. 114, 1732–1737 (2017).

32. Makepeace, K. A. T. et al. Improving Identification of In-Organello Protein-Protein Interactions Using an Affinity-Enrichable, Isotopically Coded, and Mass Spectrometry-Cleavable Chemical Crosslinker. (2020).

33. Petrotchenko, E. V., Serpa, J. J. & Borchers, C. H. An isotopically coded CID-cleavable biotinylated cross-linker for structural proteomics. Mol. Cell. Proteomics 10, M110.001420 (2011).

34. Chowdhury, S. M. et al. Identification of cross-linked peptides after click-based enrichment using sequential collision-induced dissociation and electron transfer dissociation tandem mass spectrometry. Anal. Chem. 81, 5524–5532 (2009).

35. Stadlmeier, M., Runtsch, L. S., Streshnev, F., Wühr, M. & Carell, T. A Click-Chemistry-Based Enrichable Crosslinker for Structural and Protein Interaction Analysis by Mass Spectrometry. Chembiochem 21, 103–107 (2020).

36. Zhao, L. et al. Enhanced protein-protein interaction network construction promoted by cross-linking with acid-cleavable click-chemistry enrichment. Front Chem 10, 994572 (2022).

37. Gao, H. et al. In-Depth Crosslinking in Minutes by a Compact, Membrane-Permeable, and Alkynyl-Enrichable Crosslinker. Anal. Chem. 94, 7551–7558 (2022).

38. An, Y. et al. Selective Removal of Unhydrolyzed Monolinked Peptides from Enriched Crosslinked Peptides To Improve the Coverage of Protein Complex Analysis. Anal. Chem. 94, 3904–3913 (2022).

39. Zhang, B. et al. Improved Cross-Linking Coverage for Protein Complexes Containing Low Levels of Lysine by Using an Enrichable Photo-Cross-Linker. Anal. Chem. 95, 9445–9452 (2023).

40. Matzinger, M., Kandioller, W., Doppler, P., Heiss, E. H. & Mechtler, K. Fast and Highly Efficient Affinity Enrichment of Azide-A-DSBSO Cross-Linked Peptides. J. Proteome Res. 19, 2071–2079 (2020).

41. Buncherd, H. et al. Selective enrichment and identification of cross-linked peptides to study 3-D structures of protein complexes by mass spectrometry. J. Proteomics 75, 2205–2215 (2012).

42. Steigenberger, B., Pieters, R. J., Heck, A. J. R. & Scheltema, R. A. PhoX: An IMAC-Enrichable Cross-Linking Reagent. ACS Cent Sci 5, 1514–1522 (2019).

43. Jiang, P.-L. et al. A Membrane-Permeable and Immobilized Metal Affinity Chromatography (IMAC) Enrichable Cross-Linking Reagent to Advance In Vivo Cross-Linking Mass Spectrometry. Angew. Chem. Int. Ed Engl. 61, e202113937 (2022).

44. Burke, A. M. et al. Synthesis of two new enrichable and MS-cleavable cross-linkers to define protein-protein interactions by mass spectrometry. Org. Biomol. Chem. 13, 5030– 5037 (2015).

45. Kao, A. et al. Development of a novel cross-linking strategy for fast and accurate identification of cross-linked peptides of protein complexes. Mol. Cell. Proteomics 10, M110.002212 (2011).

46. Kolbowski, L. et al. Improved Peptide Backbone Fragmentation Is the Primary Advantage of MS-Cleavable Crosslinkers. Anal. Chem. 94, 7779–7786 (2022).

47. Liu, F., Zhang, J. Z. H. & Mei, Y. The origin of the cooperativity in the streptavidin-biotin system: A computational investigation through molecular dynamics simulations. Sci Rep 6, 27190 (2016).

48. Kolbowski, L., Belsom, A., Pérez-López, A. M., Ly, T. & Rappsilber, J. Light-Induced Orthogonal Fragmentation of Crosslinked Peptides. JACS Au 3, 2123–2130 (2023).

49. Talbert, L. E. & Julian, R. R. Directed-Backbone Dissociation Following Bond-Specific Carbon-Sulfur UVPD at 213 nm. J. Am. Soc. Mass Spectrom. 29, 1760–1767 (2018).

50. Diedrich, J. K. & Julian, R. R. Site-specific radical directed dissociation of peptides at phosphorylated residues. J. Am. Chem. Soc. 130, 12212–12213 (2008).

51. Perez-Riverol, Y. et al. The PRIDE database resources in 2022: a hub for mass spectrometry-based proteomics evidences. Nucleic Acids Res. 50, D543–D552 (2022).

