## Supplemental Information for "A journey towards developing a new cleavable crosslinker reagent for in-cell crosslinking"

#### Supplement

**Organic Synthesis** ... 3
General Information ... 3
Synthesis of DiSPASO ... 4
General Procedure 1: Synthesis of Aryl Bromide (GP-1) ... 5
General Procedure 2: Reduction of Esters (GP-2) ... 6
General Procedure 3: Halogenation of Alcohols (GP-3) ... 6
General Procedure 4: Synthesis of Key Intermediate S5 (GP-4) ... 7
General Procedure 5: Sonogashira Cross-Coupling (GP-5) ... 8
General Procedure 6: Global Deprotection (GP-6) ... 9
General Procedure 7: Synthesis of NHS Protected Precursor S8 (GP-7) ... 10
General Procedure 8: Synthesis of NHP Protected Precursor (GP-8) ... 11
General Procedure 9: Synthesis of DiSPASO and DiPPASO (GP-9) ... 13
NMR Spectra ... 15
**Methods** ... 26
Reagents ... 26
Peptide Synthesis ... 26
Crosslinking reaction for Cas9 ... 27
Single peptide crosslinking ... 27
In-Solution Digest ... 27
Click reaction using Azide-S-S-biotin ... 27
Crosslinked peptide enrichment ... 28
Ribosome crosslinked with DiSPASO ... 28
Sensitivity experiment using picolyl azide as click reagent ... 29
In-cell crosslinking of HEK and HeLa cells using DiSPASO ... 30
Crosslink enrichment using Disulfide Azide Agarose beads (DAAB) ... 31
Sample preparation for confocal microscopy of crosslinked HEK cells ... 31
Confocal microscopy procedure ... 31
Relative quantitation of fluorescence signals ... 32
Mass spectrometry ... 32
Data analysis ... 33
Software adjustments ... 33
Surface area plot creation ... 34
**Supplemental figures** ... 34
**References** ... 42

#### List of Figures

- Scheme 1: DiSPASO synthetic route.
- Scheme 2: DiPPASO synthetic route from common precursor S7.
- Figure S1: Single peptide evaluation of DiSPASO.
- Figure S2: Evaluation of picolyl concentration for click reaction and enrichment sensitivity in HeLa background.
- Figure S3: Optimization of Azide-S-S-biotin, sodium ascorbate and bead amount to achieve optimal click and enrichment performance.
- Figure S4: Comparison of different bead types.
- Figure S5: Confocal microscopy pictures of crosslinked HEK 293 cells using DiSPASO.
- Figure S6: Exemplary workflow of an MS/MS2 search with MS Annika 2.0 in Proteome Discoverer.
- Figure S7: Application of ASSB-DiSPASO enrichment strategies of spike-in ribosome samples.

#### List of Tables

- Table S1. Special reagents used for DiSPASO click-reaction, enrichment, and microscopy.
- Table S2: Fragment names, substitution, and monoisotopic masses of DiSPASO fragments used for crosslinking search.
- Table S3: Search parameters for linear and crosslink search. Parameters not listed here were left at default settings
- Table S4: IUPAC and supplier names of chemical compounds and their abbreviations used in this manuscript.

### Organic Synthesis

#### General Information

**General Procedures.** All reactions were performed in round-bottom flasks or vials fitted with rubber septa and with magnetic stirring, unless otherwise stated. Reaction vessels were flushed with argon prior to use, unless otherwise stated. Liquids and solutions were transferred via syringe. All reactions were performed using anhydrous solvents obtained from Acros Organics, TCI or Sigma-Aldrich. Reaction progress was monitored by thin layer chromatography (TLC) performed on aluminum plates coated with silica gel $F_{254}$ with 0.2 mm thickness. Chromatograms were visualized by fluorescence quenching with UV light at 254 nm or by staining using potassium permanganate, followed by heating. Flash column chromatography was performed using silica gel 60 (230-400 mesh, Merck and co.), or pre-packed columns and reagent grade solvents.

**Materials**. All commercial reagents and solvents were used without further purification.

**Instrumentation**. All $^{1}$H NMR, $^{13}$C DEPTQ-135 NMR, $^{13}$C CPD NMR and $^{19}$F NMR spectra were recorded using a Bruker AV-400, AV-500, AV-600 or AV-700 spectrometer at 300 K. Chemical shifts ($\delta$) were given in parts per million (ppm), referenced to the solvent peak of $CDCl_3$, defined at $\delta$ = 7.26 ppm ($^{1}$H NMR) and $\delta$ = 77.16 ppm ($^{13}$C NMR) and the solvent peak of DMSO-$d_6$, defined at $\delta$ = 2.50 ppm ($^{1}$H NMR) and $\delta$ = 39.52 ppm ($^{13}$C NMR)[1]. Coupling constants ($J$) are reported in Hertz (Hz). $^{1}$H NMR splitting patterns are designated as singlet (s), doublet (d), triplet (t), quartet (q), quintet (quint.) or a combination thereof, as they appeared in the spectrum. If the appearance of a signal differs from the expected splitting pattern, the observed pattern is designated as apparent (app). Splitting patterns that could not be interpreted or easily visualized are designated as multiplet (m) or broad (br). Infrared (IR) spectra were obtained using Perkin-Elmer Spectrum 100 FT-IR spectrometer. Wavenumbers ($\nu_{max}$) are reported in cm$^{-1}$. Mass spectra were obtained using a Bruker maXis UHR-TOF spectrometer (70 eV), using electrospray ionization (ESI) or atmospheric-pressure chemical ionization (APCI) or an Agilent 7200B GC/Q-TOF spectrometer (70 eV), using electron ionization (EI). Optical rotations were measured on a Perkin Elmer 341 polarimeter using a 100 mm path-length cell at 589 nm ($c$ given in g / (100 mL)). Details of chromatographic conditions are indicated under each compound.

#### Synthesis of DiSPASO

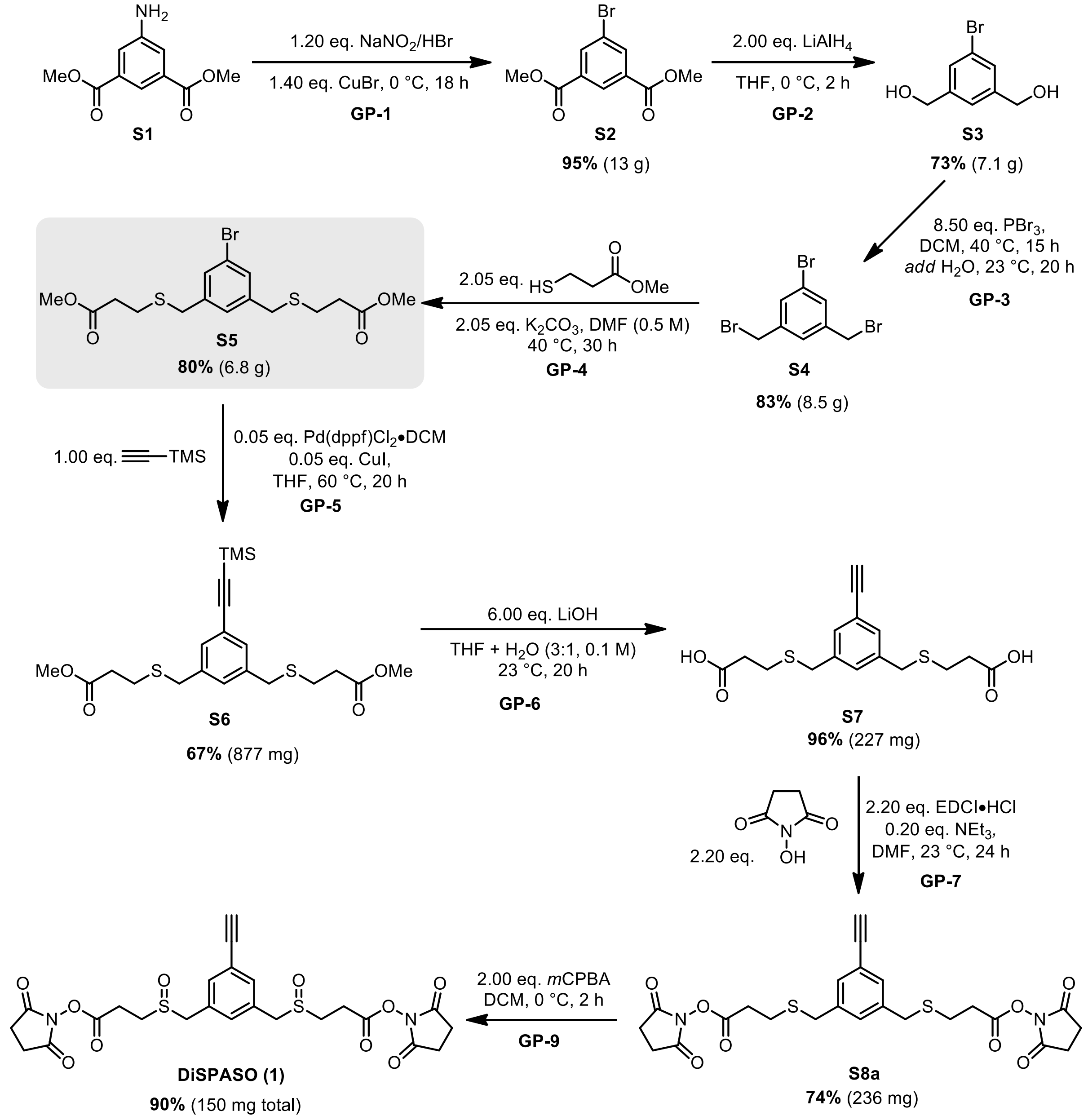

***Scheme 1:*** *DiSPASO synthetic route.*

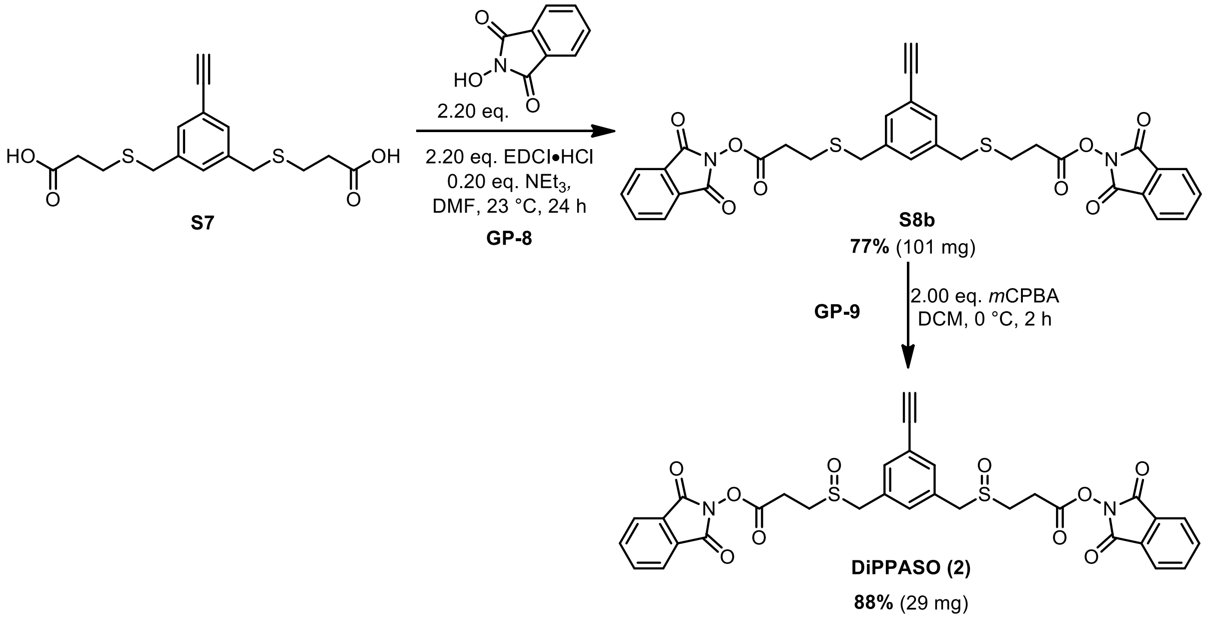

***Scheme 2:** DiPPASO synthetic route from common precursor **S7**.*

#### General Procedure 1: Synthesis of Aryl Bromide (GP-1)[2,3]

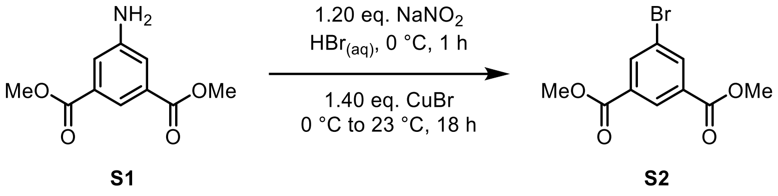

5-Amino-isophthalic acid dimethyl ester (**S1**) (1.00 eq., 50.0 mmol, 10.5 g) was added to a 1 L flask equipped with a magnetic stir bar and dissolved in 200 mL of a 15% aq. HBr solution (obtained by diluting 95 mL of a 48% aq. HBr solution with 205 mL water) at 0 °C. Then an aq. solution of $NaNO_2$ (1.20 eq., 60.0 mmol, 4.14 g in 20.0 mL dist. $H_2O$) was slowly added under vigorous stirring. After 10 minutes, the diazonium solution was added portion-wise at 0 °C to 1 L flask equipped with a stir bar containing a solution of CuBr (1.40 eq., 70.0 mmol, 10.0 g in 80.0 mL 15% aq. HBr solution) and the reaction mixture was allowed to warm up to 23 °C over 18 h. Then, the reaction mixture was diluted with ethyl acetate (200 mL) and the organic layer was separated, washed with distilled water (3 × 200 mL), dried with $MgSO_4$, filtered and concentrated under reduced pressure. The crude residue was used in the next step without further purification.

Dimethyl 5-bromoisophthalate (**S2**)

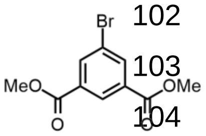

**S2**

According to general procedure **GP-1**, **S2** was isolated as a yellow solid (12.9 g, 47.2 mmol, 95%) and used in the next step without further purification. Spectroscopic data were in accordance with those reported in the literature[3].

**$^{1}$H NMR (400 MHz, $CDCl_3$):** *δ* 8.60 (s, 1H), 8.35 (d, *J* = 1.3 Hz, 2H), 3.96 (s, 6H) ppm.

General Procedure 2: Reduction of Esters (GP-2)

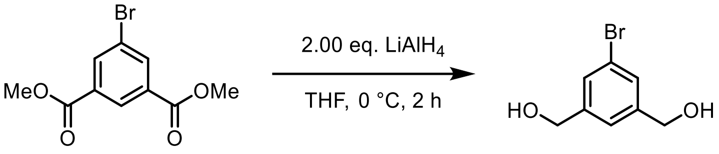

**S2** **S3**

To a 500 mL flask equipped with a magnetic stir bar and containing **S2** (1.00 eq., 45.0 mmol, 12.3 g) dissolved in THF (150 mL), were added $LiAlH_4$ pellets (2.00 eq., 90.0 mmol, 3.42 g) at 0 °C. The reaction was stirred at 0 °C for 2 h before being quenched by dropwise addition of sat. potassium sodium tartrate solution (100 mL). The organic phase was separated and the aqueous phase was extracted with diethyl ether (2 × 100 mL). The combined organic phases were dried with $MgSO_4$, filtered and concentrated under reduced pressure. The crude residue was purified by column chromatography.

(5-Bromo-1,3-phenylene)dimethanol (**S3**)

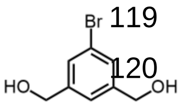

**S3**

According to general procedure **GP-2**, **S3** was isolated as a white solid (7.14 g, 32.9 mmol, 73%) after purification by column chromatography (DCM/MeOH = 19:1, $R_f$ = 0.29). Spectroscopic data were in accordance with those reported in the literature[3].

**$^{1}$H NMR (400 MHz, $CDCl_3$):** *δ* 7.45 (s, 2H), 7.29 (s, 1H), 4.69 (d, *J* = 5.9 Hz, 4H), 1.71 (t, *J* = 5.9 Hz, 2H) ppm.

General Procedure 3: Halogenation of Alcohols (GP-3)

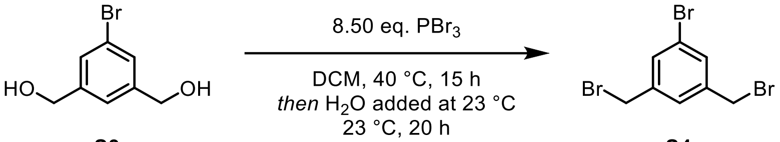

**S3** **S4**

To a 250 mL flame-dried flask equipped with a magnetic stir bar and fitted with a reflux condenser was added **S3**. (1.00 eq., 30.0 mmol, 6.5 g) and dry DCM (100 mL). A solution of $PBr_3$ (8.50 eq., 255 mmol, 24.7 mL) in dry DCM (100 mL) was added dropwise and the reaction mixture was stirred under reflux (oil bath at 40 °C) for 15 h. Then, the top of the reflux condenser was connected to a series of washing bottles (empty, 1 M NaOH solution, empty) and water (60.0 mL) was added. The reaction was stirred for 20 h at 23 °C and diluted with DCM (30.0 mL). The organic phase was separated. Then the aqueous phase was extracted with DCM (2 × 150 mL). The combined organic phases were dried with $MgSO_4$, filtered and concentrated under reduced pressure. The crude residue was purified by column chromatography.

<u>1-Bromo-3,5-bis(bromomethyl)benzene (**S4**)</u>

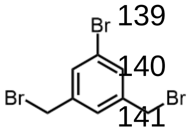
**S4**

According to general procedure **GP-3**, **S4** was isolated as a white solid (8.52 g, 24.9 mmol, 83%) after purification by column chromatography (heptane/ethyl acetate= 4:1, $R_f$ = 0.35). Spectroscopic data were in accordance with those reported in the literature[3].

**$^1$H NMR (400 MHz, $CDCl_3$):** $\delta$ 7.47 (s, 2H), 7.33 (d, $J$ = 4.3 Hz, 1H), 4.41 (s, 4H) ppm.

**$^{13}$C NMR (101 MHz, $CDCl_3$):** $\delta$ 140.5, 132.1, 128.4, 122.9, 31.6 ppm.

**LRMS ($EI^+$):** exact mass calculated for $[M+Na]^+$ ($C_8H_7Br_3Na$) requires *m/z* 341.8, found *m/z* 341.8.

**IR (neat) $\nu_{max}$:** = 3056, 3024, 2970, 2362, 2106, 1796, 1602, 1571, 1444, 1261, 1210, 1162, 1128, 1114, 998, 972, 893, 878, 865, 819, 740 cm$^{-1}$.

<u>General Procedure 4: Synthesis of Key Intermediate S5 (GP-4)</u>

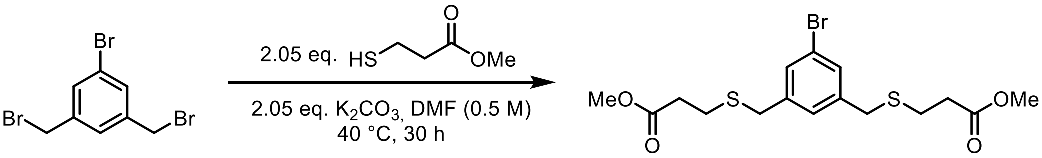

**S4** **S5**

To a 250 mL flame-dried flask equipped with a magnetic stir bar and reflux condenser, was added the tribromide **S4** (1.00 eq., 20.0 mmol, 6.86 g), methyl 3-mercaptopropanoate (2.05 eq., 41.0 mmol, 4.66 mL), $K_2CO_3$ (2.05 eq., 41.0 mmol, 5.67 g) and DMF (40.0 mL). The reaction mixture was heated to 40 °C for 30 h, then the reaction mixture was cooled down to 23 °C and diethyl ether was added (100 mL) and transferred to a separation funnel. The organic phase was washed with $H_2O$ (4 × 100 mL), then brine (3 × 100 mL), dried with $MgSO_4$, filtered, and concentrated under reduced pressure. The crude product was purified by column chromatography.

Dimethyl 3,3'-(((5-bromo-1,3-phenylene)bis(methylene))bis(sulfanediyl))dipropionate (**S5**)

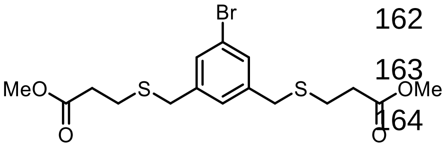

**S5**

According to general procedure **GP-4**, **S5** was isolated as a colourless oil (6.8 g, 16.1 mmol, 80%) after purification by column chromatography (heptane/ethyl acetate= 9:1 to 5:1, $R_f$ = 0.34 in 5:1).

**$^1$H NMR (400 MHz, $CDCl_3$):** $\delta$ 7.36 (d, $J$ = 1.0 Hz, 2H), 7.21 (s, 1H), 3.69 (s, 6H), 3.67 (s, 4H), 2.69 (dd, $J$ = 10.9, 3.7 Hz, 4H), 2.56 (t, $J$ = 7.1 Hz, 4H) ppm.

**$^{13}$C NMR (101 MHz, $CDCl_3$):** $\delta$ 172.3 (2C), 140.8 (2C), 130.7 (2C), 128.1, 122.8, 52.0 (2C), 35.9 (2C), 34.4 (2C), 26.5 (2C) ppm.

**HRMS (ESI$^+$):** exact mass calculated for $[M+Na]^+$ ($C_{16}H_{21}O_4BrNa$) requires $m/z$ 442.9957, found $m/z$ 442.9957.

**IR (neat) $\nu_{max}$:** = 2996, 2950, 2923, 2844, 1730, 1600, 1568, 1435, 1356, 1299, 1282, 2245, 1217, 1195, 1169, 1018, 979, 946, 893, 865, 820, 728 cm$^{-1}$.

General Procedure 5: Sonogashira Cross-Coupling (GP-5)

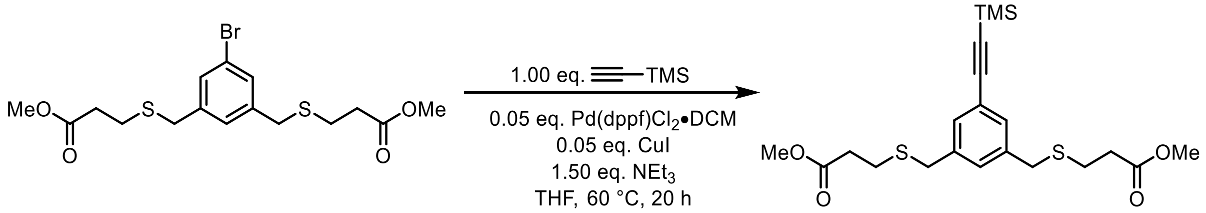

**S5** **S6**

Based on a previously reported procedure[4]. To a flame-dried Schlenk equipped with a magnetic stir bar were added aryl bromide **S5** (1.00 eq., 3.00 mmol, 1.26 g), Pd(dppf)$Cl_2$•DCM (0.05 eq., 0.15 mmol, 125 mg), CuI (0.05 eq., 0.15 mmol, 28.7 mg) and dry THF (10.0 mL) under inert atmosphere. The reaction mixture was degassed for 20 minutes using an Ar-filled balloon and an ultrasound bath. Then, trimethylsilylacetylene (1.00 eq., 3.00 mmol, 0.43 mL) and $NEt_3$ (1.50 eq., 4.50 mmol, 0.63 mL) were added and the sealed reaction mixture was stirred for 20 h at 60 °C. Next, the reaction mixture was filtered over a short pad of celite, washed with diethyl ether (20.0 mL), dried with $MgSO_4$, filtered and concentrated under reduced pressure. The crude residue was purified by column chromatography.

Dimethyl 3,3'-(((5-((trimethylsilyl)ethynyl)-1,3-phenylene)bis(methylene))bis(sulfanediyl))-dipropionate (**S6**)

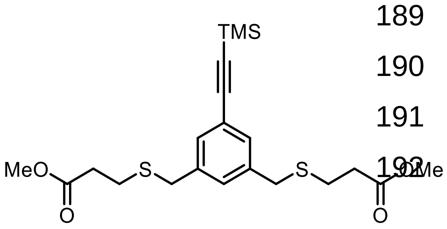

**S6**

According to general procedure **GP-5**[4], **S6** was isolated as a colourless oil (877 mg g, 2.00 mmol, 67%) after purification by column chromatography (heptane/ethyl acetate= 9:1, $R_f$ = 0.21)

**$^1$H NMR (400 MHz, $CDCl_3$):** *δ* 7.30 (s, 2H), 7.24 (s, 1H), 3.68 (d, *J* = 0.5 Hz, 6H), 3.67 (s, 4H), 2.66 (t, *J* = 7.2 Hz, 4H), 2.55 (t, *J* = 7.2 Hz, 4H), 0.24 (s, 9H) ppm.

**$^{13}$C NMR (101 MHz, $CDCl_3$):** *δ* 172.3 (2C), 138.8, 131.2 (2C), 129.7, 123.7, 104.6, 94.8, 51.9 (2C), 36.0 (2C), 34.4 (2C), 26.4 (2C), 0.1 (3C) ppm.

**HRMS (ESI$^+$):** exact mass calculated for $[M+H]^+$ ($C_{21}H_{31}O_4S_2Si$) requires *m/z* 439.1428, found *m/z* 439.1428.

**IR (neat) $\nu_{max}$:** = 2953, 2159, 1734, 1593, 1435, 1357, 1299, 1248, 1196, 1168, 1020, 980, 840, 759 $cm^{-1}$.

General Procedure 6: Global Deprotection (GP-6)

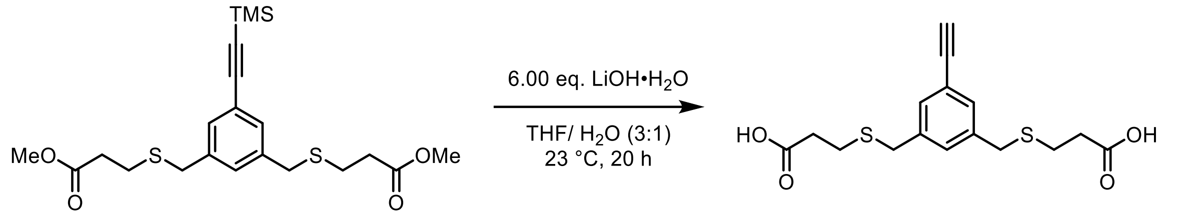

**S6** **S7**

In a flame-dried flask equipped with a magnetic stir bar **S6** (1.00 eq., 0.70 mmol, 307 mg) was dissolved in a THF/$H_2O$ mixture (3:1 *v*/*v*, 4.0 mL). LiOH•$H_2O$ (6.00 eq., 4.20 mmol, 176 mg) was added and the reaction was stirred at 23 °C for 24 h. The reaction was stopped by the addition of a 1 M HCl solution (5.0 mL) and the mixture was transferred to a separation funnel. The organic phase was separated, then the aqueous phase was washed with diethyl ether (4 × 5.0 mL). The combined organic phase was dried with $MgSO_4$, filtered and concentrated under reduced pressure. The crude residue was used for the next step without further purification.

3,3'-(((5-Ethynyl-1,3-phenylene)bis(methylene))bis(sulfanediyl))dipropionic acid (**S7**)

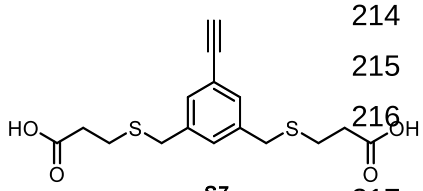

**S7**

According to general procedure **GP-6, S7** was isolated as a white solid (227 mg, 0.67 mmol, 96%) and used for the next step without further purification.

**$^1$H NMR (500 MHz, $CDCl_3$):** *δ* 7.35 (d, *J* = 1.2 Hz, 2H), 7.29 (s, 1H), 3.71 (s, 4H), 3.07 (s, 1H), 2.68 (t, *J* = 6.9 Hz, 4H), 2.58 (t, *J* = 6.9 Hz, 4H) ppm.

**$^{13}$C NMR (151 MHz, $CDCl_3$):** *δ* 177.9 (2C), 138.9 (2C), 131.4 (2C), 130.1, 122.9, 77.8, 68.1, 36.1 (2C), 34.5 (2C), 26.0 (2C) ppm.

**HRMS (ESI$^-$):** exact mass calculated for $[M-H]^-$ ($C_{16}H_{17}O_4S_2$) requires *m/z* 337.0574, found *m/z* 337.0575.

**IR (neat**) **$\nu_{max}$:** = 3231, 2916, 2664, 1688, 1590, 1420, 1403, 1337, 1302, 1264, 1237, 1197, 1161, 1141, 1041, 910, 880, 869, 805, 771 $cm^{-1}$.

General Procedure 7: Synthesis of NHS Protected Precursor S8 (GP-7)

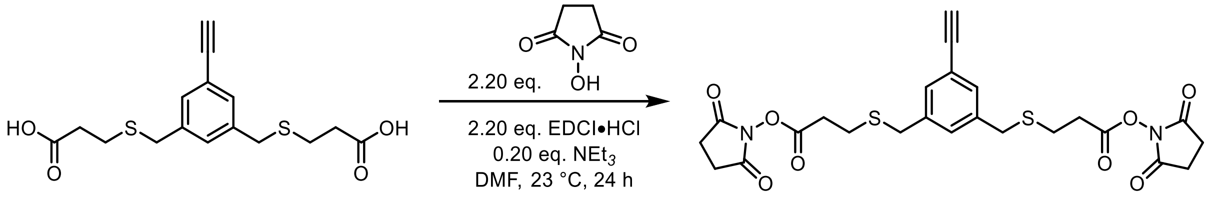

**S7** **S8a**

Inspired by previously reported procedure[5]. To flame-dried Schlenk equipped with a stir bar was added the diacid **S7** (1.00 eq., 0.60 mmol, 203 mg) and dry DMF (6.0 mL) under an inert atmosphere. Next, *N*-hydroxysuccinimide (2.20 eq., 1.32 mmol, 155 mg), 1-(3-dimethylaminopropyl)-3-ethylcarbodiimide hydrochloride (EDCI•HCl, 2.20 eq., 1.32 mmol, 253 mg) and $NEt_3$ (0.20 eq., 0.12 mmol, 17.0 µL) were sequentially added at 23 °C. The reaction mixture was stirred at 23 °C for 14 h. Then the reaction mixture was stopped by the addition of an aq. 1 M HCl solution (10.0 mL), diluted with ethyl acetate (10.0 mL) and transferred to a separation funnel. The organic phase was separated, and washed with sat. ammonium chloride solution (2 × 10.0 mL), distilled $H_2O$ (2 × 10.0 mL) and brine (2 × 10.0 mL), dried with $MgSO_4$, filtered and concentrated under reduced pressure. The crude residue was purified by column chromatography.

<u>Bis(2,5-dioxopyrrolidin-1-yl)3,3'-(((5-ethynyl-1,3-phenylene)bis(methylene)) bis(sulfanediyl))dipropionate (**S8a**)</u>

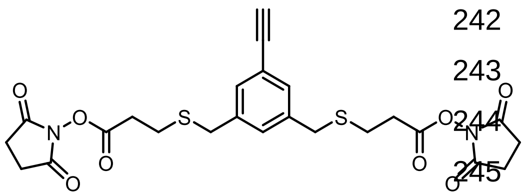

**S8a**

According to general procedure **GP-7**[5], **S8** was isolated as a colourless oil (236 mg, 0.44 mmol, 74%) after purification by column chromatography (heptane/ethyl acetate= 1:2, $R_f$ = 0.28).

**$^1$H NMR (400 MHz, $CDCl_3$):** $\delta$ 7.36 (d, $J$ = 1.2 Hz, 2H), 7.31 (s, 1H), 3.73 (s, 4H), 3.08 (s, 1H), 2.87 – 2.78 (m, 12H), 2.79 – 2.71 (m, 4H) ppm.

**$^{13}$C NMR (151 MHz, $CDCl_3$):** $\delta$ 169.1 (4C), 167.2 (2C), 138.8 (2C), 131.5 (2C), 130.1, 123.0, 83.2, 77.8, 36.0 (2C), 31.8 (2C), 25.8 (2C), 25.7 (4C) ppm.

**HRMS (ESI$^+$):** exact mass calculated for $[M+Na]^+$ ($C_{24}H_{24}N_2O_8S_2Na$) requires *m/z* 555.0866, found *m/z* 555.0868.

**IR (neat) $\nu_{max}$:** = 3232, 2916, 2664, 1688, 1590, 1420, 1403, 1337, 1302, 1264, 1237, 1197, 1161, 1141, 1041, 910, 880, 869, 805, 771 cm$^{-1}$.

<u>General Procedure 8: Synthesis of NHP Protected Precursor (GP-8)</u>

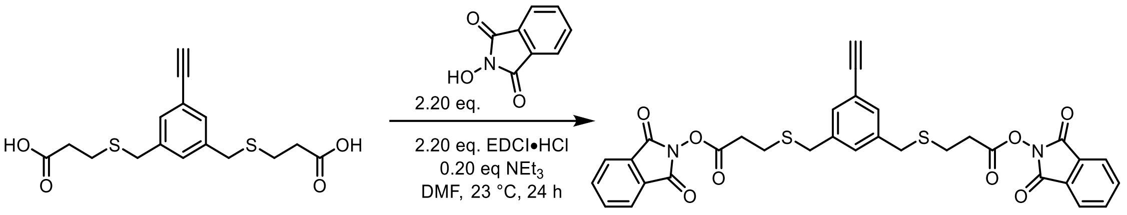

**S7** **S8b**

To flame-dried Schlenk equipped with a stir bar was added diacid **S7** (1.00 eq., 0.21 mmol, 71 mg) and dry DMF (2.0 mL) under an inert atmosphere. Then, *N*-hydroxyphthalimide (2.20 eq., 0.46 mmol, 75 mg), 1-(3-dimethylaminopropyl)-3-ethylcarbodiimide hydrochloride (EDCI•HCl, 2.20 eq., 0.46 mmol, 88 mg) and $NEt_3$ (0.20 eq., 0.04 mmol, 6.0 µL) were added sequentially at 23 °C. The reaction mixture was stirred at 23 °C for 14 h before being quenched by the addition of aq. 1 M HCl (3.0 mL), diluted with ethyl acetate (3.0 mL) and transferred to a separation funnel. The organic phase was separated, and washed with sat. ammonium chloride solution (2 × 3.0 mL), distilled $H_2O$ (2 × 3.0 mL) and brine (2 × 3.0 mL), dried with $MgSO_4$, filtered, and concentrated under reduced pressure. The crude residue was purified by column chromatography.

<u>(Bis(1,3-dioxoisoindolin-2-yl) 3,3'-(((5-ethynyl-1,3-phenylene)bis(methylene))bis(sulfanediyl))dipropionate (**S8b**)</u>

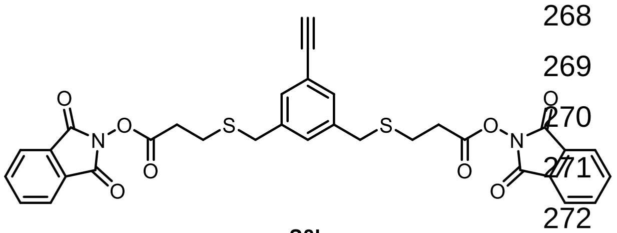

**S8b**

According to general procedure **GP-8**, **S8b** was isolated as a white solid (101 mg, 0.16 mmol, 77%) after purification by column chromatography (heptane/ethyl acetate = 9:1, $R_f$ = 0.27).

**$^1$H NMR (400 MHz, $CDCl_3$):** *δ* 7.93 – 7.84 (m, 4H), 7.83 – 7.75 (m, 4H), 7.39 (d, *J* = 1.5 Hz, 2H), 7.36 (s, 1H), 3.77 (s, 4H), 3.06 (s, 1H), 2.92 (dd, *J* = 11.0, 3.9 Hz, 4H), 2.81 (dd, *J* = 11.2, 4.2 Hz, 4H) ppm.

**$^{13}$C NMR (101 MHz, $CDCl_3$):** *δ* 168.2 (2C), 161.9 (4C), 138.8 (2C), 135.0 (4C), 131.6 (2C), 130.2, 129.0 (4C), 124.2 (4C), 123.0, 83.2, 77.8, 36.1 (2C), 31.9 (2C), 26.0 (2C) ppm.

**HRMS (ESI$^+$):** The exact mass calculated for $[M+Na]^+$ ($C_{32}H_{24}N_2O_8S_2Na$) requires *m/z* 651.0866, found *m/z* 651.0868.

**IR (neat) $\nu_{max}$:** = 3274, 2924, 2853, 1815, 1786, 1738, 1594, 1467, 1450, 1412, 1358, 1325, 1290, 1267, 1241, 1184, 1134, 1078, 1034, 963, 877, 830, 786 $cm^{-1}$.

General Procedure 9: Synthesis of DiSPASO and DiPPASO (GP-9)

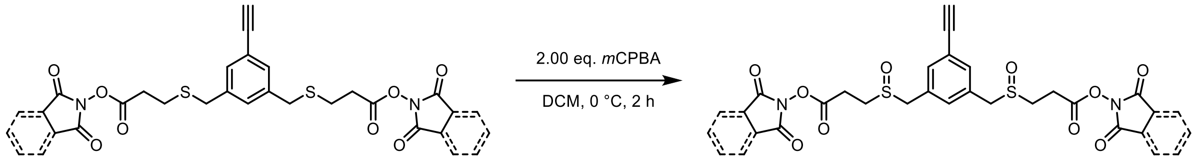

**S8a** or **S8b** **1** or **2**

To flame-dried 4 mL vial equipped with a magnetic stir bar was added the corresponding precursor **S8a** or **S8b** (1.00 eq., 0.05 mmol) and dry DCM (0.5 mL) under an inert atmosphere. Next, *m*CPBA (2.00 eq.) dissolved in 0.5 mL of dry DCM was added dropwise at 0 °C. The reaction mixture was stirred at 0 °C for 2 h followed by the addition of 3.0 mL of sat. aq. $NaHCO_3$ solution and 3.0 mL DCM. The organic phase was separated, and washed with sat. aq. $NaHCO_3$ solution (2 × 3.0 mL), distilled $H_2O$ (3.0 mL) and brine (3.0 mL), dried with $MgSO_4$, filtered, and concentrated under reduced pressure. The crude product was stored in a vial, under an inert atmosphere at –20 °C.

**DiSPASO** (**1**)

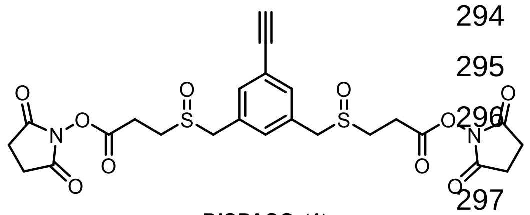

**DiSPASO** (**1**)

According to general procedure **GP-9**, **1** was isolated as a white solid (25.5 mg, 0.05 mmol, 90%).

**$^1$H NMR (600 MHz, $CDCl_3$):** *δ* 7.45 (s, 2H), 7.26 (s, 1H, confirmed by HMBC), 4.0 (s, 4H), 3.20 – 3.03 (m, 7H), 2.93 – 2.87 (m, 2H), 2.85 (s, 8H) ppm.

**$^{13}$C NMR (151 MHz, $CDCl_3$):** *δ* 168.9 (4C), 167.2, 167.2, 133.8, 133.8, 132.2 (2C), 132.2 (2C), 130.9, 130.8, 124.2, 82.0 (observed in HMBC), 79.4, 57.6, 57.6, 44.8 (2C), 25.7 (4C), 24.2, 24.2 ppm.

**HRMS (ESI$^+$):** The exact mass calculated for $[M+Na]^+$ ($C_{24}H_{24}N_2O_{10}S_2Na$) requires *m/z* 587.0765, found *m/z* 587.0765.

**IR (neat) $\nu_{max}$:** = 3270, 2923, 2853, 2359, 1812, 1782, 1735, 1595, 1429, 1367, 1207, 1090, 1046, 894, 744 $cm^{-1}$.

**DiPPASO**(Bis(1,3-dioxoisoindolin-2-yl) 3,3'-((5-ethynyl-1,3-phenylene)bis(methylenesulfinyl))dipro-pionate, **2**)

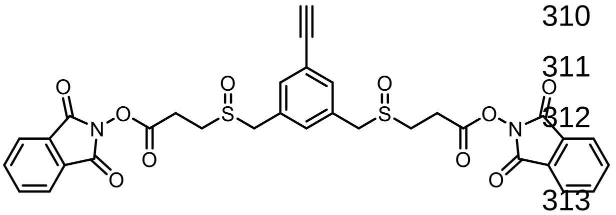

**DiPPASO** (**2**)

According to general procedure **GP-9**, **2** was isolated as a white solid (29.1 mg, 0.04 mmol, 88%).

**$^1$H NMR (400 MHz, CDCl$_3$):** *δ* 7.89 (dt, *J* = 7.1, 3.6 Hz, 4H), 7.84 – 7.76 (m, 4H), 7.47 (d, *J* = 1.1 Hz, 2H), 7.30 (s, 1H), 4.02 (s, 4H), 3.28 – 2.89 (m, 9H) ppm.

**$^{13}$C NMR (176 MHz, CDCl$_3$):** *δ* 168.2, 168.2, 161.7 (4C), 135.1 (4C), 133.9 (2C), 132.3, 132.3, 130.9 (2C), 128.9 (2C), 124.3 (4C), 124.2 (2C), 79.4, 77.4, 57.7 (2C), 57.7 (2C), 45.1 (2C), 45.0 (2C), 24.2 (2C), 24.2 (2C) ppm.

**HRMS (ESI$^+$):** The exact mass calculated for [M+Na]$^+$ ($C_{32}H_{24}N_2O_{10}S_2Na$) requires *m/z* 683.0765, found *m/z* 683.0765.

**IR (neat) $\nu_{max}$:** = 3276, 2924, 2854, 1815. 1787, 1740, 1596, 1467, 1413, 1361, 1267, 1186, 1157, 1136, 1083, 1040, 964, 877, 824, 787 cm$^{-1}$.

#### NMR Spectra

Dimethyl 5-bromoisophthalate (**S2**)

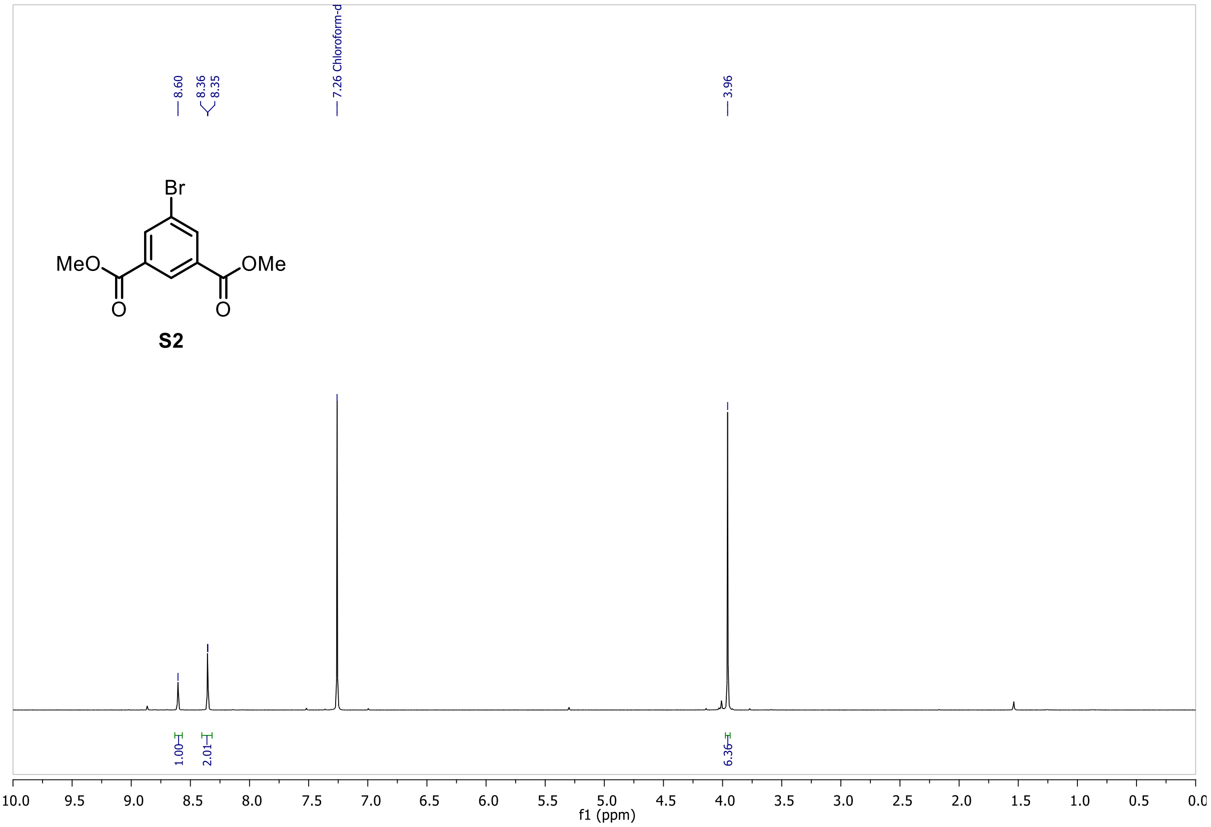

(5-Bromo-1,3-phenylene)dimethanol (**S3**)

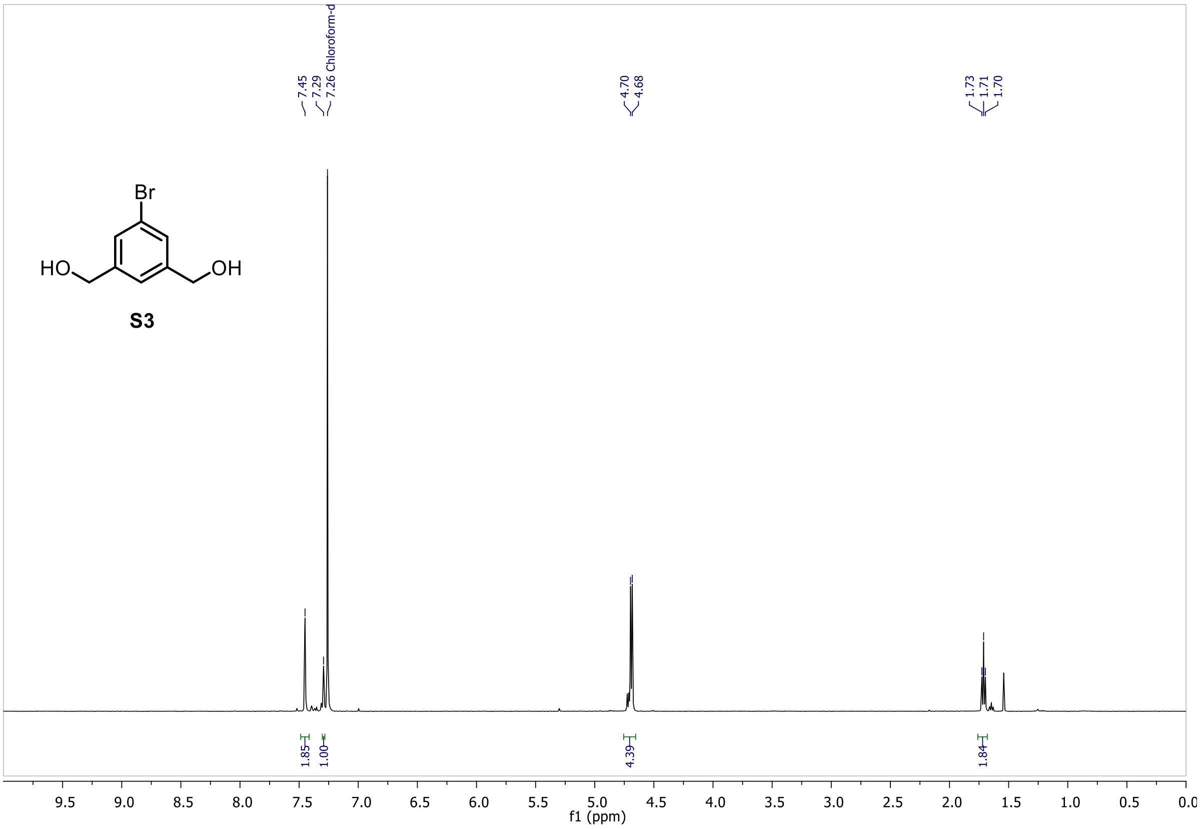

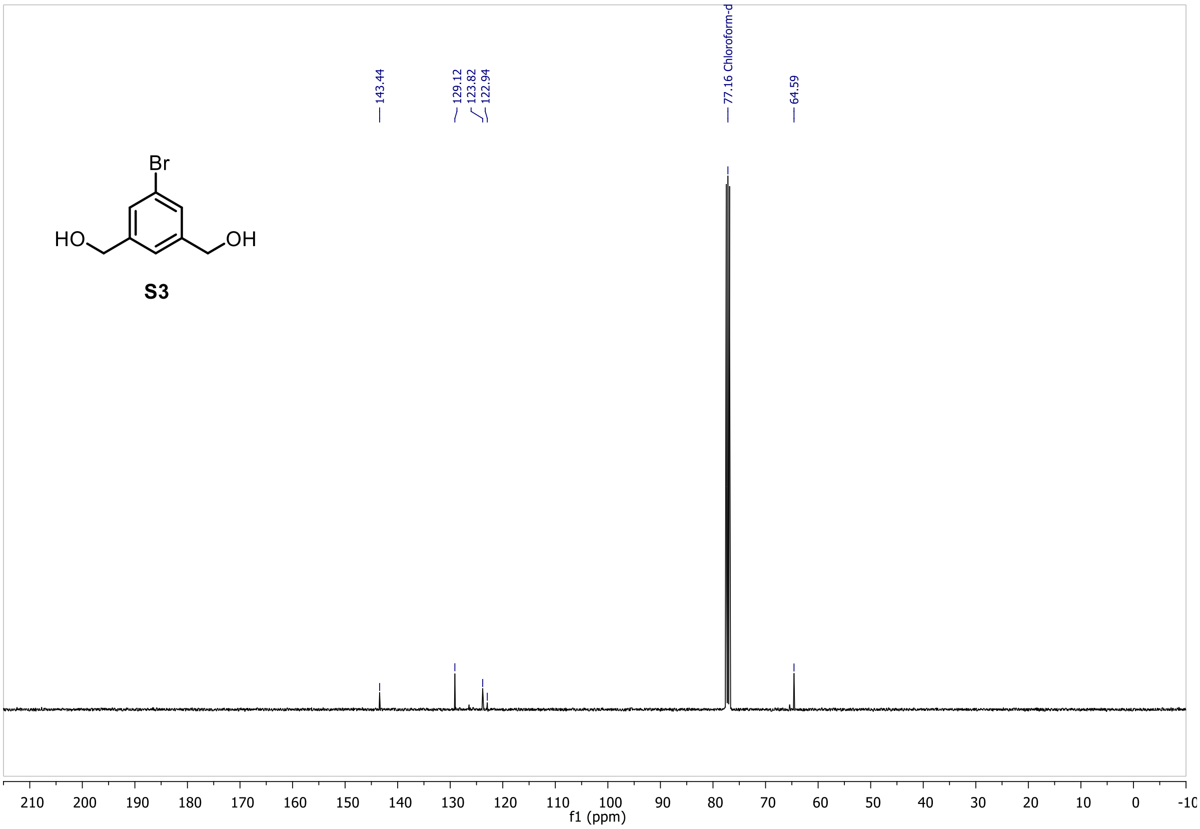

1-Bromo-3,5-bis(bromomethyl)benzene (**S4**)

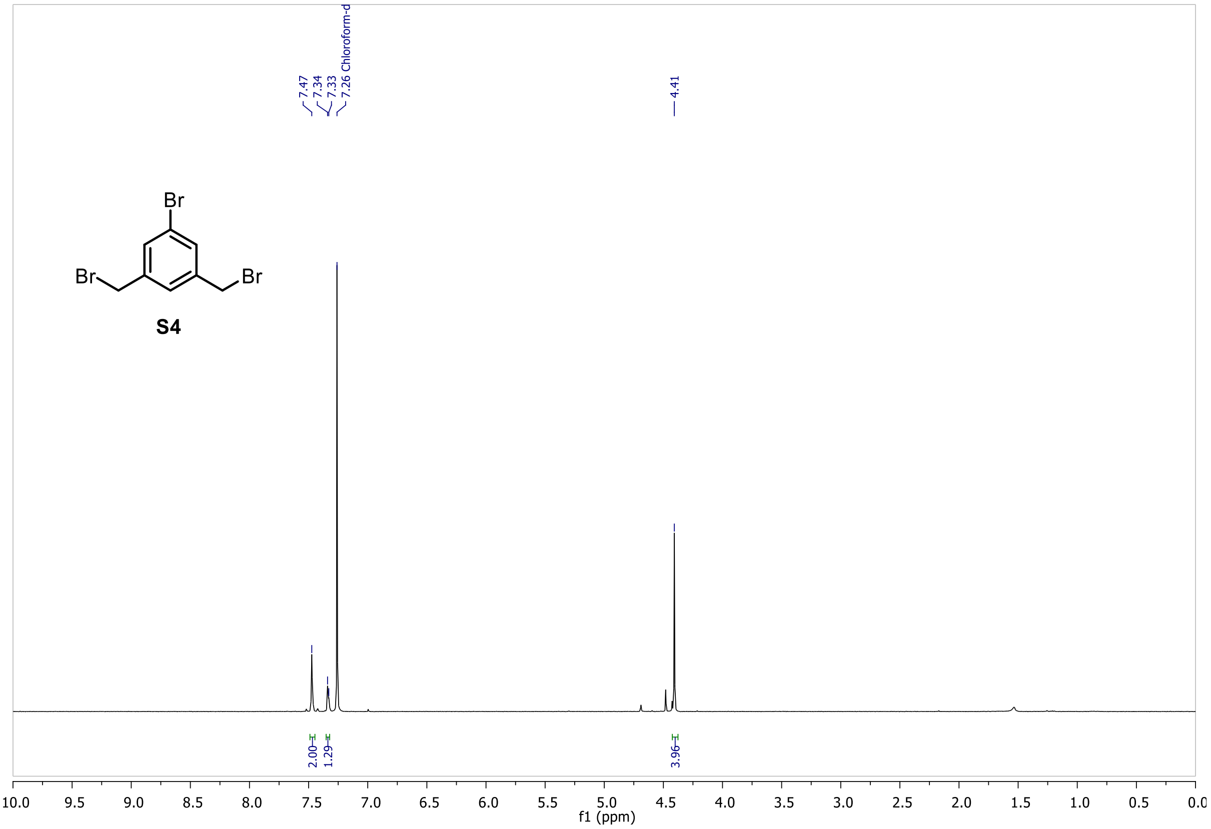

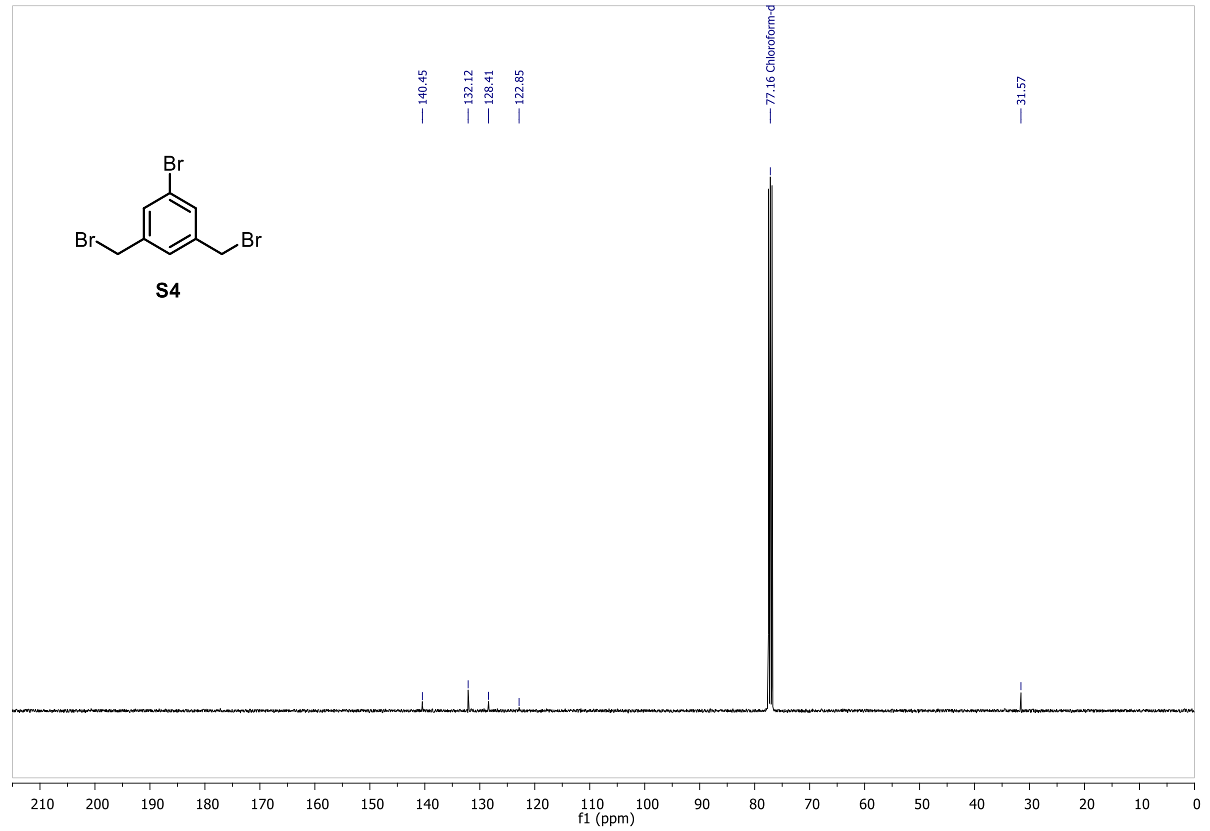

Dimethyl 3,3'-(((5-bromo-1,3-phenylene)bis(methylene))bis(sulfanediyl))dipropionate (**S5**)

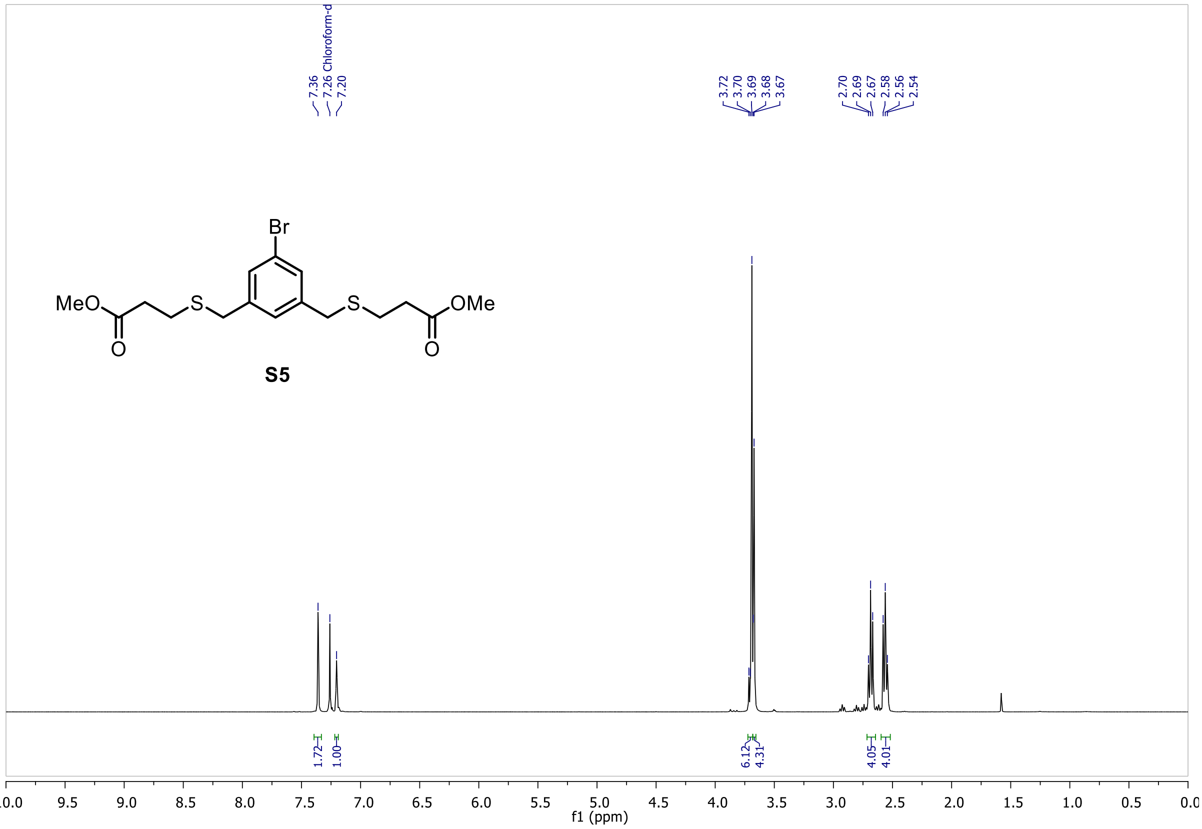

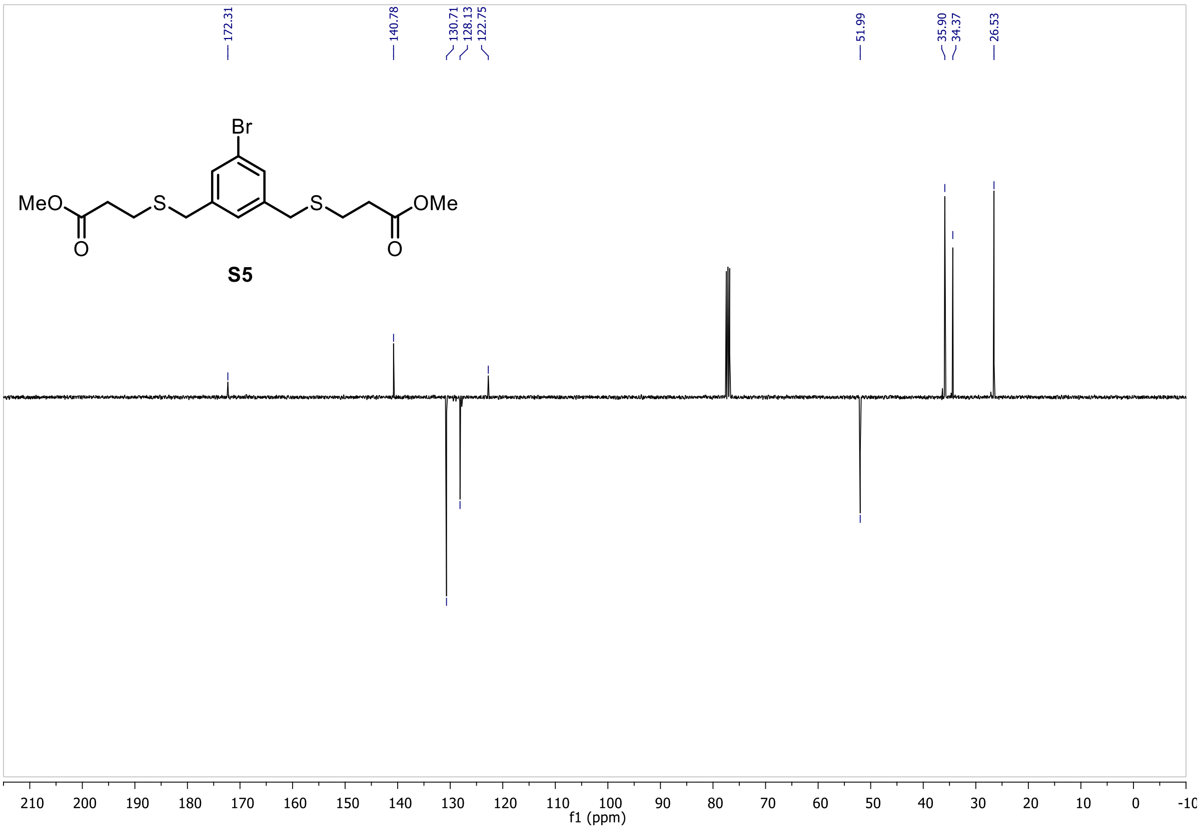

Dimethyl 3,3'-(((5-((trimethylsilyl)ethynyl)-1,3-phenylene)bis(methylene))bis(sulfanediyl))-dipropionate (**S6**)

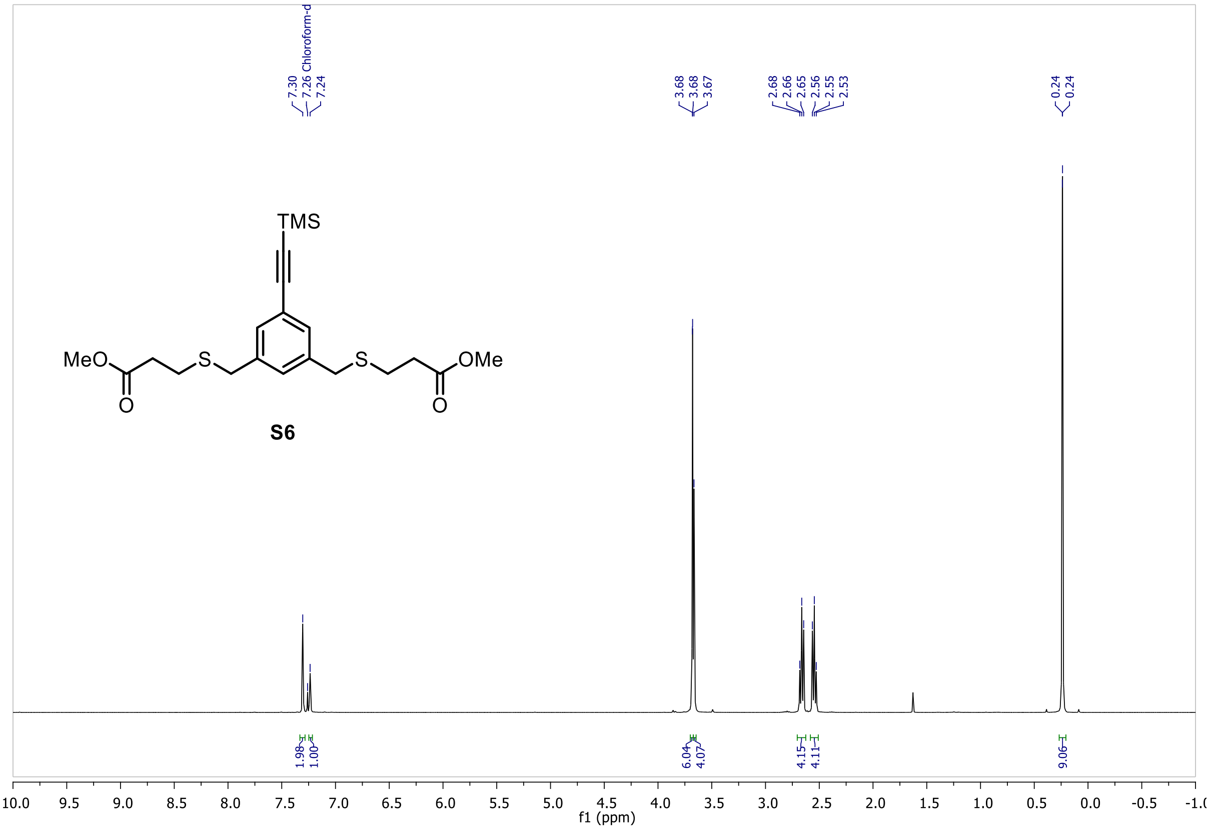

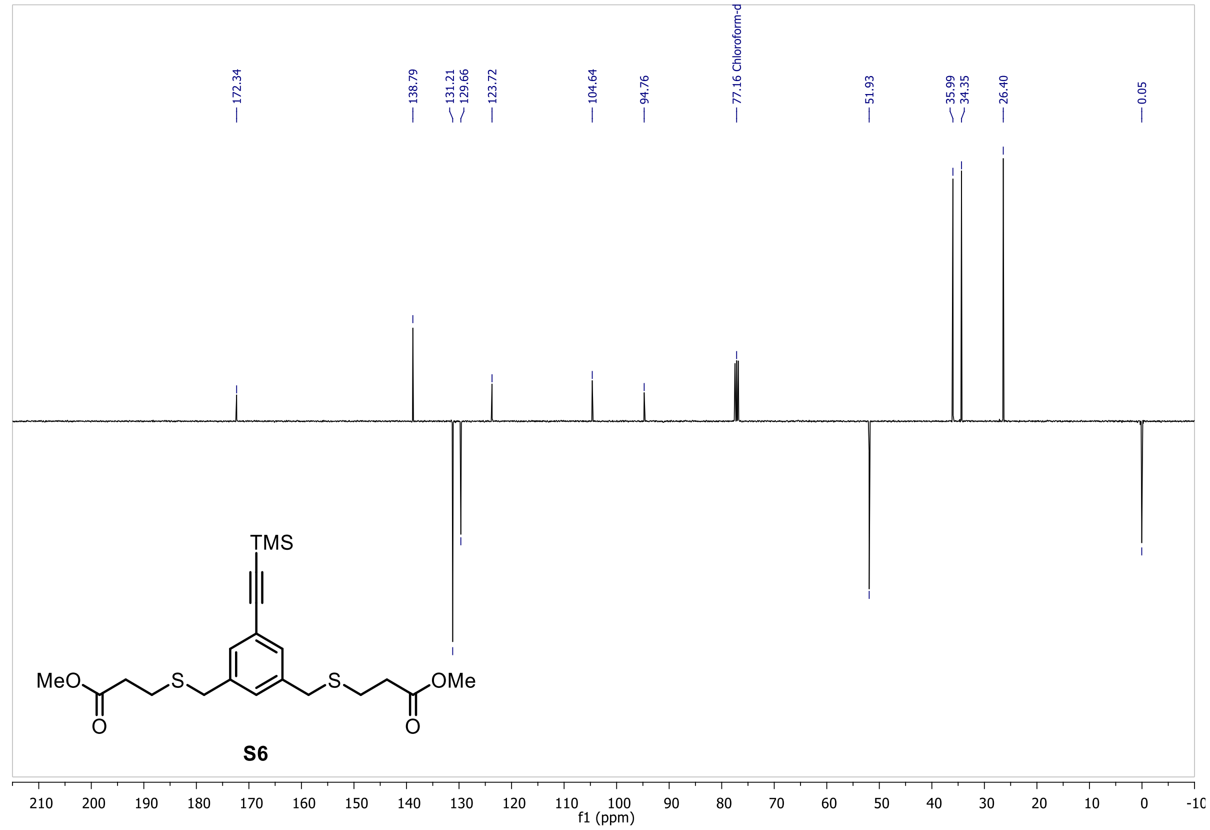

3,3'-(((5-Ethynyl-1,3-phenylene)bis(methylene))bis(sulfanediyl))dipropionic acid (**S7**)

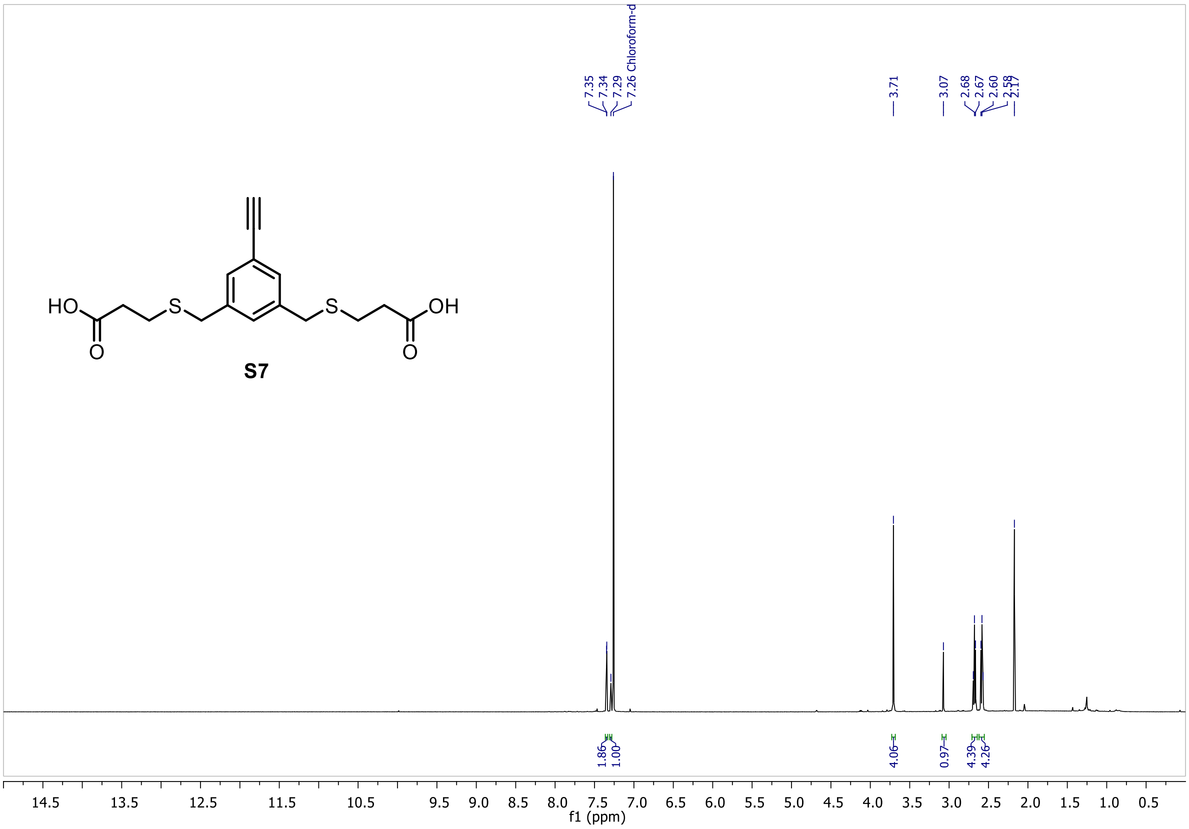

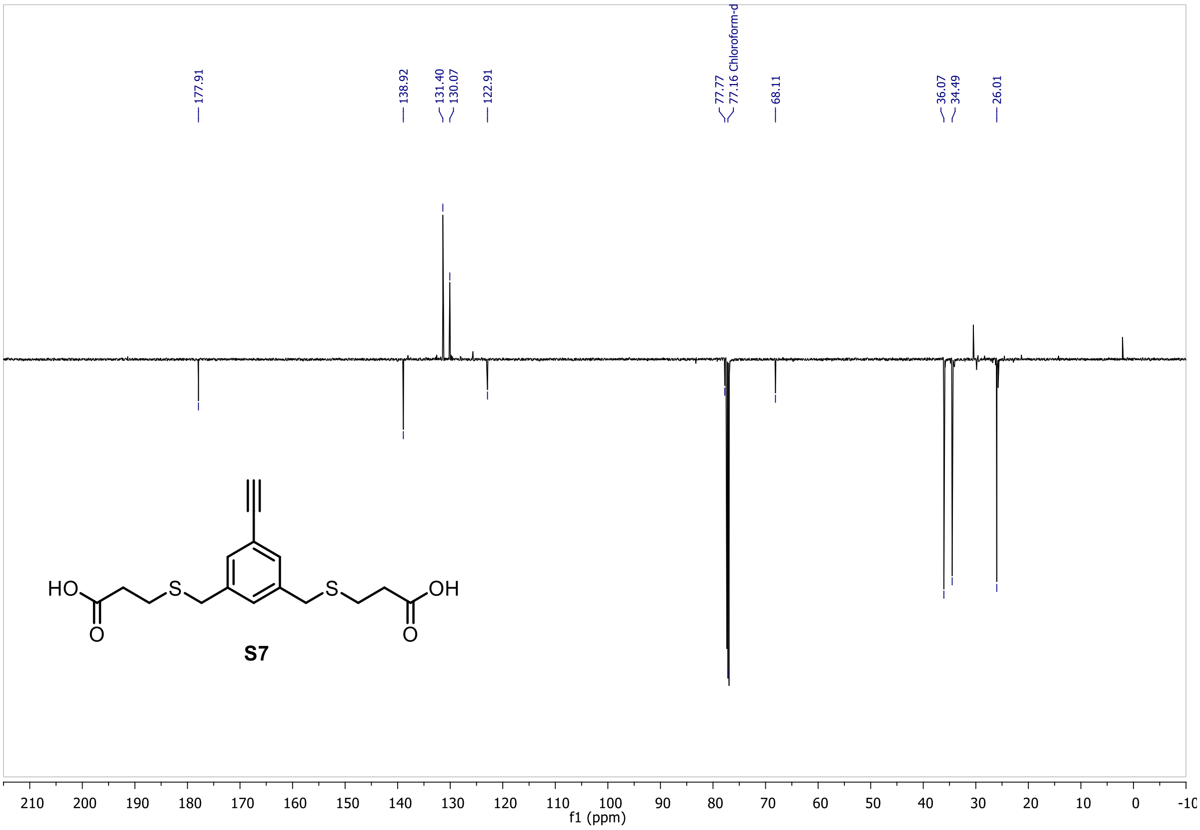

Bis(2,5-dioxopyrrolidin-1-yl)3,3'-(((5-ethynyl-1,3-phenylene)bis(methylene)) bis(sulfanediyl))dipropionate (**S8a**)

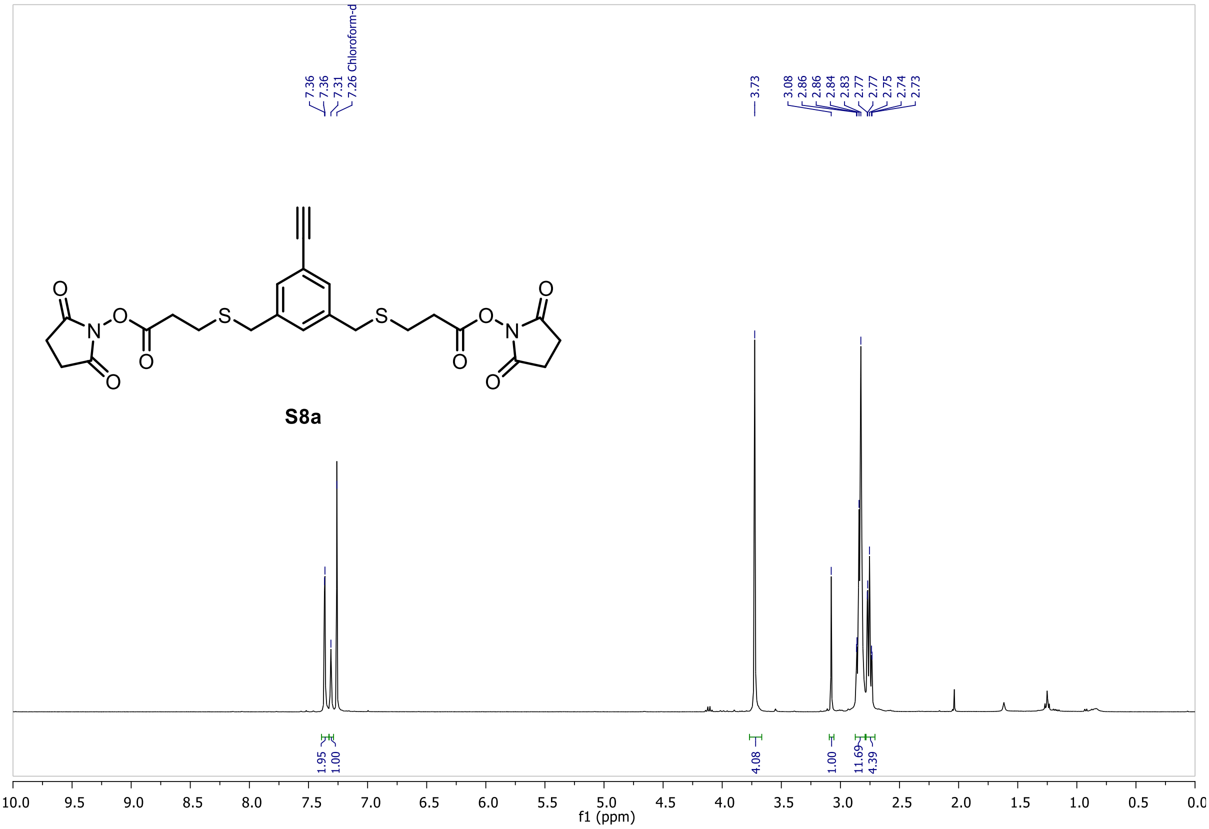

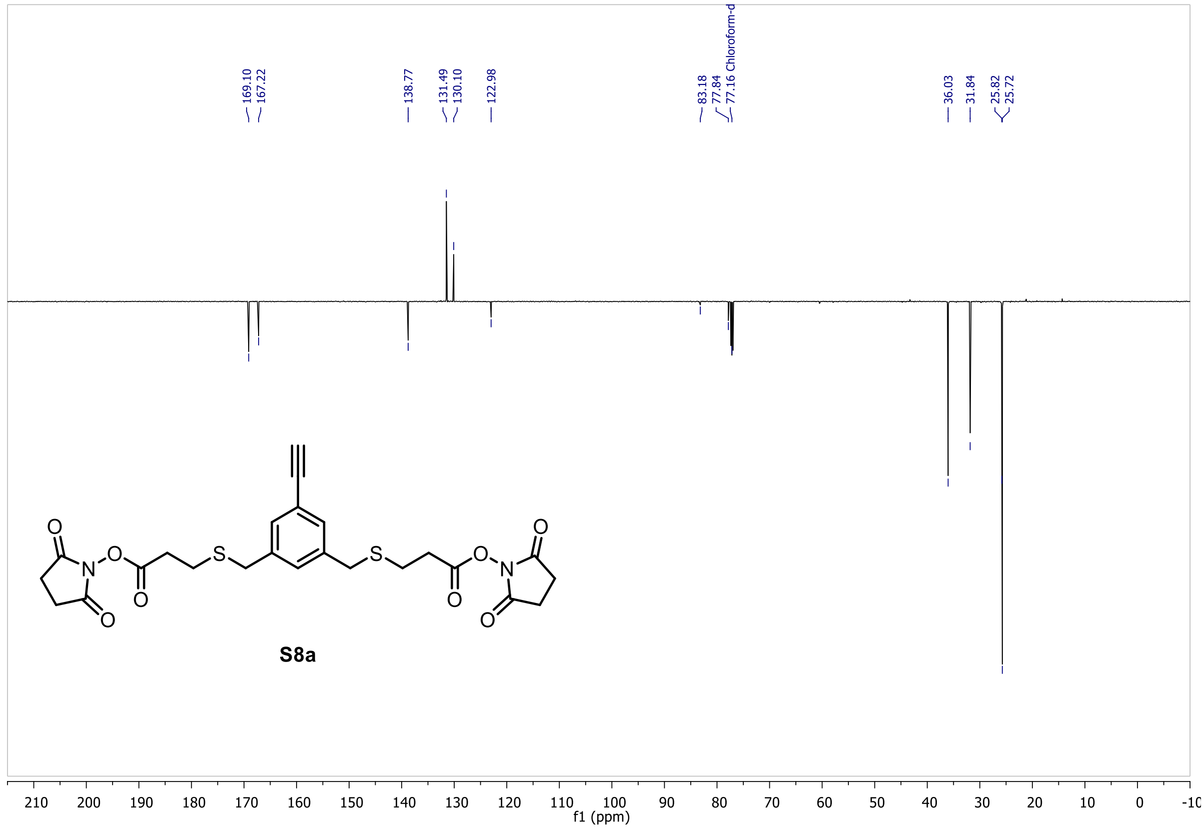

(Bis(1,3-dioxoisoindolin-2-yl) 3,3'-(((5-ethynyl-1,3-phenylene)bis(methylene))bis(sulfanediyl))dipropionate (**S8b**)

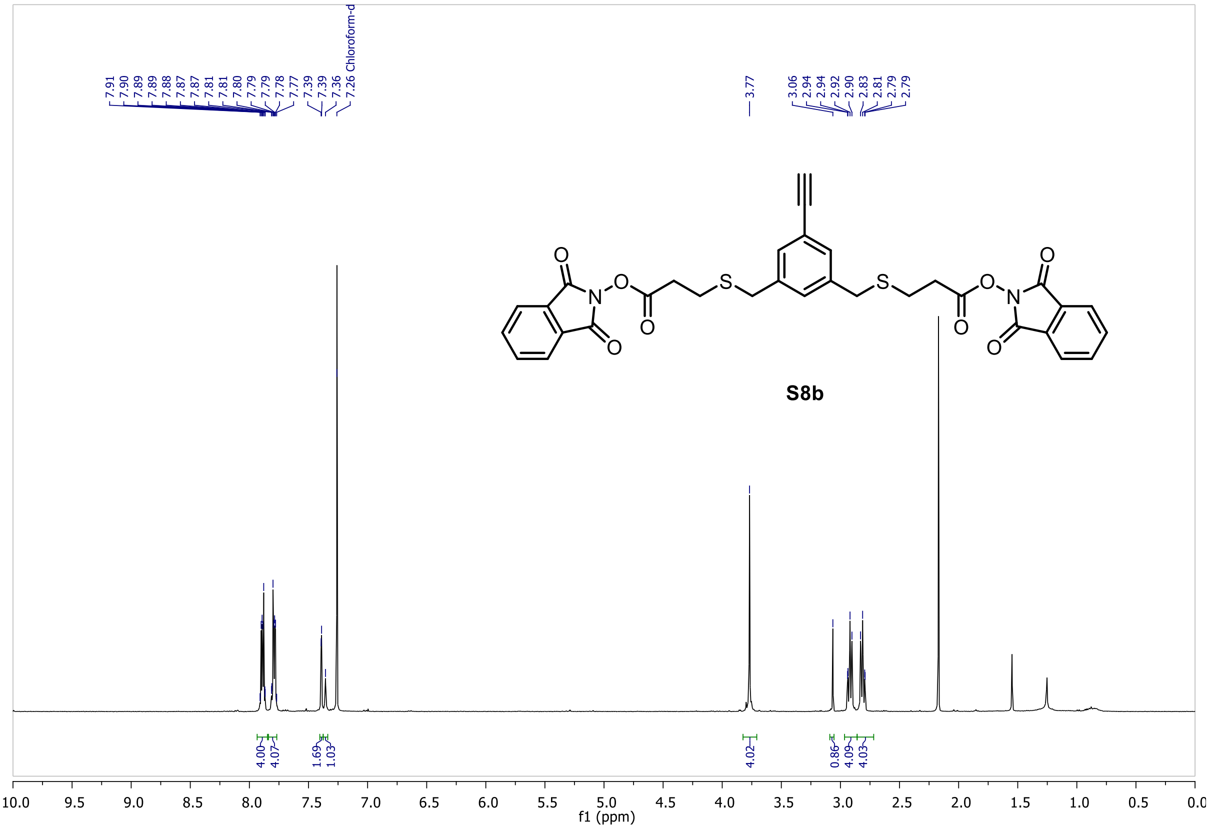

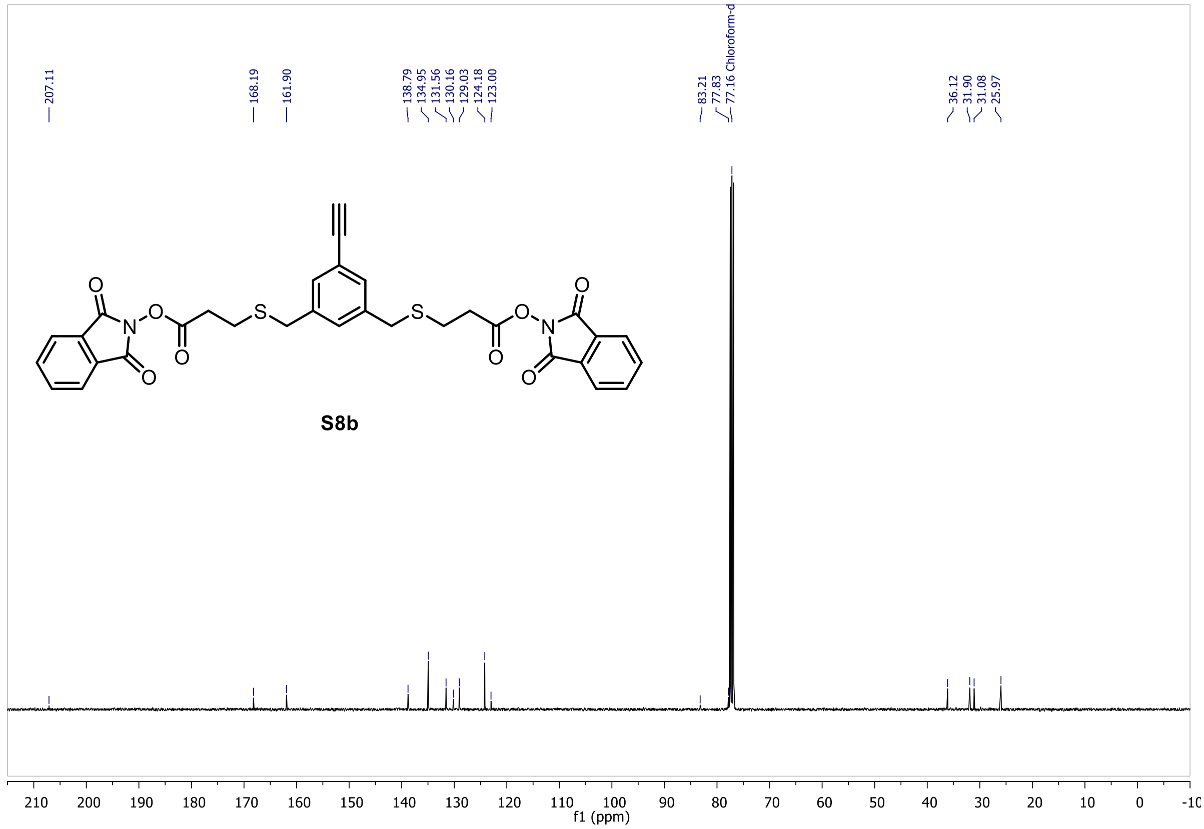

#### DiSPASO (1)

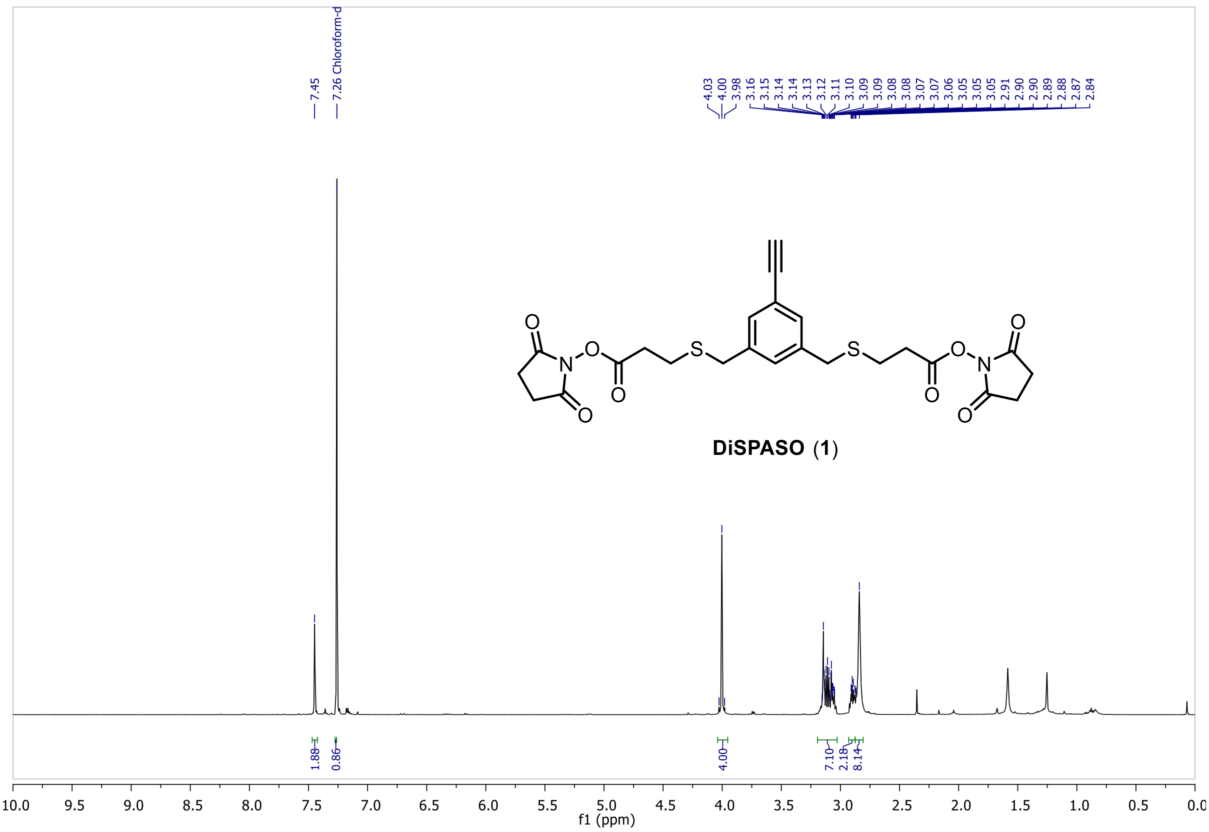
7.45
7.26 Chloroform-d
4.03
4.00
3.98
3.16
3.15
3.14
3.14
3.13
3.12
3.11
3.10
3.09
3.09
3.08
3.08
3.07
3.07
3.06
3.05
3.05
3.05
2.91
2.90
2.90
2.89
2.88
2.87
2.84
DiSPASO (1)
1.88
0.86
4.00
7.10
2.18
8.14
f1 (ppm)

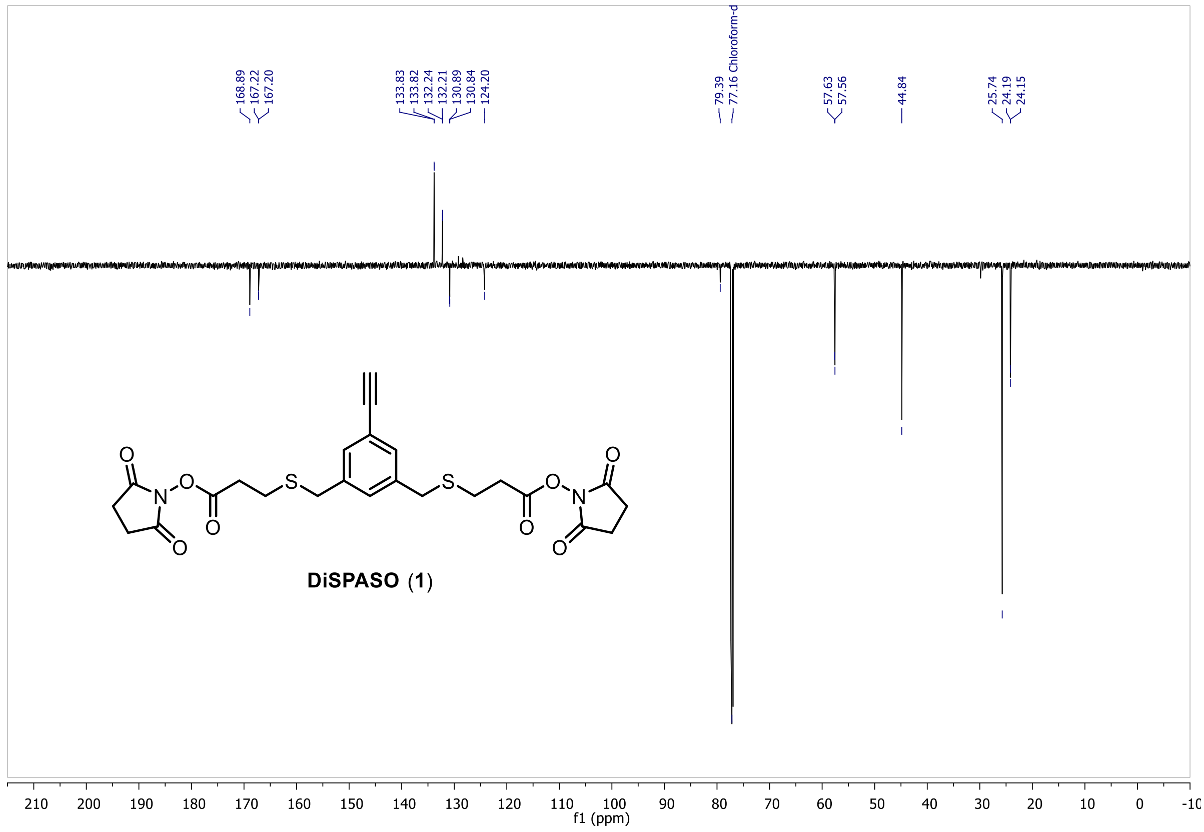
168.89
167.22
167.20
133.83
133.82
132.24
132.21
130.89
130.84
124.20
79.39
77.16 Chloroform-d
57.63
57.56
44.84
25.74
24.19
24.15
DiSPASO (1)
f1 (ppm)

Mass Spectra of DiSPASO (1)

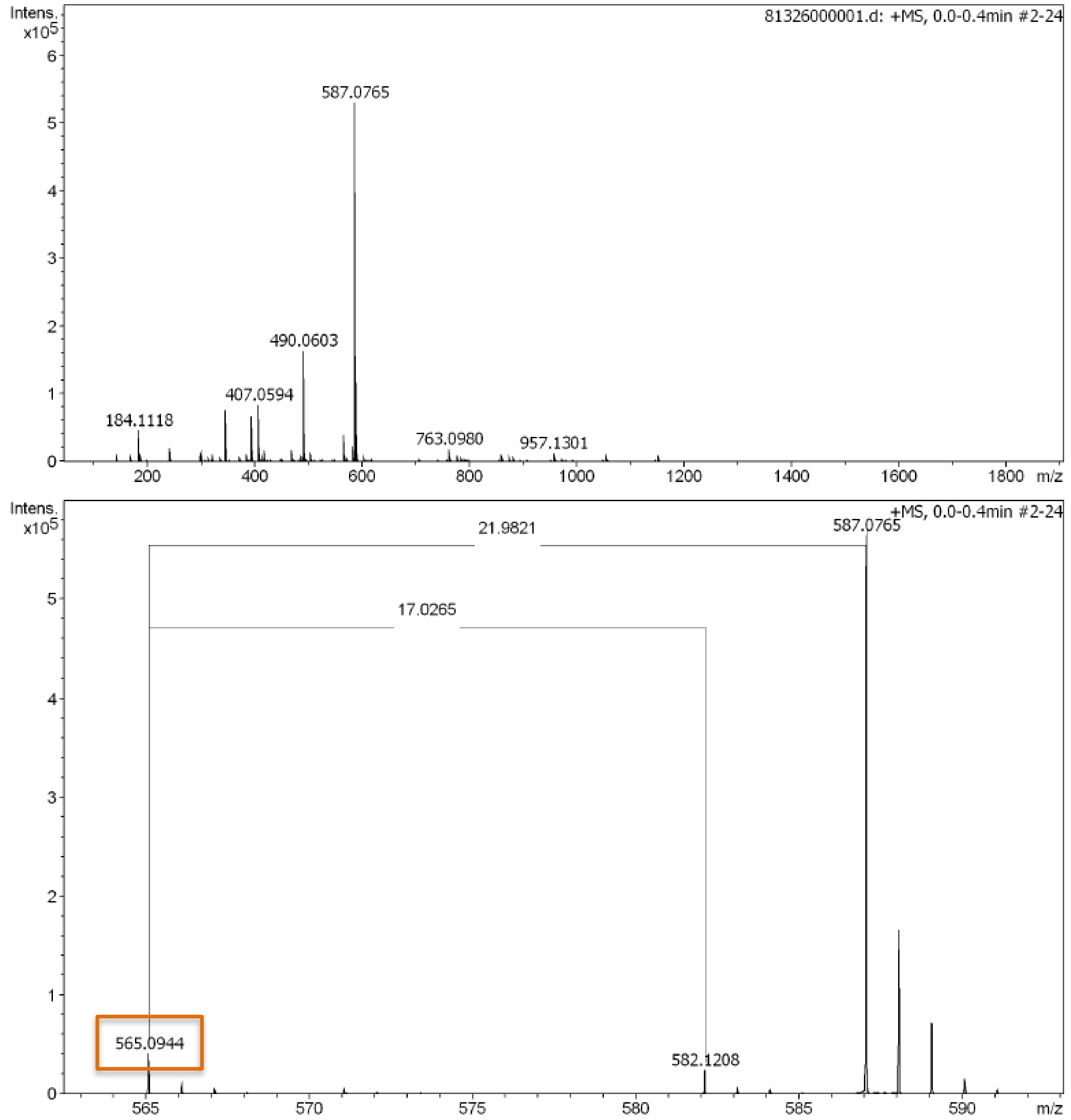

#### DiPPASO (2)

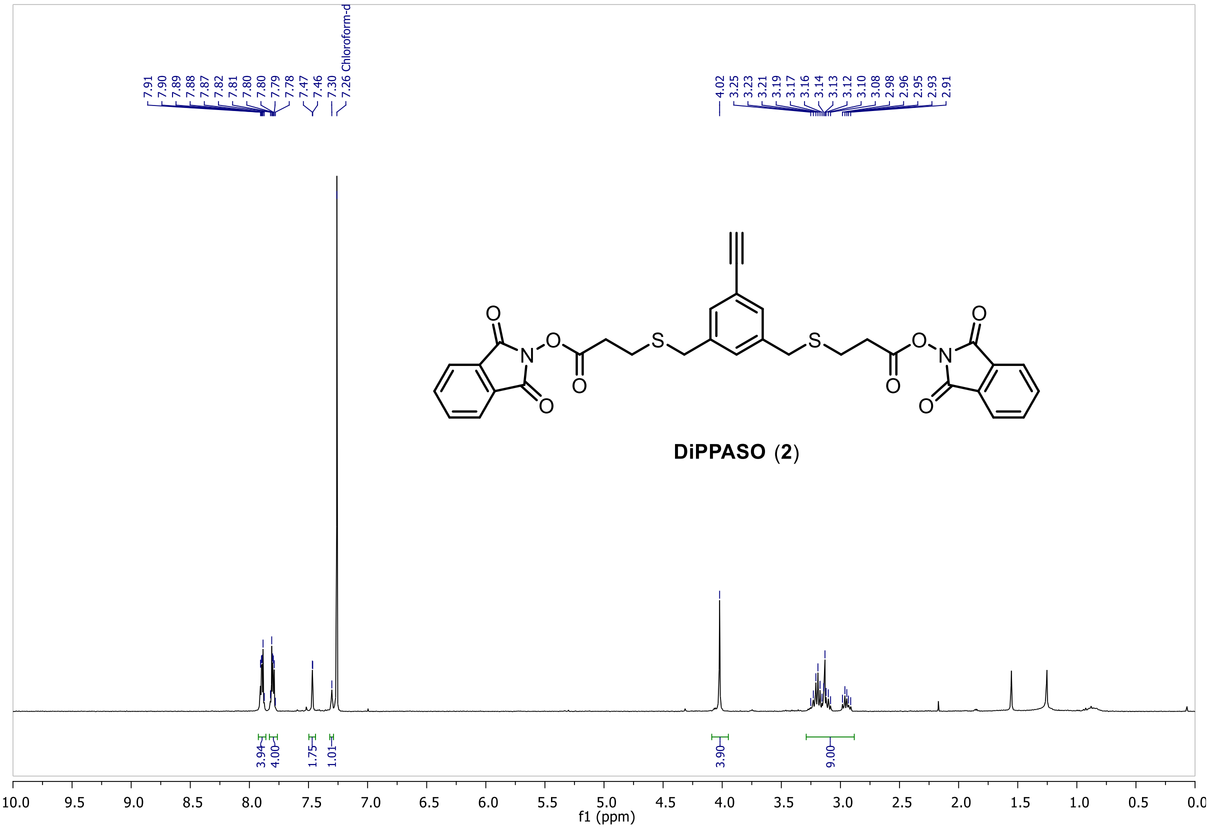
7.91
7.90
7.89
7.88
7.87
7.82
7.81
7.80
7.80
7.79
7.78
7.47
7.46
7.30
7.26 Chloroform-d
4.02
3.25
3.23
3.21
3.19
3.17
3.16
3.14
3.13
3.12
3.10
3.08
2.98
2.96
2.95
2.93
2.91
O
N
S
DiPPASO (2)
3.94
4.00
1.75
1.01
3.90
9.00
10.0
9.5
9.0
8.5
8.0
7.5
7.0
6.5
6.0
5.5
5.0
4.5
4.0
3.5
3.0
2.5
2.0
1.5
1.0
0.5
f1 (ppm)

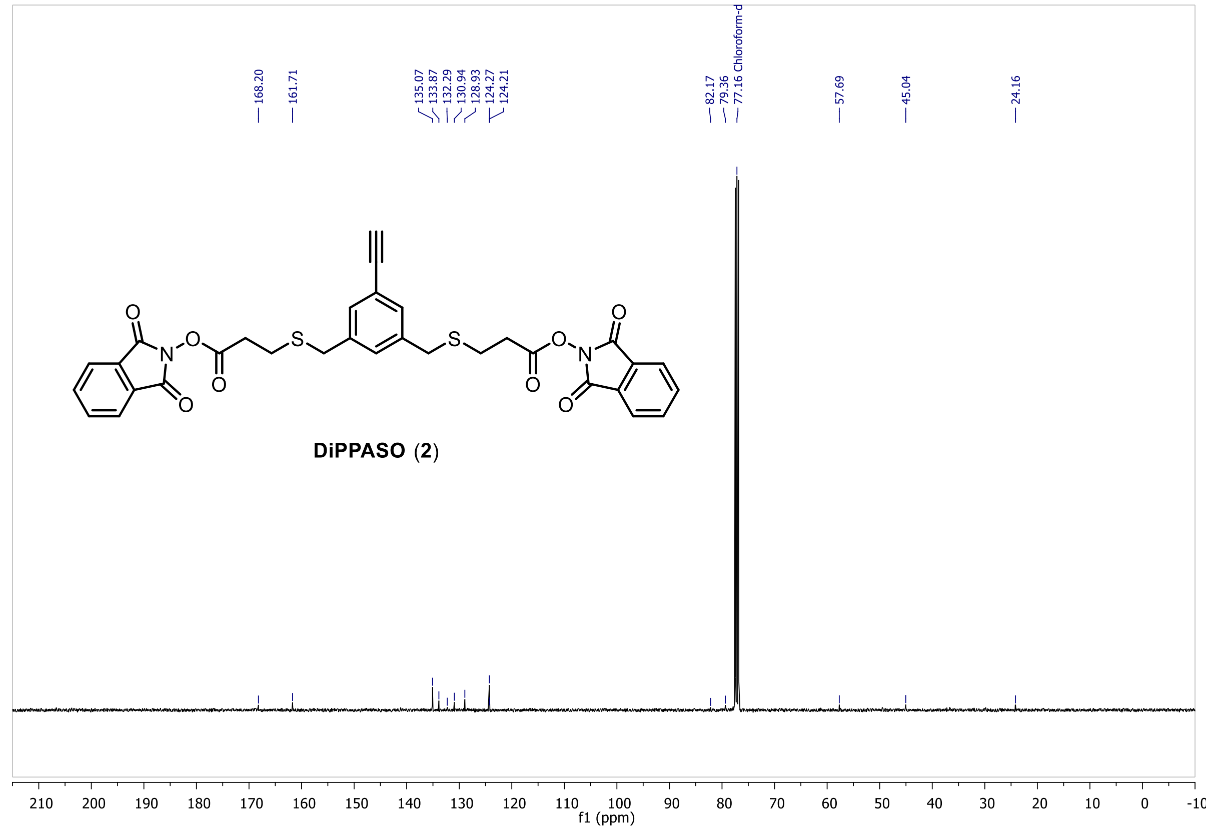
168.20
161.71
135.07
133.87
132.29
130.94
128.93
124.27
124.21
82.17
79.36
77.16 Chloroform-d
57.69
45.04
24.16
O
N
S
DiPPASO (2)
210
200
190
180
170
160
150
140
130
120
110
100
90
80
70
60
50
40
30
20
10
0
f1 (ppm)

### Methods

#### Reagents

*Table S1. Special reagents used for DiSPASO click-reaction, enrichment, and microscopy.*

| Reagent name | Catalogue number | Supplier |
|---|---|---|
| CuAAC Biomolecule Reaction Buffer Kit (BTTAA-based) | CLK-071 | Jena Bioscience |
| MBS magnetic beads Streptavidin | LOT 19-1 | Molecular Biology Services (in-house) |
| Disulfide Azide Agarose | CLK-CSTM, 1238-2 | Click Chemistry Tools |
| Pierce™ High Capacity Streptavidin Agarose | 20359 | Thermo Fisher Scientific |
| Pierce™ Streptavidin Magnetic Beads | 88816 | Thermo Fisher Scientific |
| E. coli Ribosome | P0763S | BioLabs |
| Azide-SS-biotin | BP-22877 | BroadPharm |
| µ-Dish 35 mm, high Grid-500 Glass Bottom | 81168 | Ibidi |
| Tris[(1-Benzyl-1H-1,2,3-Triazol-4-yl)methyl]amin | 678937 | Sigma-Aldrich |
| Cas9 from *S. pyogenes* fused with a Halo-tag | In house | Deng *et al*.[6] |
| Trypsin gold | V5280 | Promega |
| Lysyl endopeptidase (LysC) | 125-05061 | Wako |
| Human HeLa cells | CCL-2 | ATCC |
| Human HEK 293 cells | CRL-1573 | ATCC |

#### Peptide Synthesis

The Peptide Ac-WGGGGR**K**SSAAR-COOH was synthesised on a Liberty Blue peptide synthesiser (CEM) using standard Fmoc chemistry. For each amino acid cycle, a 4-minute coupling with DIC/Oxyma was performed. N-term was acetylated on resin with 5% Acetic anhydride/2,5% DIPEA in DMF for 20min. Peptide was purified on a Phenomenex Luna

C18(2) using a 2-45% in 45 min 0,1%TFA/ACN+0,1%TFA gradient. The identity of the peptide was confirmed using MALDI-MS (4800 MALDI TOF/TOF, Sciex).

#### Crosslinking reaction for Cas9

Cas9-Halo protein was crosslinked using either DSBSO or DiSPASO. All crosslinkers were prepared as stock solutions at a concentration of 20 mM in dry Dimethyl Sulfoxide (DMSO). For the crosslinking reaction, Cas9 was diluted in 50 mM HEPES to achieve a final protein concentration of 1 ug/uL and a crosslinker was added to a final concentration of 0.5 mM. After a 45-minute incubation at room temperature, the reactions were stopped using 100 mM Tris buffer. Each crosslink reaction was prepared in parallel with the same conditions and buffers.

#### Single peptide crosslinking

The peptide Ac-WGGGGR**K**SSAAR-COOH was dissolved in 50 mM HEPES buffer pH 7.3 to a final concentration of 5 mM and crosslinked using 2 mM DiSPASO. The crosslinker was added additionally 5x 2mM over a time course of 2.5h. The reaction was finally quenched with 100mM Tris-buffer pH 8. Click reaction was performed using increasing amounts of picolyl azide: 2, 5, 10, 30mM, 2mM CuSO4, 10mM Tris[(1-Benzyl-1H-1,2,3-Triazol-4-yl)methyl]amin (TBTA) (later replaced by BTTAA for better solubility) and 100mM Sodium Ascorbate (NaAsc). The procedure is described in more detail at point “Click reaction using Azide-S-S-biotin”.

#### In-Solution Digest

For the in-solution digest, ProteaseMax was utilised as a buffer component, prepared as a 1% stock solution in 50 mM Ammonium Bicarbonate (ABC) with a final concentration of 0.05%. Proteins were reduced using 10 mM Dithiothreitol (DTT) followed by incubation for 30 minutes at 50°C, alkylated using 50 mM Iodoacetamide (IAA) for 30min in the dark and finally digested using Trypsin (1:100, enzyme to protein ratio). The mixture was incubated overnight at 37°C to facilitate complete digestion. The digestion process was terminated by the addition of 10% Trifluoroacetic Acid (TFA), adjusting to a final concentration of 0.5%, to ensure the degradation of ProteaseMax.

#### Click reaction using Azide-S-S-biotin

For the click reaction, reagents were prepared from the CuAAC Kit provided by Jena Bioscience, including a 50mM BTTAA stock solution, 1M stock of sodium ascorbate (NaAsc), 100mM CuSO4 stock solution in H2O. A 50mM Azide-S-S-biotin (BroadPharm) stock solution was prepared in dry DMSO. The click reaction utilised crosslinked Cas9 with DiSPASO at a concentration of 1ug/uL, with the reaction environment maintained at pH 7.3. Initially, 2 mM CuSO4 and 10mM BTTAA were mixed, resulting in a blue solution. The Azide-S-S-biotin was then carefully added directly to the crosslinked Cas9 sample. The right amount of Azide was titrated to find an optimal azide concentration of 10mM (Figure S3A) Following a short 5-minute incubation period of azide and sample, the activated Cu(I) solution was added. NaAsc was then introduced to initiate the click reaction. The optimal NaAsc concentration was titrated as well with a resulting optimal concentration of 30mM (Figure S3). The total reaction volume was adjusted with 50 mM HEPES pH 7.3 and incubated for 1 hour at room temperature on a Thermo mixer set at 900 rpm.

#### Crosslinked peptide enrichment

For Azide-S-S-biotin (ASSB) enrichment, the click reaction was performed in HEPES buffer at pH 7.3, using a mixture of 2 mM $CuSO_4$, 10 mM BTTAA, 10 mM ASSB, and 30 mM NaAsc. The reaction mixture was incubated for 1 hour in the dark at room temperature. It was crucial to keep the temperature controlled, NaAsc concentration low, and the reaction shielded from light. The peptides were cleaned and desalted before the enrichment using self-made StageTips according to the procedure published before[7–9] and concentrated in an Eppendorf SpeedVac. For the bead enrichment process, 50 µL of MBS streptavidin beads, with a binding capacity of 1000 pmol free biotin per mg and a concentration of 5 mg/mL, were prepared at a bead-to-volume ratio of 1:5. The beads were washed three times with PBS. The sample, solubilized in PBS, was then added to the beads, and incubated for 1 hour at room temperature, with the supernatant retained for binding checks. Following this, the beads were washed three to four times with PBS to remove non-specific binders. The beads were then resuspended in 20 µL of PBS and eluted with 10 mM TCEP, followed by a 30-minute incubation at room temperature. A second elution was performed with 20 µL of PBS and 10 mM TCEP. The eluted sample was then alkylated with IAA for 45 minutes in the dark at room temperature, followed by another desalting and cleanup step before mass spectrometry analysis.

Additionally, to the MBS beads, Pierce™ High-Capacity Streptavidin Agarose beads with 10mg of biotinylated protein/mL and Pierce™ Streptavidin Magnetic Beads with ~55 µg biotinylated rabbit IgG/mg of beads or ~3500 pmol biotinylated fluorescein/mg of beads binding capacity were tested (Figure S4 A & B).

#### Ribosome crosslinked with DiSPASO

Enriched *E. coli* Ribosome obtained from BioLabs (S P0763S) at a concentration of 13.3 µM, was diluted to 1 mg/mL in 50 mM HEPES pH 7.3, 50 mM KCl, and 10 mM $MgAc_2$. For the crosslinking procedure, 500 µL (500 µg) of this ribosome solution was treated with 1 mM DiSPASO, with the DiSPASO stock being freshly prepared at 30 mM in dry DMSO. The mixture was incubated for 30 minutes on ice, after which an additional 1 mM DiSPASO was added followed by another 30 min of incubation at room temperature. The reaction was then quenched with 1 M Tris to a final concentration of 100 mM Tris and left for 10 minutes. The sample was digested in solution with the addition of 0.05% ProteaseMax to the ribosome solution, followed by mixing and a 10-minute incubation. The reduction of the ribosome solution was carried out using 10 mM DTT followed by 30 min incubation at 50°C. The sample underwent water bath sonication for 3 minutes post-DTT reduction. Iodoacetamide (IAA) was added to a final concentration of 50 mM and incubated for 30 minutes in the dark. LysC was added at a ratio of 1:100 and incubated for 2 hours at 37°C, followed by the addition of trypsin at a ratio of 1:100 and overnight incubation at 37°C. For desalting and cleanup, Sep Pak (50mg, Waters) columns were used to desalt and remove excess crosslinker. The columns were activated with methanol (MeOH), washed, and equilibrated with 0.1% TFA. The sample (pH 3) was then loaded onto the column, washed twice with 0.1% TFA, and eluted with 80% ACN in 0.1% TFA. The ACN was subsequently evaporated using an Eppendorf SpeedVac. For the click reaction, the total volume was adjusted to 500 µL at pH 7.3, with a protein concentration of 1 µg/µL. 2 mM $CuSO_4$ and 10 mM BTTAA were premixed as described above. 10 mM ASSB was added to the clean ribosomal peptides and click reaction initialization was performed by adding 30 mM NaAsc. The sensitivity experiment involved varying concentrations of ribosome and HEK in different ratios, namely 10:100, 2:100, 1:100, 0.5:100, and 0.25:100, with the control sample 10:100 (no enrichment). The ribosome quantities used ranged from 0.25 µg to 10 µg, while the HEK concentration was consistently maintained at 100 µg. Only the 1:100 ratio is shown in Figure S7 to exemplify that the enrichment did not work. Other rations have shown similar results.

<u>Sensitivity experiment using picolyl azide as click reagent</u>

Cas9-Halo was crosslinked (200 µg protein) using 0.25 mM DiSPASO at a total protein concentration of 1ug/ uL. The reaction was carried out for 1h at room temperature and quenched with 100mM Tris buffer pH 8. Directly following the crosslinking, four volumes of pre-cooled acetone were added to the mixture to precipitate the protein. After incubation at -20°C for 1h, the mixture was centrifuged for 10 minutes at 14000 g. The supernatant was carefully removed without disturbing the protein pellet, which was then left to air dry at room temperature for 30 minutes to evaporate the remaining acetone. Over-drying was avoided to ensure the pellet's solubility. For in-solution digestion, the lysis buffer consisted of 8M Urea in 50 mM ABC. To this, 0.05% ProteaseMax was added, and the sample was vortexed and incubated for 10 minutes. The protein sample was reduced using 10mM Dithiothreitol (DTT), for 30min at 50°C following alkylation utilising 50mM Iodoacetamide (IAA) for 30 minutes in the dark. The Urea concentration was reduced to 2 M before protein digestion. Proteolysis was initiated with LysC at a ratio of 1:100 (LysC: protein) for 2 hours at 37°C, followed by Trypsin at a 1:100 ratio and overnight incubation. For desalting and free crosslinker removal, Sep Pak (50mg) columns were utilised. The columns were activated with Methanol and equilibrated with 0.1% TFA. After loading the sample, it was washed twice with 0.1% TFA and eluted with 80% ACN in 0.1% TFA. The clean peptides were dried using an Eppendorf SpeedVac. HeLa Lysate, used as a background, was digested and cleaned in the same way. The sensitivity experiment mixtures contained a constant background of 1mg HeLa digest with increasing spike-in of crosslinked Cas9 samples. The following ratios have been used to determine the sensitivity: 10:100 (1:10), 5:100, 1:100 and 0.5:100. The click reaction setup involved a total volume of 500 µL at pH 7.3, with a total protein concentration of 2 µg/µL. The reaction contained 2 mM CuSO4, 10 mM BTTAA, 5 mM picolyl azide and 100 mM Sodium Ascorbate and was incubated for 1h at room temperature. After clean-up of the clicked and crosslinked peptides an enrichment procedure was performed, involving the use of $TiO_2$-beads.

For the enrichment process, the beads-to-peptide ratio was maintained between 8:1 and 10:1, with a minimum of 10 mg of beads utilised for each condition. Offline columns were assembled with a 10 µm filter and beads weighed directly into them. These beads were first suspended in 200 µl of 50% methanol and centrifuged using a tabletop centrifuge for 1 minute at 2000 g. The beads were then resuspended in 200 µl water, and centrifuged again, followed by two rounds of resuspension in 200 µl 1M glycolic acid solution (1M glycolic acid in a mixture of 70% ACN and 3% TFA) and centrifugation. The beads were finally resuspended in 200 µl of the glycolic acid solution. Concentrated and cleaned samples (200uL final volume), were mixed with an equal volume of the glycolic acid solution and transferred into 1.5 ml tubes. The bead slurry was added to reach a total volume of 700 µl, vortexed, and incubated on a Thermo-Mixer at room temperature for 30 minutes at 1000 rpm. Following incubation, the samples were centrifuged in the offline column. The washing process involved resuspending the beads in various solutions and centrifuging them between each step. Initially, the beads were washed twice with 200 µl glycolic acid solution, followed by two washes with 200 µl 70% ACN/3% TFA, and then two washes with 200 µl 1% ACN/0.1% TFA. For elution, the beads were resuspended in 150 µl 300 mM NH4OH and incubated for 1 minute before centrifugation. The elution step was repeated twice. The eluate was neutralised with 5-7.5 µl concentrated TFA.

In-cell crosslinking of HEK and HeLa cells using DiSPASO

In the conducted experiment, the cell culture was established using Dulbecco's Modified Eagle Medium (DMEM) supplemented with 10% Foetal Bovine Serum (FBS) (50mL) (10270, Fisher Scientific, USA), 1% Penicillin/Streptomycin (5mL) (P0781-100ML, Sigma Aldrich, Israel), and 2mM (1%) L-Glutamine (5mL) (250030-024, Thermo Scientific, Germany). The frozen HEK 293 cells were taken in culture in 6-well plates. Key steps included centrifugation of the cell suspension and resuspension of the cell pellet in fresh media. After cell culture for at least two days (around 80-90% confluency) in the incubator at 37C and 5% $CO_2$, the cells were split, involving the removal of old media, washing with phosphate-buffered-saline (PBS), addition of 0.05% Trypsin-EDTA (25300-054, Thermo Scientific, USA) solution for digestion of surface proteins, preparation of new dishes with new DMEM medium, and addition of cell suspension to new dishes. For in-cell crosslinking, the cells were washed 2x times with PBS and the crosslinker (DiSPASO) was added directly to the cells in each dish. To keep the DMSO concentration below 5%, the crosslinker stock solution was diluted in PBS to a final concentration of 5mM and immediately added to the cells after washing. Keeping the DMSO concentration low ensures that the cells stay intact and avoids crosslinking of broken or damaged cells. After incubation for 30min in an incubator the reaction was quenched with 100mM Tris-buffer for 5min. Afterwards, the cells were washed again with PBS detached with the addition of Trypsin solution and incubated for several minutes at 37 C and 5% CO2. The cells were collected in a new reaction tube, washed again with PBS and centrifuged at 300 g for 2min. Cells were resuspended in a lysis buffer (50 HEPES pH 7.3, 8M Urea and 1% Dodecyl maltoside). After 3 times sonication for 30 sec. (amplitude 80%, 0.5s cycle, UP100H Ultrasonic Processor, Hielscher), the samples were subjected to reduction using Dithiothreitol (DTT) with a final concentration of 10 mM, followed by an incubation period of 30 minutes at 50°C. Subsequently, Iodoacetamide (IAA) was added to reach a final concentration of 50 mM, and the sample was then incubated for 30 minutes at room temperature in the dark. Next, the sample was diluted to a concentration of 2M Urea using 50 mM HEPES pH 7.3. LysC enzyme was added at a ratio of 1:100, and the mixture was incubated at 37°C for 2 hours. Following, Trypsin digest at a ratio of 1:100, and incubation at 37°C overnight. After the digestion, the sample was desalted and cleaned as described earlier.

#### Crosslink enrichment using Disulfide Azide Agarose beads (DAAB)

For the DAAB enrichment, clean crosslinked peptides were diluted to 2ug/uL using HEPES buffer pH 7.3. The click reaction was carried out directly on the beads. Disulfide beads were then added to the sample, followed by a brief incubation period of the beads and sample. The activated Cu(I) solution was added to the sample (2 mM CuSO4 and 10 mM BTTAA), and 30 mM sodium ascorbate (NaAsc) was introduced to initiate the click reaction. The sample was then incubated for 2 hours at room temperature on a rotation wheel to ensure thorough mixing of the beads. Following the incubation period, the beads were washed with PBS, repeating this step three times. Elution was carried out using 10mM TCEP (final concentration), with an incubation period of 1 hour at room temperature, followed by the transfer of the supernatant to a new tube. This elution step was repeated with fresh TCEP, incubating again for 1 hour at room temperature, and the elution was subsequently combined. To facilitate the alkylation of the free disulfide group, 50 mM IAA (final concentration) was added to the elution, and the sample was incubated for 45 minutes at room temperature in the dark.

#### Sample preparation for confocal microscopy of crosslinked HEK cells

The labelling and crosslinking of HEK cells proceeded as follows: Old media was removed, and cells were washed once with PBS. 5mM DiSPASO in PBS was added and incubated for a specified time (0 min (control), 5 min, 15 min, 30 min). After each time point, cells were washed with 1mL PBS and incubated with 200uL 100mM Tris in PBS for 10 minutes. Subsequently, cells were washed twice with 1mL PBS and fixed with 1mL 3.7% formaldehyde solution in PBS for 15 minutes at room temperature. Following fixation, cells were washed twice with 1mL 3% BSA in PBS, and then 1mL PBS-T (0.5% Triton X-100 in PBS) was added, with an incubation period of 20 minutes at room temperature. For the click reaction using the Click-iT Alexa Fluor Picolyl Azide Toolkit, cells were washed twice with PBS. The Alexa Fluor 488 PCA stock solution (950 nominal molecular weight) in 210uL DMSO was prepared to achieve a final concentration of 500uM. A mixture of 2uL of CuSO4 (Compound C) with 8 uL Copper protectant (Compound D) was prepared for a single click reaction. The total volume of the click reaction was adjusted to 500uL, comprising 435 uL of 1x click reaction buffer (Compound B), 5uL of 500 uM Alexa Fluor (final 5uM), 10 uL CuSO4-Copper protectant pre-mix, and 50uL 1x click buffer additive (Compound E), resulting in a final DMSO concentration from the Alexa Fluor of 1%. For wells containing DSBSO crosslinked HEK 293 cells, AlexaFluor 555 alkyne was added instead of AlexaFluor 488. Cells were then incubated for 2 hours at 37°C in the dark. Post-incubation, cells were washed twice with PBS, and DAPI was added for 2 minutes before washing twice with PBS again. Finally, cells were stored in PBS at 4°C until microscopy, with precautions taken to prevent fluorophore bleaching.

#### Confocal microscopy procedure

Images were recorded using a spinning disc confocal scan head (Yokogawa, W1) mounted on Olympus IX3 microscope (Olympus). Multicolour images were acquired using the orca flash 4 camera (Hamamatsu). 40x/0.75 UPLFN (Olympus) was used for images in Figure 7 (main text), 60x/1.2W UPLSAPO (Olympus) and 100x/1.45O UPLXAPO (Olympus) were used for illustrations in Figure S5 left and right respectively. DAPI, AlexaFluor 488 and AlexaFluor 555 were excited using 405nm, 488nm and 561nm lasers respectively. All images were acquired with the same imaging conditions within each experimental group. The exposure times for fluorescence measurements were set to 400 ms for DSBSO (405 nm, 561 nm) and 100 ms (405 nm) and 50 ms (561 nm) for DiSPASO.

#### Relative quantitation of fluorescence signals

For quantifying the fluorescent intensities, we used a custom Fiji-macro[10]. Measurements were performed in 2D, using the central optical slice of each acquisition. Nuclear segmentation was done via the StarDist-plugin (https://github.com/stardist/stardist). Border objects and small fragments were excluded. Intensities for both channels were then measured within the segmented regions.

#### Mass spectrometry

LC-MS/MS analysis was performed using an Orbitrap Exploris 480 or Orbitrap Eclipse Tribrid mass spectrometer with Field asymmetric ion mobility spectrometry (FAIMS) interface (Thermo Fisher Scientific, Waltham, Massachusetts, United States) coupled with a Dionex UltiMate 3000 HPLC system (Thermo Fisher Scientific, Waltham, Massachusetts, United States). A trap column PepMap C18 (5 mm × 300 μm ID, 5 μm particles, 100 Å pore size) (Thermo Fisher Scientific, Waltham, Massachusetts, United States) and an analytical column PepMap C18 (500 mm × 75 μm ID, 2 μm, 100 Å) (Thermo Fisher Scientific, Waltham, Massachusetts, United States) were employed for separation. The column temperature was set to 50 °C. Sample loading was performed using 0.1% trifluoroacetic acid in water with a flow rate of 50 uL/min for 3 min. Mobile phases used for separation were as follows: (A) 0.1% formic acid (FA) in water; (B) 80% acetonitrile, 0.1% FA in water. Peptides were eluted using a flow rate of 230 nL/min, with the following gradient: from 2% to 45% phase B in 90 min, from 45% to 95% phase B in 1 min, followed by a washing step at 95% for 6 min, and re-equilibration of the column. The gradient was altered over time for optimisation purposes. The gradient stated here was the best performing in our hands.

FAIMS separation was performed with the following settings: inner and outer electrode temperatures were 100 °C, FAIMS carrier gas flow was 4.6 L/min, compensation voltages (CVs) of −50, −60, and −70 V were used in a stepwise mode during the analysis. The mass spectrometer was operated in a data-dependent mode with cycle time 2s, using the following full scan parameters: *m/z* range 375-1500, nominal resolution of 120 000, with a target of 250% charges for the automated gain control (AGC), and automated selection of maximum injection time. For higher-energy collision-induced dissociation (HCD) MS/MS scans, a stepped normalised collision energy (NCE) of (25%; 27%; 32%) for DiSPASO and 21%; 27%; 32%) for DSBSO, the resolution was 30 000. Precursor ions were isolated in a 1.4 Th window with no offset and accumulated for a maximum of 54 ms or until the AGC target of 100% was reached. Precursors of charge states from 3+ to 6+ were scheduled for fragmentation. Previously targeted precursors were dynamically excluded from fragmentation for 25 seconds. The sample load was typically 500 ng and 200 ng. Detailed parameters can be found in each raw file under the instrument method section.

#### Data analysis

Raw files were analysed using Thermo Proteome Discoverer (v. 3.1.0.638). Searches were performed against the Cas9 sequence (Uniprot ID: Q99ZW2) plus a Crapome database (downloaded from https://www.thegpm.org/crap/). For in-cell crosslinking searches, fasta files were created for each experiment series separately by searching against the full Human database (Uniprot ID UP000005640, last update 04.08.2022, 20528 sequences). Identified proteins with less than 3 peptide spectrum matches (PSM) were filtered out. For the ribosome samples, the *E. coli* K12 database (Uniprot IDUP000000625, last update 24.04.2023, 5296 sequences) was used to create a specific database for crosslink searches.

Linear peptides were identified using MS Amanda search engine (v. 3.0.20.558)[10] and the crosslinked peptides were identified using MS Annika (v. 2.0)[11,12]. The search workflow included a recalibration step for each file, followed by a first search using MS Amanda to identify linear peptides and monolinks. Subsequently, spectra with highly confident identifications of a linear peptide were filtered out and not considered for the cross-link search. Finally, a crosslink search was performed using MS Annika. The workflow used in Proteome Discoverer is shown in Figure S6. Search parameters for linear and crosslink searches can be found in Supplementary Table 1 and Table 2. The FDR was estimated using the MS Annika validator node with 1% FDR (high confidence) for all single peptide and Cas9 crosslinking data and 5% (high and medium confidence) for all in-cell crosslinking data on CSM and residue pair levels. The FDR calculation is based on a target-decoy approach[11].

For data filtering and visualisation Python 3.9.7 was used with the following packages: pandas[13], numpy[13,14], matplotlib[15] (pyplot, venn), seaborn[16], scipy and bioinfokit[17,18].

#### Software adjustments

We have updated MS Annika, our cross-linking search engine, to better accommodate the fragmentation behaviour of DiSPASO. By adjusting the "Additional Crosslink Doublet Distances" parameter, multiple doublets can now be searched in a single run, provided they share the same light fragment as specified in the crosslinker definition in Proteome Discoverer. For instance, in one search run, it is feasible to explore doublets such as alkene - thiol, alkene - sulfenic acid, alkene -ETFP, alkene – ETHMP, and alkene - full crosslinker mass by designating alkene - thiol as the default doublet and adding the remaining doublet distances in the MS Annika settings. However, including a smaller doublet in the same run is not feasible as it would alter the role of the alkene fragment due to its lower mass. The additional doublet distances are determined by subtracting the monoisotopic mass of the lighter fragment from that of the heavier one. Additionally, MS Annika now considers all potential crosslinker fragments as peptide modifications during search, further enhancing crosslink identification. This entails specifying all expected fragments as neutral losses in the crosslinker definition in Proteome Discoverer, allowing MS Annika to calculate corresponding theoretical ions for precise identification during the database search.

$$Doublet\ distance\ = Mass(fragment_{heavy}) - Mass(fragment_{light})$$

$$Doublet\ distance_{A\,-\,SA} = Mass(sulfenic\ acid) - Mass(alkene)$$

$$Doublet\ distance_{A\,-\,SA} = 119.0041\ - 69.02146$$

$$Doublet\ distance_{A\,-\,SA} = 49.98264$$

*Equation 1. Exemplary calculation of the doublet distance for the doublet alkene - sulfenic acid (A - SA). The subtraction of the mass of the lighter fragment from the mass of the heavier fragment gives the doublet distance. For the case of the A - SA doublet, subtracting the mass of the alkene fragment from the mass of the sulfenic acid fragment yields a doublet distance of approximately 49.98 (Da).*

##### Surface area plot creation

To determine the lipophilicity and other physicochemical properties of crosslinker compounds, a Python script utilizing RDKit and pandas libraries was used. First, the cLogP (partition coefficient) of the compounds was calculated. The cLogP value quantifies a compound's tendency to partition between an organic solvent (typically octanol) and water, reflecting its lipophilicity. This was achieved by defining a function calculate_clogp that utilizes RDKit's MolFromSmiles and MolLogP functions to calculate the cLogP value for each compound represented by its SMILES notation. Similarly, the topological polar surface area (tPSA) was calculated using the function calculate_tpsa, which utilizes RDKit's CalcTPSA function to compute the tPSA based on the molecule's topology. Additionally, a scatter plot was generated to visualize the relationship between cLogP and tPSA values for all crosslinker compounds. The plot provides insights into the correlation between the lipophilicity and polar surface area of the compounds, which are crucial factors in drug design and membrane permeability prediction. Furthermore, the partition coefficient (P) highlights its significance in understanding a compound's hydrophobicity.

#### Supplemental figures

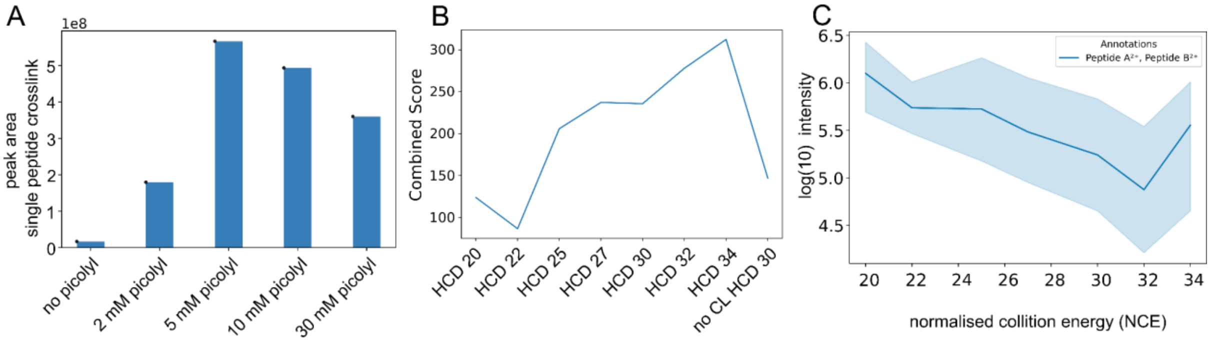

*Figure S1: Single peptide evaluation of DiSPASO. A: Titration experiment of a single Peptide crosslinked with DiSPASO. The picolyl azide concentrations were set in increasing order to 2 mM, 5 mM, 10 mM and 30 mM. The non-crosslinked peptide was used as a control. Increasing the amount of picolyl azide also increased the peak area of the crosslinked product with a maximum of 5 mM compound. B: Titration series of high-performance collision energies concerning the combined score of peptides a and b within the crosslink. The score of a crosslinked peptide increases with high energies with a maximum at HCD 34. C: Log(10) intensities of the 32 Da fragment doublet pair after fragmentation of DiSPASO crosslinked single peptide. The intensity of the doublet decreases with higher energies but rises again with HCD 34.*

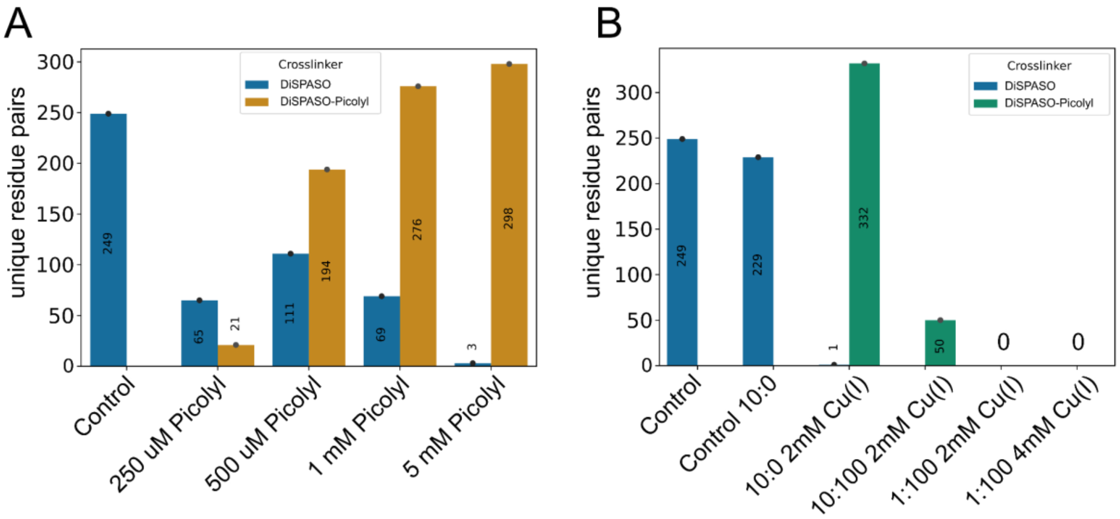

*Figure S2: Evaluation of picolyl concentration for click reaction and enrichment sensitivity in Cas9 only and with HeLa background. A: Picolyl azide concentration titration using crosslinked Cas9 with DiSPASO. Non-enriched DiSPASO crosslinked peptides were used as the control sample (blue), whereas the titration series with increasing amounts of picolyl azide is shown in yellow. The maximum number of identified crosslinked peptides could be achieved with 5 mM picolyl azide as already shown in Figure S1A with single peptide crosslinking. B: Challenging the picolyl azide enrichment with HeLa background. When Cas9 crosslinked with DiSPASO is spiked into HeLa background (10:100 and 1:100) crosslinked peptides can't be enriched sufficiently anymore. The numbers drop from a 10:0 ratio (not spiked-in) to 10:100 with a 100ug HEK lysate background. Crosslinks are not identifiable anymore at in-cell crosslinking levels of 1:100.*

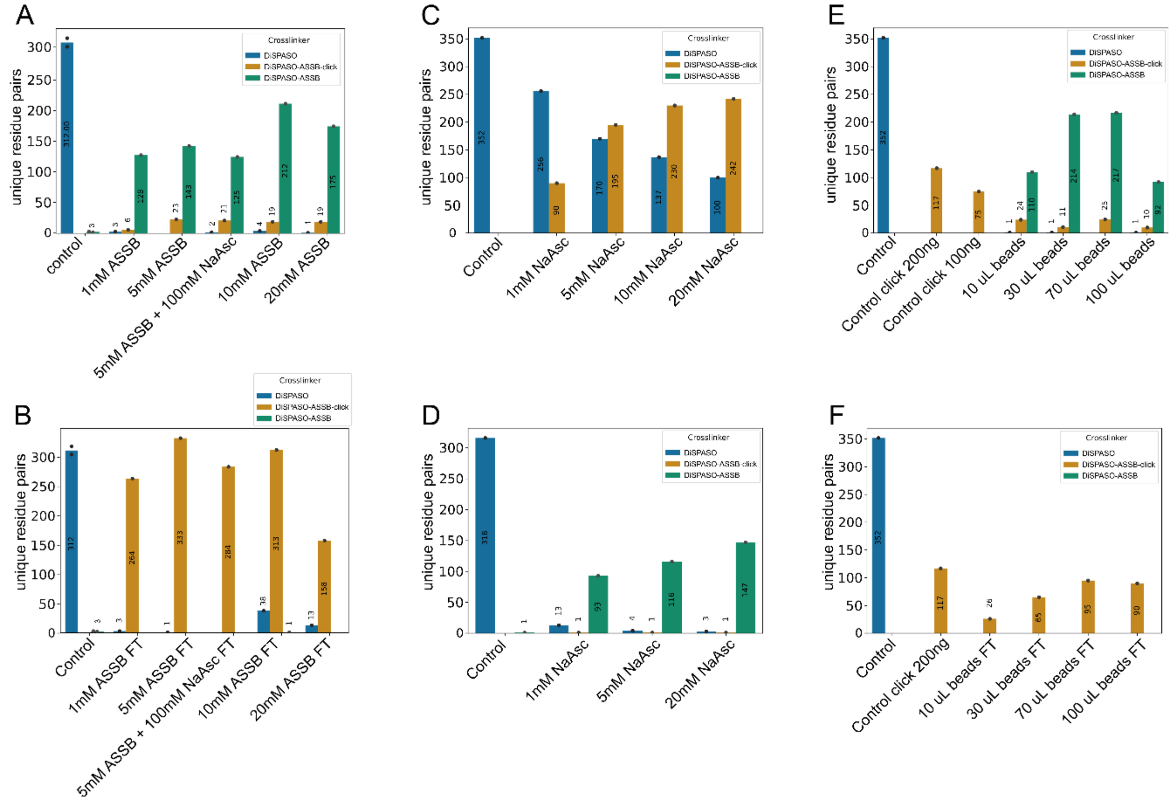

*Figure S3: Optimization of Azide-S-S-biotin, sodium ascorbate and bead amount to achieve optimal click and enrichment performance. A: Titration of the optimal Azide-S-S-biotin (ASSB) amount to*

*achieve high click reaction performance. The concentration was set to 1 mM, 5 mM, 5 mM + 100 mM sodium ascorbate, 10 mM and 20 mM. After 20 mM the maximum solubility in the reaction mix is reached. At 10 mM ASSB, the total number of identified residue pairs reaches its maximum. B: The flowthrough of this experiment was also assessed to check for potential losses during the workflow. The overall loss seems to be independent of the ASSB concentration and is in general high. C: Titration of the optimal sodium ascorbate (NaAsc) concentration for click reaction. The concentration was set to 1 mM, 5 mM, 10 mM and 20 mM. After 10 mM NaAsc, the enrichment performance reaches almost its plateau. The click reaction efficiency can be estimated by following the decrease of non-clicked DiSPASO crosslinked peptides (blue) to the increase of click product after click reaction (yellow). D: Performance of biotin-streptavidin bead enrichment after ASSB optimization. The optimal enrichment performance could be achieved with 10 mM ASSB and 20 mM Sodium ascorbate. The NaAsc concentration was set to 30 mM for further experiment to ensure high click performance while reducing potential side reaction of the copper-based click reaction. E: To reduce the loss of the bead-based enrichment strategy a titration of the right bead amount was performed. The volume of the beads was set to 10 uL, 30 uL, 70 uL and 100 uL of MBS bead slurry. Most residue pairs could be identified with 70 uL beads for enrichment. F: The flowthrough after bead enrichment was tested again to assess the loss after bead optimization, the loss could be reduced by half but is still present even after using 100 uL of beads.*

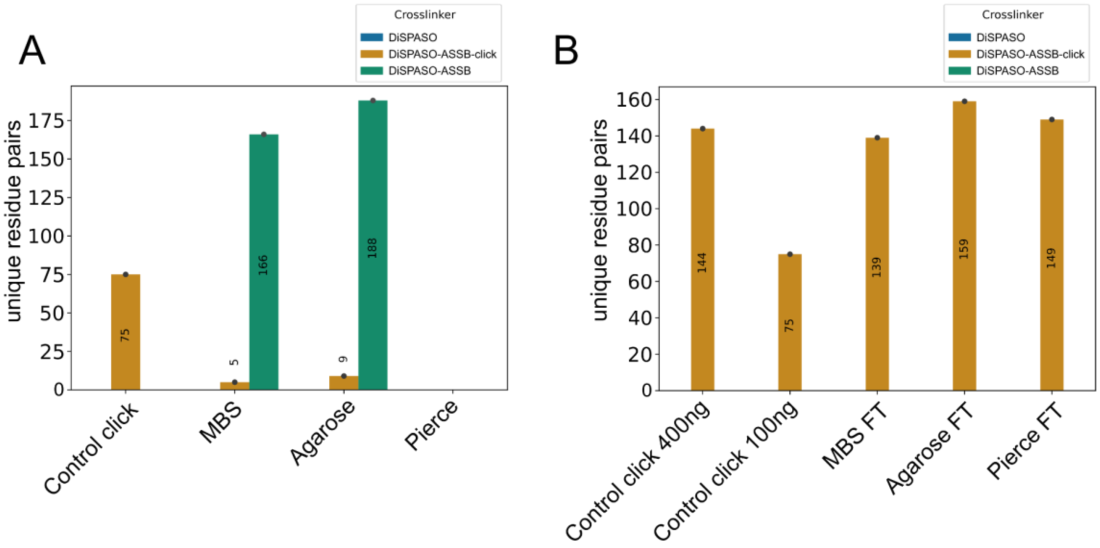

*Figure S4: Comparison of different bead types. A: Different bead types were tested to ensure that the loss of crosslinks after enrichment is not caused by the beads. Three types of beads have been tested, Pierce™ High-Capacity Streptavidin Agarose (Agarose), Pierce™ Streptavidin Magnetic Beads (Pierce) and MBS Magnetic beads-streptavidin (MBS) with Agarose showing the most identified crosslinks after bead enrichment. B: Flowthrough of the experiment showing a high loss of crosslinked peptides after enrichment. The loss of crosslinked peptides might also be independent of the bead type.*

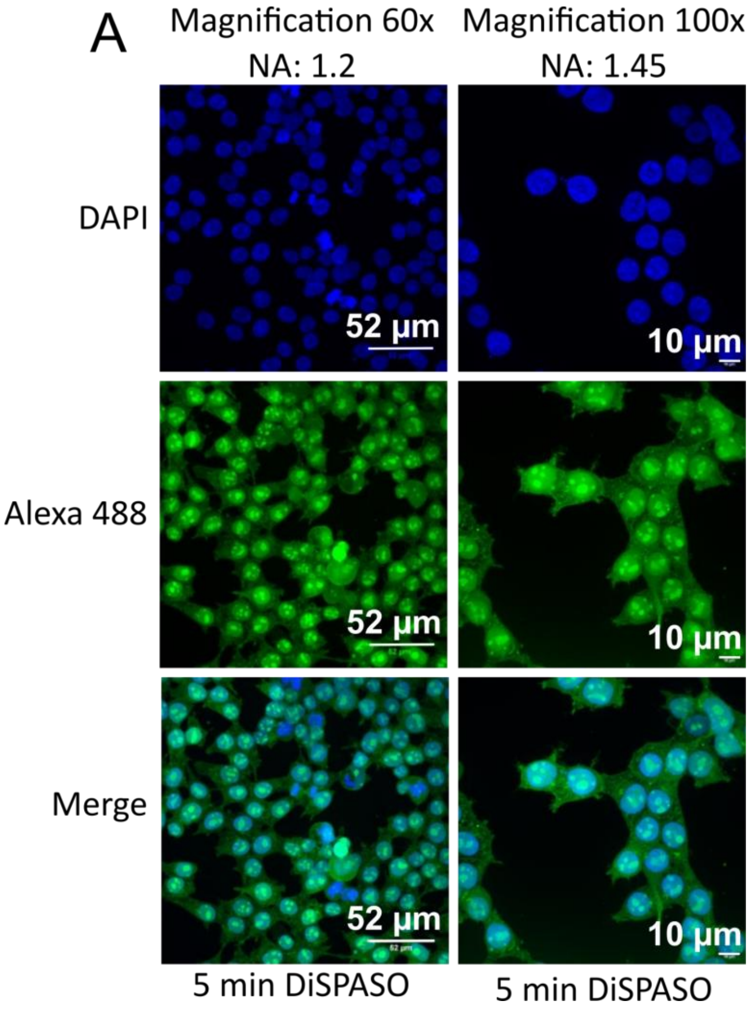

*Figure S5: Confocal microscopy pictures of crosslinked HEK 293 cells using DiSPASO. A: Confocal microscopy images of DiSPASO during in-cell crosslinking experiments with a crosslink duration of 5min. The nuclei fluorescence signal of DAPI is shown in the upper panel in blue, fluorescence of crosslinked peptides after click reaction to Alexa 488 (green) in the middle panel and a merge of both channels on the bottom. The images were taken on an Olympus Spinning Disk Confocal microscope (2-024) using a magnification of either 60 and a numerical aperture of 1.2 (left panel) or a numerical 100 and a numerical aperture of 1.45 (right panel).*

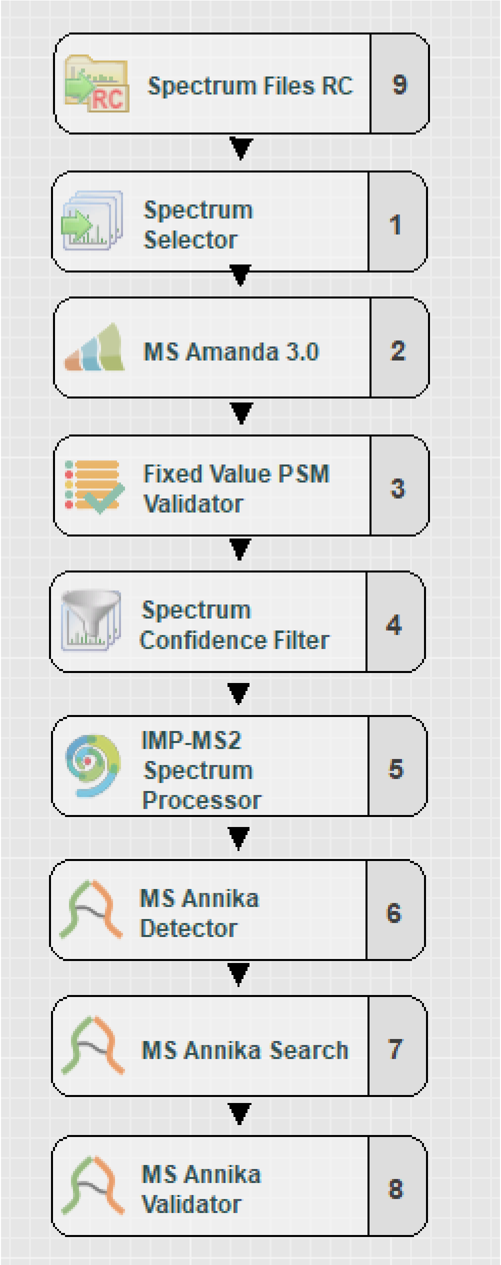

*Figure S6: Exemplary workflow of an MS/MS2 search with MS Annika 2.0 in Proteome Discoverer. The imported raw files are recalibrated by the Spectrum file RC node, followed by a crude selection of suitable spectra for a first search by MS Amanda for linear peptide identification. Spectra that could not map to a linear or monolink peptide are transferred to the MS Annika crosslink search and validation nodes. The crosslink doublet peaks are deisotoped for calculating the peptide's monoisotopic mass, and spectra are adjusted and then searched with MS Annika to identify the cross-linked peptides.*

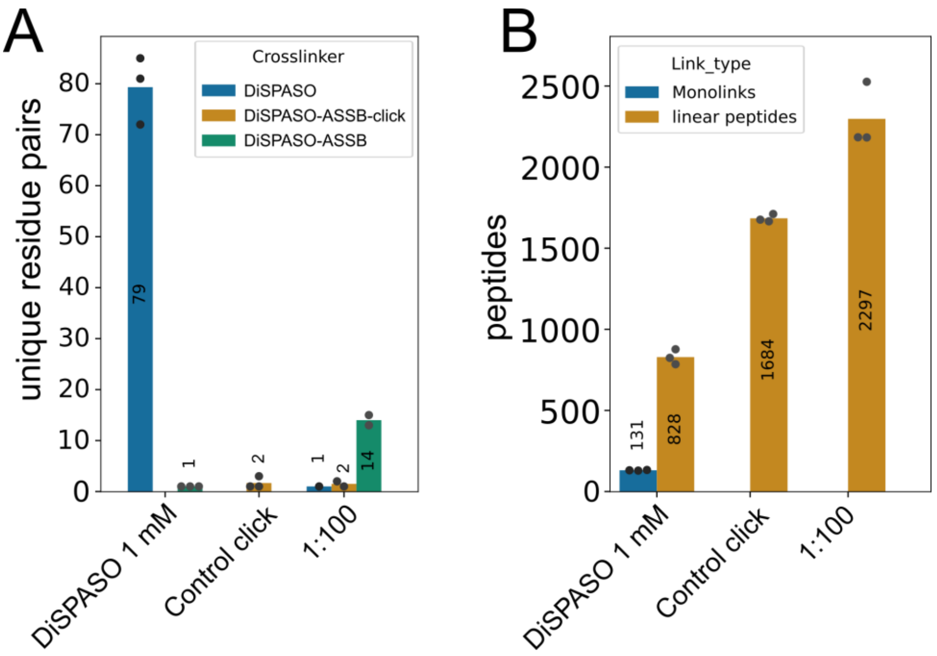

*Figure S7: Application of ASSB-DiSPASO enrichment strategies of spike-in ribosome samples. A: Crosslinked E. coli (K12) ribosome spike-in with HEK 293 cell lysate as background. The ribosome spike-in increased from 0.25ug to 10ug in a constant background of 100ug HEK lysate (only the 1:100 mix is shown here). Part of the 10:100 sample was used as a control (Control-click), showing the performance of ribosome crosslinking without enrichment. The second control sample shows the overall performance of ribosome crosslinking without click reaction or enrichment (blue). The enrichment of 1ug ribosome crosslinked in a 100ug HEK 293 cell lysate background is shown in green. B: Analysis of monolinks and linear peptides of the ribosome spike-in experiment. The monolinks and linear peptides of the "crosslink control" sample show fewer peptides due to ribosome-only related linear peptides that can occur here. The HEK background peptides of the spike-in could be not depleted.*

*Table S2: Fragment names, substitution, and monoisotopic masses of DiSPASO fragments used for crosslinking search.*

| Fragment name | Substitution | Monoisotopic mass [Da] |
|---|---|---|
| DiSPASO | C16H14O4S2 | 334.0333 |
| Alkene (common for all CLs) | C3H2O | 54.01056 |
| Sulfenic acid (common for all CLs) | C3H4O2S | 103.9932 |
| Thiol (common for all CLs) | C3H2OS | 85.98264 |
| ETFP | C13H10O2S2 | 262.0122 |
| ETHMP | C13H12O3S2 | 280.0227 |
| EMP | C13H12O2S | 232.055 |
| **Picolyl** | | |
| DiSPASO-Picoly | C27H30N5O8PS2 | 647.1273 |

| | | |
|---|---|---|
| ETFP | C24H26N5O6PS2 | 575.1062 |
| ETHMP | C24H28N5O7PS2 | 593.1167 |
| EMP | C24H28N5O6PS | 545.1497 |
| **ASSB** | | |
| DiSPASO-ASSB | C20H22N4O5S3 | 494.0752 |
| ETFP | C17H18N4O3S3 | 422.0541 |
| ETHMP | C17H20N4O4S3 | 440.0646 |
| EMP | C17H20N4O3S2 | 392.0976 |
| **ASSB_click** | | |
| DiSPASO-ASSB_click | C30H38N6O6S5 | 738.1456 |
| ETFP | C27H34N6O4S5 | 666.1245 |
| ETHMP | C27H36N6O5S5 | 684.1350 |
| EMP | C27H36N6O4S4 | 636.1680 |
| **DAAB** | | |
| DiSPASO-DAAB | C24H31N5O6S3 | 579.1279972 |
| ETFP | C21H25N5O4S3 | 507.10687 |
| ETHMP | C21H27N5O5S3 | 525.11743 |
| EMP | C21H27N5O4S2 | 477.15045 |

*Table S3: Search parameters for linear and crosslink search. Parameters not listed here were left at default settings*

| Parameter name | Parameter value |
|---|---|
| **Linear search** | |
| MS1 tolerance | 6 ppm |
| MS2 tolerance | 15 ppm |
| Miss cleavages | 3 |
| Fixed modification | Carbamidomethyl [57.021 Da] |
| Variable modification | Oxidation [15.995 Da], DiSPASO amidated [351.059 Da], DiSPASO loop [334.033 Da], DiSPASO hydrolysed [352.043 Da], |

| | |
|---|---|
| | DiSPASO Tris [454.099 Da],<br>DSBSO amidated [325.065 Da], DSBSO loop [308.038 Da], DSBSO hydrolysed [326.049 Da], DSBSO Tris [429.112 Da],<br>Variable modifications for all other crosslinker definitions were calculated accordingly |
| Variable modification for in-cell searches (additional) | Phospho [79.966 Da], Deamidation [0.984 Da], Acetyl Protein N-term [42.011 Da] |
| **Crosslink search** | |
| MS1 tolerance | 6 ppm |
| MS2 tolerance | 15 ppm |
| Miss cleavages | 3 |
| Fixed modification | Carbamidomethyl [57.021 Da] |
| Variable modification | Oxidation [15.995 Da] |
| **IMP-MS2 spectrum processor** | |
| Perform de-isotoping | False |

*Table S4: IUPAC and supplier names of chemical compounds and their abbreviations used in this manuscript.*

| Number of compounds | IUPAC name | Abbreviation |
|---|---|---|
| 1 | bis(2,5-dioxopyrrolidin-1-yl) 3,3'-((5-ethynyl-1,3-phenylene)bis(methylenesulfinyl))dipropanoate | DiSPASO |
| 2 | Bis(1,3-dioxoisoindolin-2-yl) 3,3'-((5-ethynyl-1,3-phenylene)bis(methylenesulfinyl))dipro-pionate | DiPPASO |
| 3 | (4-(6-(azidomethyl)nicotinamido)butyl)phosphonic acid | Picolyl azide |
| 4 | (4-(6-((4-(3,5-bis(((3-((2,5-dioxopyrrolidin-1-yl)oxy)-3-oxopropyl)sulfinyl)methyl)phenyl)-1H-1,2,3-triazol-1-yl)methyl)nicotinamido)butyl)phosphonic acid | BPNB |
| 5 | N-(2-((2-azidoethyl)disulfanyl)ethyl)-5-(2-oxohexahydro-1H-thieno[3,4-d]imidazol-4-yl)pentanamide | ASSB |
| 6 | 2-((2-(4-(3,5-bis(((3-(2,5-dioxopyrrolidin-1-yl)-3-oxopropyl)sulfinyl)methyl)phenyl)-1H-1,2,3-triazol-1-yl)ethyl)thio)acetamide | BAED |
| 7 | Disulfide azide beads (supplier name) | DAAB |
| 8 | 1,1'-(3,3'-((5-(1-(3-aminopropyl)-1H-1,2,3-triazol-4-yl)-1,3- | BAPD |

| | | |
|---|---|---|
| | phenylene)bis(methylenesulfinyl))bis(propanoyl))bis(pyrrolidine-2,5-dione) | |
| 9 | 3-((3-ethynyl-5-thioformylbenzyl)sulfinyl)propanal | ETFP |
| 10 | 3-((3-ethynyl-5-((hydroxythio)methyl)benzyl)sulfinyl)propanal | ETHMP |
| 11 | Acrylaldehyde (Alkene) | A |
| 12 | 3-((3-ethynyl-5-methylbenzyl)sulfinyl)propanal | EMP |
| 13 | 3-(hydroxythio)propanal (Sulfenic acid) | SA |
| 14 | (E)-3-mercaptoacrylaldehyde (Thiol) | T |

#### References

1. Fulmer, G. R. *et al.* NMR Chemical Shifts of Trace Impurities: Common Laboratory Solvents, Organics, and Gases in Deuterated Solvents Relevant to the Organometallic Chemist. (2010) doi:10.1021/om100106e.
2. Mazik, M. & König, A. Recognition properties of an acyclic biphenyl-based receptor toward carbohydrates. *J. Org. Chem.* **71**, 7854–7857 (2006).
3. Chen, Z. *et al.* A New Multidentate Hexacarboxylic Acid for the Construction of Porous Metal−Organic Frameworks of Diverse Structures and Porosities. (2010) doi:10.1021/cg100316s.
4. Crosignani, S. *et al.* Discovery of Potent, Selective, and Orally Bioavailable Alkynylphenoxyacetic Acid CRTH2 (DP2) Receptor Antagonists for the Treatment of Allergic Inflammatory Diseases. (2011) doi:10.1021/jm200866y.
5. Marcum, J. S., Taylor, T. R. & Meek, S. J. Enantioselective Synthesis of Functionalized Arenes by Nickel-Catalyzed Site-Selective Hydroarylation of 1,3-Dienes with Aryl Boronates. *Angew. Chem. Int. Ed Engl.* **59**, 14070–14075 (2020).
6. Deng, W., Shi, X., Tjian, R., Lionnet, T. & Singer, R. H. CASFISH: CRISPR/Cas9-mediated in situ labeling of genomic loci in fixed cells. *Proc. Natl. Acad. Sci. U. S. A.* **112**,

11870–11875 (2015).

7. Rappsilber, J., Mann, M. & Ishihama, Y. Protocol for micro-purification, enrichment, pre-fractionation and storage of peptides for proteomics using StageTips. *Nat. Protoc.* **2**, 1896–1906 (2007).
8. Ishihama, Y., Rappsilber, J. & Mann, M. Modular stop and go extraction tips with stacked disks for parallel and multidimensional peptide fractionation in proteomics. *J. Proteome Res.* **5**, 988–994 (2006).
9. Müller, F., Graziadei, A. & Rappsilber, J. Quantitative photo-crosslinking mass spectrometry reveals protein structure response to environmental changes. *Anal. Chem.* (2019) doi:10.1021/acs.analchem.9b01339.
10. Dorfer, V. *et al.* MS Amanda, a Universal Identification Algorithm Optimized for High Accuracy Tandem Mass Spectra. (2014) doi:10.1021/pr500202e.
11. Pirklbauer, G. J. *et al.* MS Annika: A New Cross-Linking Search Engine. *J. Proteome Res.* (2021) doi:10.1021/acs.jproteome.0c01000.
12. Birklbauer, M. J., Matzinger, M., Müller, F., Mechtler, K. & Dorfer, V. MS Annika 2.0 Identifies Cross-Linked Peptides in MS2–MS3-Based Workflows at High Sensitivity and Specificity. *J. Proteome Res.* (2023) doi:10.1021/acs.jproteome.3c00325.
13. Nelli, F. *Pandas in 7 Days: Utilize Python to Manipulate Data, Conduct Scientific Computing, Time Series Analysis, and Exploratory Data Analysis (English Edition)*. (BPB Publications, 2022).
14. Harris, C. R. *et al.* Array programming with NumPy. *Nature* **585**, 357–362 (2020).
15. Hunter, J. D. Matplotlib: A 2D Graphics Environment. *Comput. Sci. Eng.* **9**, 90–95 (2007).
16. Waskom, M. seaborn: statistical data visualization. *J. Open Source Softw.* **6**, 3021 (2021).
17. Garreta, R. & Moncecchi, G. *Learning Scikit-Learn: Machine Learning in Python*. (Packt Pub Limited, 2013).
18. Virtanen, P. *et al.* SciPy 1.0: fundamental algorithms for scientific computing in Python. *Nat. Methods* **17**, 261–272 (2020).

840
